# Liquidity of gene co-expression trajectories across the lifespan highlights delayed maturation and the perinatal GABA switch in schizophrenia risk

**DOI:** 10.64898/2026.06.10.731457

**Authors:** Loredana Bellantuono, Fabio Di Camillo, Christopher Borcuk, Gianluca C. Kikidis, Joel E. Kleinman, Madhur Parihar, Thomas M. Hyde, Daniel R. Weinberger, Giulio Pergola

**Affiliations:** Department of Translational Biomedicine and Neuroscience, University of Bari Aldo Moro, Bari, Italy; Istituto Nazionale di Fisica Nucleare, Bari, Italy; Lieber Institute for Brain Development, Johns Hopkins Medical Campus, Baltimore, MD, USA; Department of Psychiatry and Behavioral Sciences, Johns Hopkins University School of Medicine, Baltimore, MD, USA; Department of Neurology, Johns Hopkins University School of Medicine, Baltimore, MD, USA; Department of Neuroscience, Johns Hopkins University School of Medicine, Baltimore, MD, USA; Department of Genetic Medicine, Johns Hopkins University School of Medicine, Baltimore, MD, USA

**Keywords:** Lifespan transcriptomic trajectories, genetic risk, schizophrenia, gene co-expression networks, E/I imbalance

## Abstract

Schizophrenia genetic and environmental risk factors play out largely in early life, biasing development toward a pathogenic trajectory that becomes clinically apparent in early adulthood, when the disorder typically onsets. Here, we convert snapshots of gene expression in postmortem brain at a moment in time into a dynamic lifetime series employing the “liquidity” metric, a novel tool to track the evolution of networks across time from a multi-systemic perspective. The landscape of normal prefrontal cortical development becomes increasingly “liquid” during the first two decades of post-natal life, with sharp discontinuities in known critical periods such as birth and adolescence. Neurotypical individuals free of apparent neuropathology with relatively elevated polygenic risk scores for schizophrenia exhibit a generalized delay in the dynamics of liquidity across these trajectories compared to below-average-risk individuals. Impacted biological processes strongly converge on delayed GABA-A receptor functional maturation, involved in establishing Excitatory/Inhibitory balance in brain during early development. Similar to patients with schizophrenia, neurotypical high-risk individuals show an increased expression ratio between the genes *SLC12A2* (protein NKCC1) and *SLC12A5* (protein KCC2) relative to low-risk, involved in the control of the equilibrium potential of chloride ions that regulates GABA-A function. These results provide evidence that genetic risk for schizophrenia is associated with a delayed maturational profile and delayed maturation of GABAergic signaling without detectable neuropathology and well before the age of clinical onset. Interestingly, the same effect is not observed in the hippocampus and is not observed with genetic risk for other neuropsychiatric and immune conditions.

The dynamics of maturation of GABA-A signaling in the dorsolateral prefrontal cortex emerge as an early contributor to translating genetic risk into an altered developmental trajectory associated with schizophrenia.

## Introduction

Studies of gene expression in postmortem human brain have opened a window into psychiatric neuroscience. Many efforts have focused on schizophrenia (SCZ) because of its relevant societal cost, high heritability, extensive catalog of identified risk-associated genes, and unmet clinical needs (*1, 2*). Despite its clinical onset in young adulthood, SCZ risk factors are thought to play out in early development, even in utero (*3–7*). Genome-Wide Association Studies (GWAS) have revealed many variants statistically associated with SCZ (*8*). These variants colocalize with genes involved in synaptic biology, but narrowing down more specific biological mechanisms has been challenging because genetic risk does not apparently converge onto specific processes, with many genes potentially involved (*9*). However, prior research has shown that risk genes converge into co-expression networks (*10–18*). Most gene co-expression studies have pooled together subjects of different ages, although the neurodevelopmental origin of SCZ demands capturing the evolution of gene co-expression and of a risk trajectory across multiple age periods (*7, 19, 20*). Overcoming these limitations requires dynamic methods to account for network changes across the lifespan.

As one example, Pergola et al. (*21*) analyzed the variation of SCZ risk gene convergence into co-expression networks over time. Putative SCZ risk-associated genes clustered in the dorsolateral prefrontal cortex (DLPFC) into sets (called “modules”) at perinatal age (fetal life to 5 years old) and kept being co-expressed at juvenile age (5 to 25 years old) above chance level, while their co-expression partners changed in the transition from perinatal to juvenile age. Borcuk et al. (*9*) also suggested that risk genes for SCZ identified via GWAS are co-expressed with different genes during perinatal and postnatal life. These findings suggested that co-expression partner genes across time represent a shifting molecular environment for risk genes necessary to translate early risk into biological risk and manifest illness by adulthood.

Previous approaches, however, were limited in several respects. First, the module membership criterion used in most previous work is a binary (yes/no) criterion of co-expression that misses the much more nuanced representation of co-expression available in weighted networks. Second, the definition of disjoint time windows, while capturing potentially meaningful developmental stages, is not granular enough to render trajectories of gradual changes in molecular readouts. A sliding age-window approach is more effective to this end (*21*). Previous work also left unanswered how overall genomic risk relates to time-dependent shifts in the molecular environment at an individual subject level, despite reports that individual polygenic risk for schizophrenia is associated with co-expression network features (*11, 18, 22, 23*).

Here, we aimed at identifying genes potentially involved in the physiological mechanisms underlying genetic risk trajectories in two independent samples of neurotypical subjects. To evaluate risk convergence for SCZ in networks across the lifespan, we quantified the tendency of genes to vary their co-expression relations over time through the metric ℒ_*t*1,*t*2_ (*g*), known as *liquidity* (*24*). This measure represents the volatility of the links with gene *g* in the comparison of the two co-expression networks referred to times *t*_1_and *t*_2_(see ***Materials and Methods***). Besides overcoming the limitations entailed by using modules in co-expression networks, liquidity models co-expression changes across the whole lifespan, and can be evaluated at the level of single genes. We studied liquidity in a cohort of subjects free of apparent neuropathology (“neurotypical”) to validate its effectiveness as a probe of gene co-expression variability, and then in groups with relatively high and low Polygenic Risk Score (PRS) for SCZ to understand how gene-gene co-expression relationships change as a function of overall genomic risk throughout life.

Specifically, we grouped samples from neurotypical individuals into chronologically sequential, partially overlapping sliding age windows. For each window, we constructed the corresponding co-expression network and computed gene liquidity values by comparing it with the co-expression network of the first window in the sequence. We then repeated this procedure for two distinct cohorts of neurotypical subjects with relatively high and low PRS, carefully matched for confounding variables, and compared the respective liquidity distributions. We aimed to extract potential candidates from the pool of examined genes to explore the physiological mechanisms underlying the trajectory of risk. The research workflow is summarized in the graphical abstract in Figure 1.

**Figure 1.**
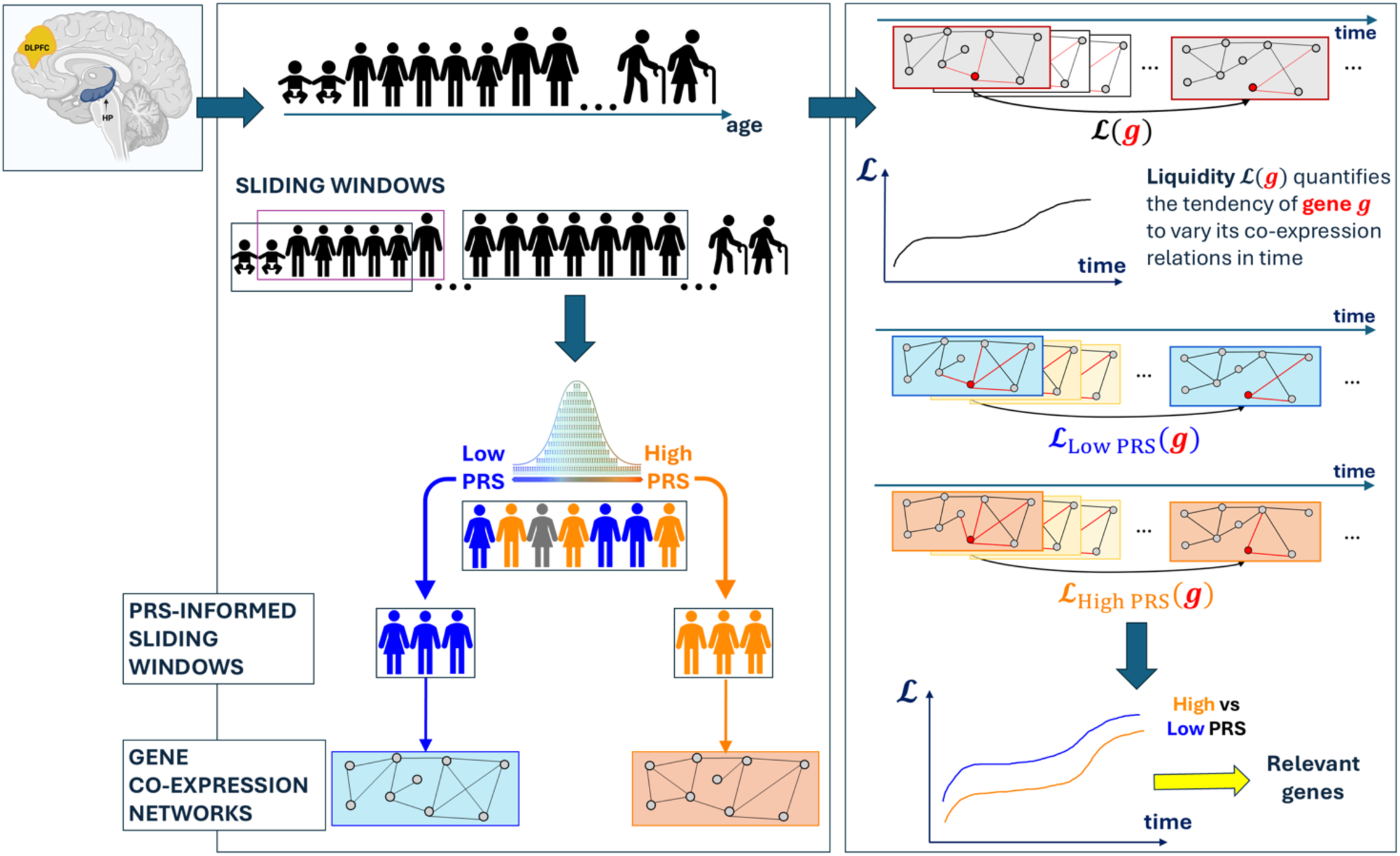
Graphical abstract of the study design. We introduced the liquidity framework, measuring time variability of gene co-expression patterns, and tested its effectiveness as a probe to identify, out of many genes, relevant candidates involved in pathways potentially explaining the heterogeneity of risk associated trajectories in neurotypical individuals. Liquidity distributions referred to high and low-PRS subjects were compared, considering co-expression networks obtained from chronologically sequential sliding windows. DLPFC, dorsolateral prefrontal cortex; HP, hippocampus; PRS, polygenic risk score. The brain picture was created with <u>BioRender.com</u>.

## Results

The findings presented hereafter are based on a dataset including 541 postmortem brain samples from neurotypical individuals of European and African American ancestry, on which tissue homogenate RNA-sequencing was performed via the Illumina Ribo-Zero Kit (*21*). We replicated specific results in high- and low-risk cohorts of neurotypical controls using a non-overlapping dataset consisting of 91 brain samples RNA sequenced with the Poly(A) sequencing (*25*). More detailed demographic information on samples included in both Ribo-Zero and Poly(A) datasets is reported in Supplementary Table S1.

### Sliding window analysis

For each brain region surveyed (DLPFC and HP), we grouped subjects by age in ascending order into sliding windows of 40 samples each, with each window being shifted by a single sample with respect to the previous one (*21*). We computed the liquidity metric for all genes across these windows, taking the initial window as a reference, to quantify the tendency of each gene to vary its co-expression partners with changes in age.

The plots in Figure 2(a) indicate that liquidity generally increases throughout the lifespan (as expected, since it is referenced to the initial window) and, remarkably, that it exhibits a steeper rise in crucial phases of neurodevelopment, i.e., birth and adolescence. To provide a quantitative assessment of these rapid growth periods, we analyzed the time derivative of liquidity, which exhibits several distinct peaks corresponding to the maximum rates of change. Specifically, for DLPFC, we identified peaks in the sliding windows with median ages of -0.007, 0.21, 0.30 (birth to the first four months of life), and 15 years (adolescence). For HP, the derivative highlighted sharp liquidity increases at median ages of 0.30, 15, and 37 years. These results suggest that this liquidity measure captures biological insight into stages of neurodevelopment.

**Figure 2.**
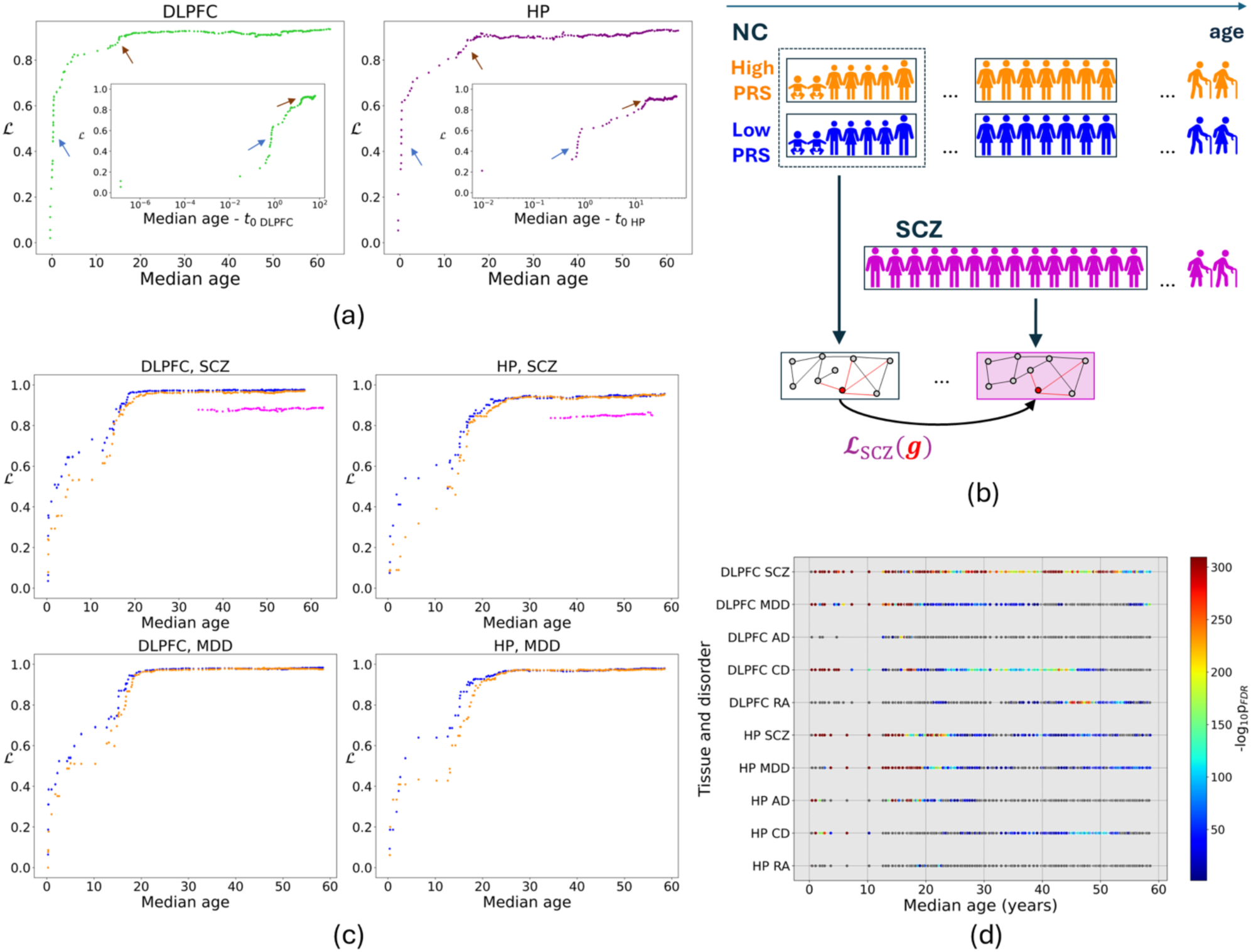
Liquidity of genes in sliding age windows and comparison between high- and low-PRS liquidity across tissue-disorder combinations. Panel (a) shows the liquidity obtained by comparing the co-expression network in each 40-subject sliding age window to the earliest one, taken as a reference; the liquidity evolution profile is reported as a function of the median age of subjects within each sliding window (main frames). The same quantity is reproduced in the insets, where the median age of each window is measured with respect to that of the earliest sliding window for the brain tissue of interest (*t*_0 *DLPFC*_ = *t*_0 *HP*_ = −0.4027 years) and expressed in logarithmic scale. This visualization emphasizes the steeper liquidity increases at birth (blue arrows) and during adolescence (brown arrows). Panel (b) represents the approach used to calculate the liquidity of the co-expression networks of SCZ patients, relative to a reference network constructed from an integrated cohort of the initial high- and low-PRS NC subjects. Panel (c) shows the liquidity profiles in the co-expression networks obtained from high-PRS (orange) and low-PRS (blue) subjects for the considered polygenic disorders of a psychiatric nature; only in the plots referred to SCZ, liquidity profiles for patients (magenta) are reported; dots indicate the median of the liquidity distributions, and error bars represent the 95% confidence interval of the median, as determined from 10000 bootstrap resamples. Panel (d) illustrates the comparison between the liquidity distributions of all genes in co-expression networks associated with high- and low-PRS subjects for different combinations of tissue and disorder, in terms of the p-value *p*_*FDR*_ (with Benjamini-Hochberg correction for multiple comparisons in all sliding windows) of a Wilcoxon signed-rank test with the alternative hypothesis that the liquidity distribution referred to high-PRS subjects is lower than that of the low-PRS cohort; sliding windows for which the difference between the high- and low-PRS liquidity distributions is not significant are represented as grey dots. Results reported in panels (c) and (d) are expressed as a function of the median age of the 60-subject sliding windows from which the high-PRS and low-PRS cohorts have been extracted. DLPFC, dorsolateral prefrontal cortex; HP, hippocampus; PRS, polygenic risk score; SCZ, schizophrenia; MDD, major depressive disorder; AD, Alzheimer’s disease; CD, Crohn’s disease; RA, rheumatoid arthritis.

### PRS-informed sliding windows

For both DLPFC and HP, we compared the liquidity distributions in co-expression networks in subjects referred to as relatively high- and relatively low-PRS for SCZ, major depressive disorder (MDD; chosen as a psychiatric disorder outside the psychosis spectrum), Alzheimer’s disease (AD; chosen as a neurodegenerative disorder), Crohn’s disease, and rheumatoid arthritis (respectively, CD and RA; chosen as non-brain disorders). We constructed a new series of sliding windows, this time consisting of 60 subjects each, to account for reduced statistical power following the dataset partition through which we divided subjects in each sliding window in high-and low-PRS cohorts. We explicitly controlled for subject attributes (first, second, and third genomic eigenvariates (*26*) to model ancestry stratification, age, sex, RNA integrity number, and postmortem interval) and sequencing variables (including number of reads successfully mapped to the reference genome during alignment, fraction of reads mapped to the mitochondrial chromosome, fraction of reads assigned unambiguously to a gene, ribosomial RNA, and individual neuronal proportion). To further mitigate ancestry confounds, besides matching genomic eigenvariates, we standardized PRSs within-ancestry prior to subject selection. A complete list of these confounders and further information on the workflow to define high-PRS and low-PRS subsets are provided in Materials and Methods.

For each sliding window, we constructed the high- and low-PRS co-expression networks, using WCGNA (*21*) from the corresponding cohorts of risk-stratified subjects. We computed the liquidity metric for all genes across the time series of high- and low-PRS co-expression networks, taking the respective initial networks as references, to quantify the tendency of each gene to change its co-expression partners with age. Then, we constructed gene co-expression networks from SCZ patients, ordered by ascending age and grouped into sliding windows, and computed their liquidity relative to a reference network, obtained by merging the earliest high- and low-PRS windows of NC subjects (see Materials and Methods). This procedure, motivated by the absence of SCZ patients in the perinatal period, is outlined in Figure 2(b). Figure 2(c) reports the liquidity profiles in the co-expression networks obtained from high- and low-PRS subjects, for the considered disorders of psychiatric interest; Supplementary Figure S1 shows the analogous outcomes for AD, CD, and RA. The plots for DLPFC samples stratified by risk for SCZ highlight a decreasing hierarchy in gene co-expression dynamics from low-PRS NC to patients: liquidity is highest in networks of low-risk individuals and progressively declines through high-risk subjects and SCZ patients. The same liquidity hierarchy between high- and low-PRS cohorts for SCZ has been obtained using the Poly(A) replication dataset of DLPFC samples (see Supplementary Figure S2). Figure 2(d) highlights, across all tissue-disorder combinations, the sliding windows in which the liquidity distribution of all genes in co-expression networks for high-PRS NC subjects is significantly lower than that of their low-PRS counterparts (analogous plots displaying significant differences of the opposite sign are provided in Supplementary Figure S3). While the DLPFC-SCZ, DLPFC-MDD, DLPFC-CD, HP-SCZ, and HP-MDD cases consistently exhibit high-risk co-expression patterns that are significantly less liquid than low-risk patterns across nearly the entire age range, the statistical significance of these differences is markedly more pronounced in the DLPFC-SCZ case (Figure 2(d)).

As shown in Supplementary Figures S4-S5, the effects highlighted by the liquidity data did not emerge when merely considering the mean gene expression distributions in the PRS-informed sliding windows, suggesting that this insight is not accessible via gene expression alone, but readily emerges as a network dynamics property.

### Functional genetics of SCZ-PRS liquidity differences

An essential advantage of liquidity analysis lies in quantifying the volatility of co-expression patterns at the single-gene level, considering all the connections with other genes in the network, each with its own weight, regardless of module membership. For each gene, we quantified the significance of differences between the gene-wise liquidity distributions related to high- and low-PRS cohorts across the lifespan. The obtained significance ranking was then used as an input to perform gene set enrichment analyses (GSEA). The outlined procedure is schematically illustrated in the upper part of Figure 3(a) and described in detail in Materials and Methods. Specifically in the context of DLPFC-SCZ, we found consistent results across three distinct tools (PANTHER (*27*); gProfiler (*28*); STRING (*29*)), pointing to GABA neurotransmission, especially the GABA-A receptor (Supplementary File S1). To assess the robustness and temporal specificity of these findings, we used alternative time windows subsequent to the first as references for the liquidity calculation. As our chronological series started on the second trimester of pregnancy, we considered as alternative starting points the first 6 windows, all beginning with subjects within the second trimester of pregnancy and all featuring a median age < 1 year old. We found that the GSEA outcomes concerning the GABA receptor complex persisted when liquidity was calculated relative to the second and third chronological windows, mitigating the concern that results are driven by the first subject in the first window.

**Figure 3.**
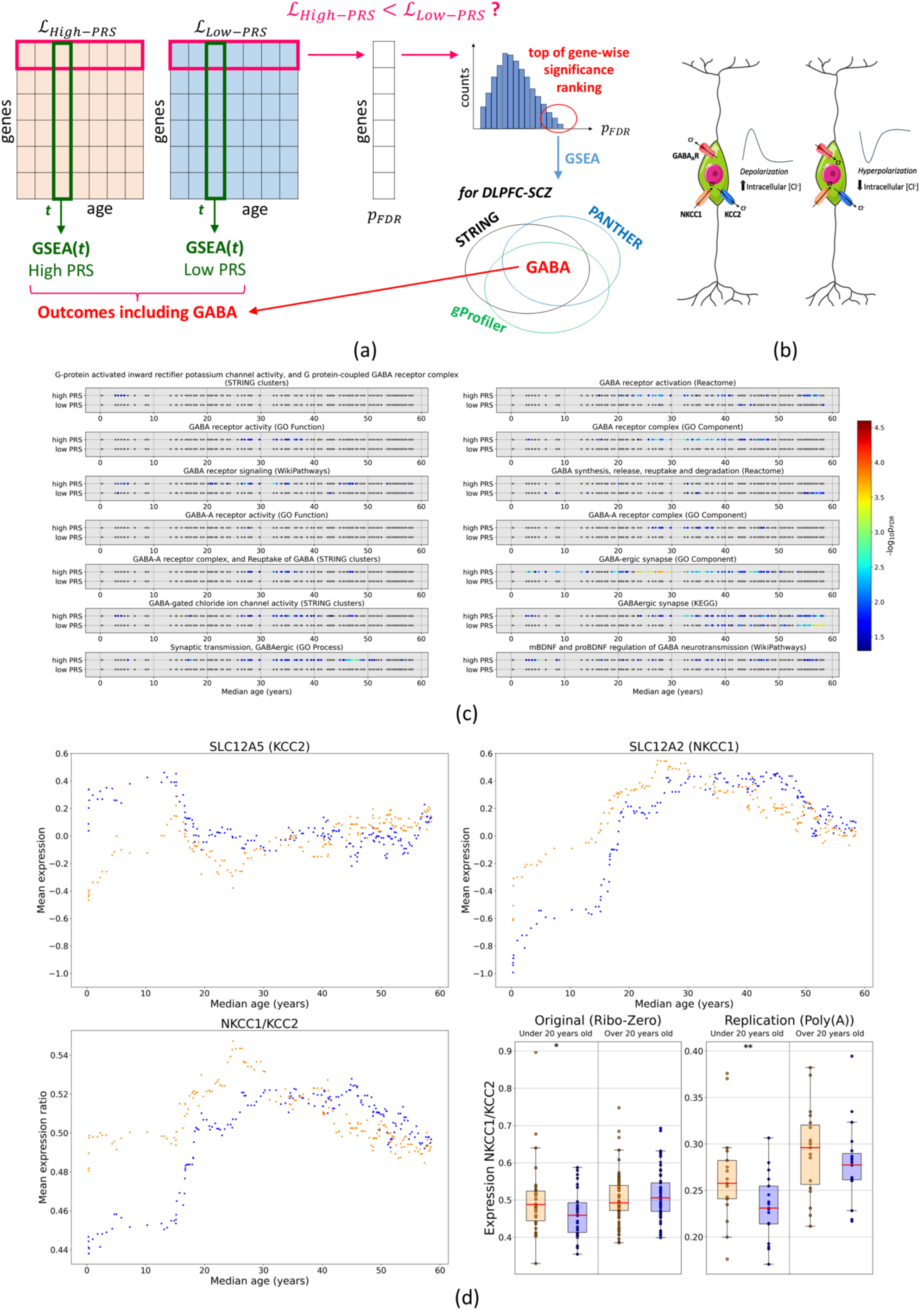
Comparing high- and low-PRS liquidity of genes in the DLPFC highlights the role of the GABA-switch in genetic risk for SCZ. Panel (a) shows a graphical representation of the GSEA procedures. The upper part illustrates a scheme of the enrichment from the significance ranking of the gene-wise difference between high- and low-PRS liquidity for SCZ; in the case of DLPFC-SCZ, the analysis unveils a GABAergic pathway. The lower part depicts the GSEA taking as inputs the liquidity scores at fixed sliding windows, processed separately for high- and low-PRS groups; the outcomes of this procedure, obtained via the STRING software, were subsequently filtered to include only those containing the term “GABA” in their description. Panel (b) illustrates the GABA action switch from excitatory to inhibitory during neurodevelopment (31). Panel (c) reports the GABA-related findings returned by the GSEA based on high- and low-PRS liquidity scores at fixed sliding windows. For each outcome, the corresponding STRING category is indicated in parentheses. The panel displays the enrichment significance for these results across all sliding windows, with non-significant values represented in gray. Notably, all significant instances represent cases where the genes in the enrichment are significantly less volatile than the remaining gene universe. Panel (d) shows information on the expression of the genes SLC12A2 (NKCC1) and SLC12A5 (KCC2) in high- and low-PRS subjects for SCZ; the top plots report standardized residuals of expression, while the bottom left plot shows the ratio of the original expression values. The bottom right plot shows the high- and low-PRS distributions of NKCC1/KCC2 expression values for subjects grouped by age (under and over 20 years old) in both the original (Ribo-Zero) and replication (Poly(A)) datasets, highlighting significant differences (one-way Wilcoxon rank-sum test) between high- and low-PRS distributions referred to the same age group and dataset. Results reported in panels (c) and (d) are expressed as a function of the median age of the 60-subject sliding windows from which the high-PRS and low-PRS cohorts have been extracted. Panel (b) created using Servier Medical Art Commons Attribution 3.0 Unported License (http://smart.servier.com). Servier Medical Art by Servier is licensed under a Creative Commons Attribution 3.0 Unported License. PRS, polygenic risk score; GSEA, gene set enrichment analysis; DLPFC, dorsolateral prefrontal cortex; SCZ, schizophrenia.

The relatively lower liquidity in neurotypicals with high risk for SCZ points to developmental delays in high-risk compared to low-risk individuals, an effect that was not significant in time windows centered on adult ages. Additionally, in the *regulation of postsynaptic membrane potential* ontology (biological process), GSEA identified the involvement of glutamate receptor genes, along with genes related to GABA, thus forwarding the development of the excitatory-inhibitory (E/I) imbalance as a potential mechanism of risk. The molecular and biological implications of this finding are discussed in the Supplementary Material (**Focus on glutamate genes returned by GSEA**).

Furthermore, when replicating these analyses on samples from DLPFC and HP grouped based on high- and low-PRS for other disorders (Supplementary Files S4-S12), results indicated that the emergence of GABA-A genes from GSEA was specific to the DLPFC-SCZ case. When merely comparing the distributions of mean gene expression on all the high and low-PRS groups (top left panel of Supplementary Figure S4), the significance ranking did not yield any outcome related to the GABA-A genes. Therefore, this insight is evident only when analyzing the dynamic evolution of co-expression networks through the lifespan.

Considering the potential role of the GABA-A excitatory to inhibitory function switch in SCZ (*30*), we further investigated the expression of the critical genes returned by the STRING, PANTHER and gProfiler GSEA on liquidity within the gene ontologies *GABA-A receptor activity* (molecular function) and *GABA-A receptor complex* (cellular component). Interestingly, while most of these GABA-A genes did show significant differences between the high- and low-PRS cohorts, these differences had mixed directions (see Supplementary Table S2), unlike liquidity. Given that connection patterns in the high-PRS group changing significantly less than those in the low-PRS group, we next investigated the precise drivers of this divergence, aiming to disentangle whether the seeming delay reflected lower liquidity in high-PRS networks or higher liquidity in low-PRS networks. To this aim, we implemented an additional GSEA procedure in STRING, using the high- and low-PRS liquidity scores of the DLPFC genes at fixed sliding windows as inputs (Figure 3(a), lower part). For each PRS group and each sliding window, we filtered the GSEA results for descriptions containing the term “GABA”, obtaining the outcomes reported in Figure 3(c). We found that all of the detected liquidity differences reflected GABA-specific drops in liquidity relative to the remaining gene universe. Liquidity drops were more common and more significant, although not exclusive, of the high-PRS group, suggesting generally slower GABA-related co-expression dynamics in these subjects.

### Focus on GABA signaling-related genes NKCC1 and KCC2

Among other processes involved, GABA-A function depends on chloride resting potential, which changes during early neurodevelopment (*32–34*). Furthermore, the enrichment of GABA-gated chloride channels directly relates to the developmental shift in chloride gradients, which is driven by the changing balance between NKCC1 and KCC2 transporters. Additionally, GSEA results in Figure 3(c) regarding BDNF regulation are complemented by the finding that 26 of the 33 BDNF genes, including *SLC12A5*, are significantly less liquid in the high-PRS than in the low-PRS network. Together, these results are aligned with the role of BDNF in upregulating KCC2 (coded by *SLC12A5*) expression, thereby orchestrating the transition of GABAergic signaling from excitatory to inhibitory (*33*).

In SCZ, Hyde et al. (*30*) observed an increased NKCC1/KCC2 ratio (with NKCC1 coded by gene *SLC12A2*) in postmortem brain from patients. We thus hypothesized that these differences between patients and controls would be reflected in high- vs. low-risk individuals without overt neuropathology. Figure 3(d) (top and bottom left panels) shows the mean expression values in terms of the standardized residuals alongside the ratio of the original expression values of the genes mentioned above in the high- and low-PRS cohorts for SCZ. Here, we observed that high-risk subjects had higher median ratio values than low-risk subjects at younger ages. The gap in the NKCC1/KCC2 ratio between the high- and low-risk networks decreased in sliding windows with a median age of around 20. To investigate the impact of PRS on the NKCC1/KCC2 gene expression dynamics across different life stages, we grouped subjects by age using a sliding threshold within the juvenile range (5–25 years) (*21*). Our analysis revealed that high-PRS individuals exhibited a significantly higher NKCC1/KCC2 gene expression ratio during this developmental window compared to the low-PRS group. These findings were robust across various age upper cut-offs within the juvenile range, as detailed in Supplementary Table S3, and were further validated in an independent replication set of samples sequenced using Poly(A) RNA sequencing. In contrast, we found no significant genetic risk effect on the NKCC1/KCC2 gene expression ratio in subjects older than any juvenile age threshold in either the discovery or replication datasets. As illustrated in the bottom right panel of Figure 3(d) (representative case with age threshold at 20 years), these results imply that risk-dependent differences in the NKCC1/KCC2 gene expression ratio are specific to the juvenile phase.

## Discussion

This study aimed to analyze the developmental dynamics of gene co-expression in the brain and its relationship with polygenic risk for SCZ and negative control conditions. We used “liquidity” as an analytical tool to extract unique information on brain development in neurotypical subjects that is not accessible through mere gene expression analyses. Liquidity of gene co-expression relationships increased most dramatically during the first two decades of life. With liquidity of gene co-expression thus consistent with this developmental biology, we explored how it played out in the context of polygenic risk for SCZ. Compared to subjects at relatively low risk, subjects with relatively higher risk showed a generalized delay in the dynamics of co-expression relations during neurodevelopment, consistent with previous observations in patients relative to neurotypical controls (30). Using liquidity as a searchlight, we revealed a specific mechanism previously observed only in case-control comparisons, here associated with polygenic risk within neurotypical subjects. Results emphasized the role of genes involved in GABA-A receptor maturation, particularly genes that regulate the function of the channel, as a potential mechanism involved in this apparent delay.

The inferred delay of neurodevelopmental co-expression patterns in high-risk neurotypical subjects echoes the abundant evidence of neurodevelopmental deviations observed in the anamnesis of patients with SCZ (*20, 35, 36*) and relevant to subsequent diagnosis (*37*). GABA (γ-aminobutyric acid) is the primary inhibitory neurotransmitter in the central nervous system. The associated Cl- influx in mature neurons (given the low intracellular Cl- concentration) results in hyperpolarization (*38*). However, in immature neurons, GABA binding causes Cl-eflux (due to a high intracellular Cl-concentration), leading to depolarization (*39–45*). The GABA-switch, transitioning from depolarization to hyperpolarization typically occurring in early postnatal life, is critical for widespread brain development (*46, 47*), regulating initial network functions, neuronal migration, circuit establishment, and synapse maturation during neurodevelopment (*48–54*), and has already been investigated as a potential target for therapeutic drug development (*55*). It is worth noting that GABAergic signaling in immature neurons is not unidirectionally excitatory (*56–58*), and that the regulation of chloride resting state potential mediated by the NKCC1/KCC2 ratio exerts multiple effects on other systems, including glutamate. Therefore, we report results on these genes as a case study offered by the liquidity framework, and we interpret results as implicating the patterning of E/I balance in the developing cortex. Notably, KCC2 promotes dendritic spine maturation and synapse formation independently of Cl⁻ transport, thereby coordinating the assembly of excitatory and inhibitory circuits during early postnatal development (*46*).

It is interesting to note that our enrichment result highlighted the “mBDNF and proBDNF regulation of GABA neurotransmission” pathway, which also was significantly less liquid in DLPFC high-PRS SCZ networks (Figure 3(c)). Indeed, BDNF-related signaling has been implicated in the developmental regulation of the GABA switch, primarily through effects on chloride homeostasis and KCC2 expression (*30, 59, 60*). More specifically, mature BDNF and proBDNF appear to exert dissociable effects on GABAergic maturation: BDNF/TrkB signaling has been linked to the promotion of inhibitory maturation and increased KCC2 expression in developmental contexts, whereas proBDNF/p75NTR signaling has been associated with reduced GABAergic efficacy, including GABA_A_ receptor internalization and maintenance of immature chloride handling (*61, 62*). Recent experimental evidence further shows that both proBDNF and mBDNF can modulate chloride extrusion in immature neurons, with proBDNF promoting KCC2 endocytosis and prolonging a depolarizing *E*_*GABA*_state (*63*). Although the precise developmental effects of BDNF-related signaling on NKCC1 and KCC2 remain context- and stage-dependent, these observations support the interpretation that altered stabilization of this pathway may converge on delayed maturation of chloride-dependent GABAergic signaling, thereby providing a mechanistic rationale for examining the NKCC1/KCC2 ratio in high-PRS subjects.

In patients with SCZ, an elevated NKCC1/KCC2 ratio has been observed, suggesting relatively immature GABA physiology (*30, 64, 65*). Moreover, SCZ susceptibility genes, including *CHRNA7*, *GAD1*, and *BDNF*, have been observed to exert regulatory control over KCC2 expression (*66, 67*). Notably, our data show that altered NKCC1 and KCC2 expression patterns emerge in neurotypical individuals at increased genetic risk for SCZ. Interestingly, the difference subsides in adulthood, at an age in which neurotypical high-risk subjects apparently catch up on the NKCC1/KCC2 ratio, concurrent with the typical onset of SCZ. Other implicated genes include glutamatergic neurotransmission, a process previously associated with polygenic risk for SCZ in the context of co-expression networks (*22*). These observations are consistent with extensive evidence implicating disruptions in E/I balance in SCZ and other neurodevelopmental disorders (*68–71*). Our data add to this earlier work by centering this disruption long before the onset of the clinical disorder based only on genetic risk.

This investigation highlights liquidity as a measure of developmental network dynamics, offering a more rigorous and general framework than previous findings on gene co-expression trajectories (*21*).

Overall, our findings suggest that alterations in gene co-expression liquidity associated with SCZ risk primarily reflect changes in GABA-A–related regulatory mechanisms, possibly influencing transmission/I balance, given the tight functional coupling between inhibitory and excitatory synapses (*72*). It is remarkable that these effects can be observed as a function of cumulative risk based on common genetic variants. The case study on the NKCC1/KCC2 ratio suggests a potentially druggable mechanism of SCZ risk.

## Materials and Methods

### Samples and data preprocessing

For a detailed description, refer to (*21*). In brief, brain tissue samples from neurotypical controls and individuals diagnosed with SCZ of European or African American ancestry with RNA integrity number (RIN) ≥ 6 were obtained from the LIBD Human Brain Repository. All included NC subjects exhibited minimal age-related neuropathology, negative toxicology reports for substance or drug use, and no lifetime diagnoses of psychiatric or neurological disorders, assessed following DSM-IV criteria.

DLPFC samples were taken from Brodmann area 9/46, HP samples included the dentate gyrus, CA3, CA2, CA1 and subicular complex. RNA-seq was performed using the Illumina Ribo-Zero Kit. Gene-level mRNA expression was quantified as RPKMs (reads per kilobase per million mapped reads) and annotated using GENCODE release 25 (GRCh38.p7). Genes with RPKM ≥ 0.1 and minimal floor effects (maximum 20% zeroes) were log-transformed (log_2_(*RPKM* + 1)), excluding mitochondrial genes. Outliers (>3 SDs) were identified and removed. Explicit confounders, i.e., sex, ancestry, RIN, gene assignment rate, ribosomal RNA (rRNA) rate, mitochondrial mapping rate and estimated individual neuronal proportion were regressed out. Age effects were modeled and protected using linear and quadratic terms. Residuals were rank-normalized to address non-normality in expression data.

### Replication dataset

The replication study was conducted using RNA-seq data obtained through the Poly(A) sequencing (*25*). and available on the LIBD repository. The portion of the dataset we used for replication consisted of 91 independent DLPFC brain samples with no level of relationship or overlap to the main dataset, as assessed with identity by decent (IBD) analyses (all overlapping genetic marker values < 0.01). The steps of the preprocessing workflow adopted for Poly(A) data are identical to the ones used for the Ribo-Zero dataset, except gene mapping rate was regressed out in place of gene assignment rate. Regressing gene assignment rate led to adjusted expression values that were distorted and had no correspondence to non-adjusted expression, leading us to regress gene mapping rate in its place.

### Quality Control

For all genotyping datasets we applied quality control (QC) following the standard Rapid Imputation and Computational Pipeline (RICOPILI) (*73*) using PLINK v2 (*74*). We only retained SNPs with a genotype call rate superior to 95%, in Hardy Weinberg Equilibrium (*p* > 1 · 10^−6^) and with a major allele frequency rate over 1% to exclude rare variants. We removed individuals with a sample call rate inferior to 95%, outliers for heterozygosity (F > 0.2) and subjects with non-matching genetic sex corresponding to the reported sex. We also performed an identity by descent analysis using PLINK v2, establishing the degree of relatedness. We included only unrelated subjects in the analyses (*pi*_*hat relatedness* < 0.125).

### Construction of co-expression networks from sliding-age time windows

For each tissue, we identified chronologically sequential sliding windows of 40 subjects, such that each window was shifted by a single sample from the previous one. As described in detail in (*21*), normalized residuals, obtained by regressing out confounders from expression data using linear models (see ***Samples and data preprocessing*** section), were used to construct “signed hybrid” networks with WGCNA’s blockwiseModules function, soft-thresholding positive correlations using a parameter *β* that ensured scale invariance and matched network connectivity across all networks. We discarded any window where the connectivity matching could not be attained (3 for DLPFC, 5 for HP).

### Liquidity

The tendency of genes to vary their co-expression relations in time was quantified through the metrics

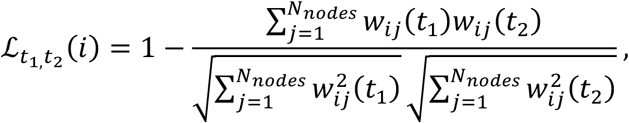

known as *liquidity*, which represents the volatility of the links with gene *i* in the comparison of the two co-expression networks referred to times *t*_1_ and *t*_2_. In the above definition, *w*_*i*j_(*t*) is the weight of the link between the nodes *i* and *j* at time *t*, while *N*_*nodes*_ is the total number of network nodes.

### Detection of maximum increase time points in liquidity scores

For each sliding window, we constructed a gene co-expression network and calculated the liquidity scores for all genes. We then analyzed the median liquidity across all genes as a function of the median age of the subjects within each window. To identify the specific time points at which median liquidity increased most rapidly, corresponding to the stages of maximum variability in co-expression patterns, we computed its first temporal derivative. Prior to differentiation, we smoothed the data using a Savitzky-Golay filter with a five-point window and a second-order polynomial to mitigate noise. We then identified the time points corresponding to the maxima of the first derivative that exceeded a prominence threshold of 0.15. This criterion enabled us to identify peaks that stood out significantly above the surrounding baseline, ensuring we captured only relevant liquidity increase patterns rather than minor local fluctuations.

### Statistical tests for distribution comparison

Unless otherwise specified in the ***Results***, pairs of distributions were compared using the paired Wilcoxon signed-rank test (liquidity of genes in high-vs. low-PRS groups on corresponding time windows, expression of genes in high- vs. low-PRS groups on corresponding time windows) or the Wilcoxon rank-sum test (NKCC1/KCC2 gene expression ratio in high- vs. low-PRS subjects grouped into younger and older cohorts based on various age thresholds within the 5-25 year juvenile range). False discovery rate corrections for multiple comparisons were applied to the results of all statistical tests via the Benjamini-Hochberg procedure; hence, the corresponding p-values were indicated as *p*_*FDR*_. The difference between two distributions was considered statistically significant when *p*_*FDR*_ < 0.05.

### Ancestry scores

To assess genetic ancestry, we implemented a cross-validation algorithm for a lasso and elastic-net regularized generalized linear model using HapMap3 (*75*) as a reference panel, to calculate eigenvalues and assign superpopulations based on the overlap of principal components between our sample and the reference sample. We distinguished individuals in European and African-American ancestry.

### Imputation

Samples were pre-phased (*76*) for imputation using the SHAPEIT v1.0.1 (*77*): marker strand alignments were checked against the 1000 genomes reference alleles. Any SNPs misaligned with the genomic reference were flipped prior to phasing to reduce SNP mismatches during the imputation process. The alternative allele frequency, SNP call rate, and *χ*2 (reference panel vs. data) were calculated, and allele switches and strand flips were determined, comparing ref/alt from the reference panel with the data. Every chromosome was imputed separately.

We performed imputation using the Michigan Imputation Server (*78*), a free online imputation service that creates chunks of 20 Mb size and executes Minimac4 (performing imputation with pre-phased haplotypes with a MaCH algorithm) for each chunk. As a reference Panel, we used the 1000 Genome Phase 3 v5 (*79*). The array build was based on GRCh37 / hg19. Phasing for each chunk was done with Eagle v2.4 (*80*).

After imputation, we retained SNPs with a maximum genotype imputation probability over 0.9 and a minor allele frequency over 1%.

### Principal Component Analysis

To assess principal component analysis (PCA), we used chip-wide SNP data passing specific quality controls. Hence, we included SNPs with an MAF > 1%, a Hardy Weinberg Equilibrium *p*-value *p* > 10^−6^, and a missing rate < 2%. We excluded SNPs located in genomic regions known to exhibit extended linkage disequilibrium or structural variation, such as the Major Histocompatibility Complex region (Chr6:25-35Mb) and the Chromosome 8 inversion region (Chr8:7-13Mb) (*73*). SNPs passing this quality control were pruned to minimize linkage disequilibrium between SNPs (*R*2 < 0.2, 200 SNPs window). With the resulting SNPs, we calculated Eigenvectors and Eigenvalues of the covariance matrix with SNPRelate (*81*) to identify principal components of the genotype data.

### Polygenic score calculation

We calculated PRS (*82*) with PRSice v2 (*83*) using the Odds Ratio (OR) obtained from the third wave of the European and African mixed ancestry SCZ GWAS performed by the Psychiatric Genomics Consortium (PGC) (*8*). The 1000 genome reference served to estimate the LD during clumping, in which we iteratively picked a SNP (index SNP) and removed variants within a certain genomic window that correlated to the index SNP. We removed SNPs in LD within 500 kb to both ends of the index SNP, using an *R*2 threshold of 1 and a *p*-value threshold of 0.1. We additionally calculated PRS for MDD (*84*), AD (*85*), CD (*86*) and RA (*87*).

### Construction of high-PRS and low-PRS subwindows

For each tissue, we identified a sequence of 60-subject sliding windows, each shifted by a single sample relative to the previous one. Then, inside each window, we examined the distribution of the PRS values for the disorder of interest and built the high-PRS and low-PRS subwindows using the subjects with top-25 and bottom-25 PRS values, respectively. These subwindows were required to have significantly different PRS distributions (as verified with Wilcoxon rank-sum test with *p* < 0.05) for the corresponding populations. At the same time, we corrected the subwindow pairs corresponding to a given window so that their distributions for subject variables were statistically indistinguishable, while limiting the statistical discriminability of sequencing variables’ distributions as much as possible. Specifically, we compared the distributions associated with the subject parameters

- First three genomic eigenvariates (PC 1-3), explaining over 90% of the variance
- Age
- Sex
- RIN
- Postmortem interval

and with the sequencing variables

- numMapped (number of reads which successfully mapped to the reference genome during alignment),
- mitoRate (decimal fraction of reads which mapped to the mitochondrial chromosome, of those which map at all),
- totalAssignedGene (decimal fraction of reads assigned unambiguously to a gene, with featureCounts (88) of those in total),
- rRNArate (decimal fraction of reads assigned to a gene whose type is ‘rRNA’, of those assigned to any gene),
- neu (estimated individual neuronal proportion).

Statistical discriminability was determined by the Wilcoxon rank-sum test (*p* < 0.05) for continuous variables and by the z-score test (*p* < 0.05) for discrete variables.

Each sliding window included 60 subjects, of which 25 had high PRS, 25 had low PRS, and 10 had intermediate PRS. The correction process treats the variables in the priority order listed above. If the high- and low-PRS cohorts were characterized by statistically different distributions (*p* < 0.05) for the variable of interest, the algorithm identified the subject swap that made these distributions statistically indistinguishable while minimally altering the high- and low-PRS value distributions and not inducing differences in the previously considered confounding variables. The swap always involved one of the 10 intermediate-PRS subjects and one in the high- or low-PRS cohorts. Figure 4 provides a schematic illustration of this correction algorithm in case the high- and low-PRS groups initially exhibit a statistically significant difference only in their gender proportions. At the end of the comprehensive procedure for confounder correction, we obtained:

- for each sliding window, statistical indistinguishability of the related PRS-informed subwindows with respect to the subject variables;
- for each sliding window with median age within the perinatal and juvenile groups defined in (*21*) (fetal life to 25 years old), statistical indistinguishability of the PRS-informed subwindows with respect to both the subject and sequencing variables.

**Figure 4.**
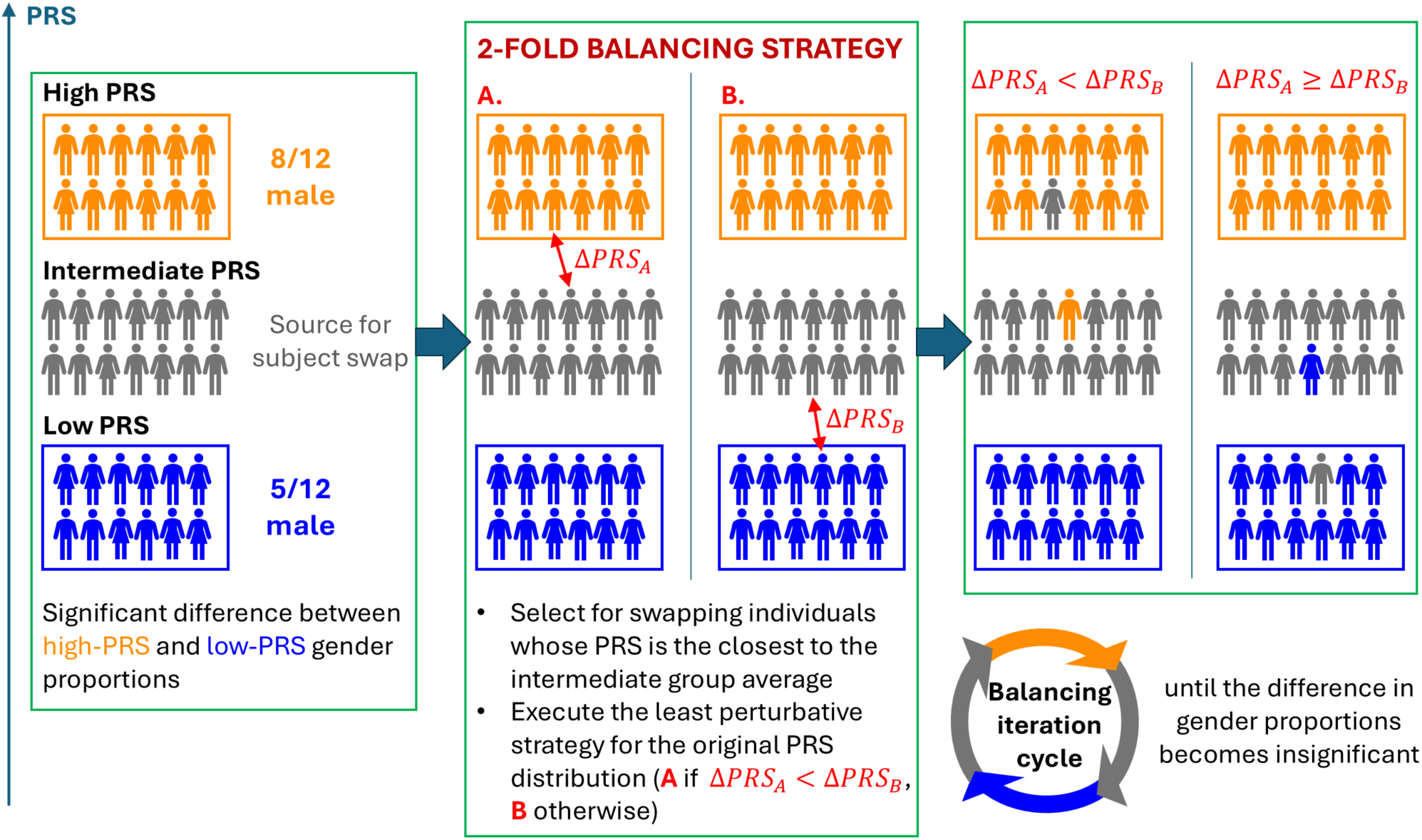
The scheme illustrates the application of the two-fold balancing strategy in case of significant differences for the confounding factor “Sex”, a binary categorical variable, between the high-PRS and low-PRS groups in a given time window. A higher balance of the confounding factor can be achieved by swapping either a subject from the high-PRS group (strategy A) or a subject from the low-PRS group (strategy B) with one from the intermediate-PRS group. In both cases, the swapped subjects are chosen to mitigate the imbalance in gender proportions between the high- and low-risk groups while minimizing the PRS variation. The decision to follow strategy A or B depends on which swapping entails the overall minimum PRS change. The process is iterated until the difference in the “Sex” confounding factor between the high- and low-PRS groups becomes insignificant. For other confounding factors, which are continuous variables, a similar approach is followed, where the goal of swapping is to make the difference between the variable distributions in the high- and low-PRS groups insignificant, while providing minimal perturbation to the PRS distributions. At each stage, the balancing process must also prevent reintroducing significant differences in the distributions of previously corrected variables.

The distributions of all confounding factors across the sliding windows, following the correction of significant differences between high- and low-PRS groups, are shown in Supplementary Figures S6–S29 for all tissue-disorder combinations. All the applied corrections preserved the statistical significance of the difference between PRS distributions in corresponding subwindows.

### Construction of the high-PRS and low-PRS co-expression networks

The construction of high-PRS and low-PRS co-expression networks involved a soft-thresholding procedure analogous to the one employed to construct the 40-subject sliding-age window networks used to examine the temporal variation of the liquidity distribution, briefly outlined in ***Construction of co-expression networks from sliding-age time windows***, and extensively described in detail in (*21*). A relevant difference concerns the connectivity matching condition: while in the previous case, the minimum of the connectivity distribution across windows was taken as a reference, in the case of PRS-informed sliding windows, we chose as reference value the lowest 1^st^ percentile of the distribution, less sensitive to the presence of outliers. This modification was due to the smaller cohorts (25 instead of 40 subjects) used to generate PRS-informed networks, leading to larger fluctuations. We discarded any PRS-informed sliding window for which connectivity matching could not be achieved.

### Construction of co-expression networks from samples of patients with schizophrenia and liquidity evaluation

For each tissue, we sorted the brain samples of individuals with diagnosed SCZ (134 for DLPFC and 112 for HP) by ascending age and grouped them into 50-subject sliding windows. Co-expression networks were constructed using the same connectivity matching procedure as for the high-PRS and low-PRS cohorts. Liquidity could not be computed in the same way as in the NC network cases, due to the absence of patients in the Perinatal age. Therefore, we took as a reference an initial network obtained by merging the earliest-age high-PRS and low-PRS subwindows, containing 25 subjects each.

### Construction of PRS-informed co-expression networks from the replication dataset

We repeated the construction of high- and low-PRS co-expression networks for SCZ using the Poly(A) replication dataset. Since this cohort included fewer subjects than the Ribo-Zero dataset, we adjusted the sliding window size to balance the need for temporal resolution, ensuring windows remained representative of specific neurodevelopmental stages, with the requirement for adequate statistical power. Consequently, we implemented sliding windows of 40 subjects. We verified with the Wilcoxon rank-sum tests that the age distribution of the earliest 40-subject window in the Poly(A) dataset was statistically indistinguishable from that of the initial 60-subject window used in the Ribo-Zero analysis. Within each 40-subject window, we defined high- and low-PRS subgroups consisting of 17 subjects each; the remaining 6 subjects with intermediate PRS values were used to correct confounding factors between the high- and low-risk groups. This 17-to-40 subject ratio was selected as it provided the closest approximation to the 25-to-60 ratio employed in our Ribo-Zero pipeline. The subsequent procedures for network construction from these subwindows were identical to those described in the section ***Construction of the high-PRS and low-PRS co-expression networks***.

### Significance ranking of liquidity differences between high- and low-PRS

To elucidate the different gene contributions to liquidity variations, we compared the distributions {ℒ_*t*0,*t*1_ (*g*), ℒ_*t*0,*t*2_ (*g*), …, ℒ_*t*0,*tN*_(*g*)}*_h_*_*i*g*h*−*SC*Z *PRS*_ and {ℒ_*t*0,*t*1_ (*g*), ℒ_*t*0,*t*2_ (*g*), …, ℒ_*t*0,*tN*_(*g*)}_*lo*w−*SC*Z *PRS*_ within sliding windows *t*_1_, *t*_2_, …, *t*_*N*_, referred to the initial window *t*_0_, for each gene *g* in the DLPFC co-expression networks associated with subjects with high and low PRS for SCZ. The comparison was made using the paired Wilcoxon signed-rank test. After Benjamini-Hochberg correction for multiple comparisons, we associated each gene with a significance level of the difference between temporal liquidity distributions in the networks related to high- and low-PRS subjects for SCZ. Then, we performed GSEA of a ranked gene list based on these − log_10_ *p*_*FDR*_ values, with *p*_*FDR*_ the p-value of the Wilcoxon signed-rank test after the Benjamini-Hochberg correction for multiple comparisons, quantifying the significance of the differences between the high- and low-PRS temporal liquidity distributions for each gene in the DLPFC co-expression networks. The same procedure was repeated to draw a gene-wise significance ranking of liquidity differences between high- and low-PRS for other tissue-disorder combinations. Furthermore, to assess the robustness and temporal specificity of GSEA outcomes, we replicated the workflow in the DLPFC-SCZ case, using the earlier chronological windows immediately following the first as alternative reference points for the liquidity calculation.

### Enrichment methods

GSEA was performed on the gene ranking of − log_10_ *p*_*FDR*_ values using three tools employing specific statistical methodologies:

- STRING (*29*), which utilizes an Aggregate Fold Change (AFC) statistic and a two-sided Kolmogorov-Smirnov (KS) test. The former is a permutation-based, non-parametric test that involves calculating the average of user-provided gene values within each gene set, which is then compared against averages derived from randomized gene sets of identical size. The latter is applied in the case of large gene sets, when the AFC randomization method becomes prohibitively slow, after converting the user-provided gene values to ranks.
- PANTHER (*27*), which generates a reference distribution using all input data values, partitions the list into groups based on Gene Ontology (GO) classification, and estimates the probability of a functional category distribution being randomly drawn from the reference one through the application of the Mann-Whitney Rank-Sum Test (U-Test).
- gProfiler (*28*), which conducts, in the presence of a natural ranking of genes, hypergeometric testing for each possible prefix of the list, starting from the first gene and iteratively adding the subsequent ones (using the ordered gene list option), and produces outputs including the smallest enrichment p-value for each term and the corresponding gene list length.

The adopted multi-tool approach ensured a comprehensive exploration of functional enrichments within the analyzed gene sets, considering the strengths and characteristics of each statistical method employed. The enrichment analysis results, reported in detail in Supplementary File S1, are characterized by an outstanding consistency in the context of the same tool and in comparing different tools.

Furthermore, given that the trajectories of liquidity in high-PRS and low-PRS for SCZ in DLPFC exhibit the greatest divergence in windows with a median age under 16 years, we conducted additional enrichment analysis focusing on this specific period. Considering the small number of time windows, we integrated this part of the analysis by employing clusterProfiler (89), which explicitly utilizes the enrichGO() function for overrepresentation analysis (ORA) based on a hypergeometric test (Fisher’s exact test), comparing the list of significant genes (*p*_*FDR*_ < 0.05) with a list of all genes.

### Significance ranking of mean expression differences between high- and low-PRS

The same procedure used to determine the significance ranking of the gene-wise difference between high-PRS and low-PRS liquidity distributions was applied to compare the temporal profiles of DLPFC expression across the sliding windows in the high- and low-PRS cohorts for SCZ. Specifically, we computed the mean expression, in terms of the standardized residuals, of each gene on the subjects in the high- and low-PRS cohorts and performed the GSEA on the significance ranking of the differences. The same analysis was repeated restricting the observation to the neurodevelopmental phase alone, namely to the sliding windows with median age below 20 years.

### Temporal evolution of GABAergic enrichment via sliding-window GSEA

Following the GSEA described in ***Significance ranking of liquidity differences between high- and low-PRS***, we performed an additional GSEA in the DLPFC-SCZ case using STRING. This procedure used liquidity scores from fixed sliding windows as input, yielding specific enrichment outcomes at each time point, separately for each PRS group. From the complete set of results generated across all sliding windows for both high- and low-PRS groups, we filtered for those containing the term “GABA” in their description. Finally, we examined the significance of these selected pathways over time and across PRS groups.

### Comparing NKCC1/KCC2 gene expression ratio between high- and low-PRS subjects

To construct the NKCC1/KCC2 gene expression ratio, we first adjusted each gene’s cleaned expression (expression value after regression of logRPKM with confounders) to the mean value of its pre-cleaned expression (logRPKM):

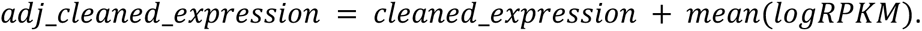

The NKCC1/KCC2 ratio was then computed per subject as

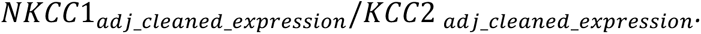

We computed the mean NKCC1/KCC2 ratio per sliding-window network in the DLPFC-SCZ case to visualize how it changes over time across high- and low-PRS networks. Visual inspection reveals that high-PRS networks have increased ratio values at lower ages, with the gap between high and low risk substantially reducing by around 20 years (bottom left panel of Figure 3(d)).

Then, we aimed to quantitatively compare age-dependent changes in NKCC1/KCC2 ratio between high and low-risk subjects. This step was performed on our original data as well as on the separate replication set sequenced with Poly(A) (*25*) and preprocessed using the steps described in ***Replication dataset***. For each set, we created two subject groups: a younger group of under *Y* years old and an older group of over *Y* years old, with the age threshold *Y* spanning the 5-25 juvenile range. Within each age group, subjects were separated into high- and low-risk cohorts corresponding to the top and bottom 40% of the PRS distribution referred to the considered age range, controlling for subject attributes and sequencing variables with the same method described in the ***Construction of high-PRS and low-PRS subwindows*** subsection. Supplementary Figures S33-S44 provide an overview of the distributions of confounding factors following correction for significant differences between the high- and low-PRS cohorts, across the various age thresholds *Y* considered.

Finally, within each age group, we compared the NKCC1/KCC2 ratio of high and low-risk cohorts using the Wilcoxon rank-sum test (Supplementary Table S3).

## DATA AVAILABILITY

The raw RNA-Seq FASTQ files from the LIBD post-mortem dataset, which include data for the dorsolateral prefrontal cortex (DLPFC) and the hippocampus (HP), can be accessed through the Database of Genotypes and Phenotypes (dbGaP) DLPFC: jhpce#bsp2-dlpfc [http://research.libd.org/globus/jhpce_bsp2-dlpfc/index.html]; HP: jhpce#bsp2-hippo [http://research.libd.org/globus/jhpce_bsp2-hippo/index.html]). Additionally, raw genotype data from the LIBD post-mortem study are available via dbGaP under the accession code <u>phs000979.v3.p2</u>. The LIBD post-mortem processed RNA-Seq data, along with the accession codes for the raw RNA-Seq FASTQ files and genotypes used in this study, are also publicly accessible at https://eqtl.brainseq.org/phase2/.

## CODE AVAILABILITY

The scripts used for the analyses conducted in this study are available in a public Zenodo repository at 10.5281/zenodo.14675916.

## ACKNOWLEDGMENTS

We are grateful Dr. Nicola Pedreschi for fruitful discussions about the liquidity metrics. We express our gratitude to Dr. Leonardo Sportelli for support in the identification of metadata referred to the replication dataset, and tools used in the enrichment analysis. We thank Dr. Antonio Lacalamita for useful suggestions concerning the optimization of the computational pipeline used in this study.

## AUTHOR CONTRIBUTIONS

Conceptualization: L.B., D.R.W., and G.P. Data curation: L.B., C.B., G.C.K., and M.P. Formal analysis: L.B., F.D.C., and C.B. Funding acquisition: D.R.W., and G.P. Investigation: L.B., F.D.C., C.B., D.R.W., and G.P. Methodology: L.B., F.D.C., C.B., and G.P. Project administration: G.P. Resources: J.E.K., T.M.H., and D.R.W. Software: L.B., and M.P. Supervision: D.R.W., and G.P. Validation: L.B., F.D.C., C.B., and G.P. Visualization: L.B., and F.D.C. Writing (original draft): L.B., F.D.C., C.B., D.R.W., and G.P. Writing (review and editing): all authors.

## FUNDING

This work was supported by the Research Projects of National Relevance (PRIN) 2022 Prot P2022HNBJX awarded to GP. GCK is supported by a Collaboration Grant from Exprivia Spa to GP under the ministerial decree D.M. n. 352/22 under a collaboration agreement. LB and GP have obtained funding for this work under the National Recovery and Resilience Plan (NRRP), Mission 4 Component 2 Investment 1.4-Call for tender no. 3138 of 16 December 2021 of Italian Ministry of University and Research funded by the European Union–NextGenerationEU, award number: Project code: CN00000013, Concession Decree No. 1031 of 17 February 2022 adopted by the Italian Ministry of University and Research, CUP: D93C22000430001, Project title: “National Centre for HPC, Big Data and Quantum Computing”. Brain procurement, processing and RNA sequencing was funded by the Lieber Institute for Brain Development with a contribution from the Brainseq Consortium.

## COMPETING INTERESTS

D.R.W. serves on the Scientific Advisory Boards of Sage Therapeutics and Pasithea Therapeutics. G.P., F.D.C. and G.C.K. received lecture fees from Lundbeck. The remaining authors declare no competing interests.

