## Supplementary Information for "Liquidity of gene co-expression trajectories across the lifespan highlights delayed maturation and the perinatal GABA switch in schizophrenia risk"

***Demographic characteristics of the LIBD Ribo-Zero and Poly(A) datasets.*** The demographic profiles of the Ribo-Zero and Poly(A) datasets are summarized in Table S1, which details the age, sex, ancestry distribution, and number of genes for both cohorts.

*Table S1. Demographic features of the LIBD Ribo-Zero and Poly(A) datasets. NC, neurotypical controls; SCZ, patients with schizophrenia; AA, African American; EUR, European; DLPFC, dorsolateral prefrontal cortex; HP, hippocampus.*

| Dataset | Region | NC sample size | SCZ sample size | Ancestry AA/EUR | Male (Female) [ratio] | Age mean ± sd (years) | Age range (years) | Number of genes |
|---|---|---|---|---|---|---|---|---|
| Ribo-Zero | DLPFC | 263 | 134 | NC: 150/113<br>SCZ: 69/65 | NC: 178 (85) [2.09]<br>SCZ: 93 (41) [2.27] | NC: $34.6 \pm 22.3$<br>SCZ: $49.9 \pm 16.0$ | NC: -1 – 84<br>SCZ: 17 – 97 | NC: 21129<br>SCZ: 21264 |
| | HP | 278 | 112 | NC: 152/126<br>SCZ: 60/52 | NC: 191 (87) [2.20]<br>SCZ: 75 (37) [2.03] | NC: $35.8 \pm 21.3$<br>SCZ: $50.5 \pm 15.2$ | NC: -1 – 84<br>SCZ: 17 – 97 | NC: 20421<br>SCZ: 20265 |
| Poly(A) | DLPFC | 91 | 0 | 37/54 | 65 (26) [2.50] | $27.12 \pm 24.08$ | -1 – 85 | 20428 |

***Comparing PRS-informed liquidity distributions in co-expression networks for AD, CD, and RA.*** Figure S1 shows the liquidity profiles referred to the DLPFC and HP co-expression networks obtained from high- and low-PRS subjects, for the considered polygenic disorders of a non-psychiatric nature AD, CD, and RA.

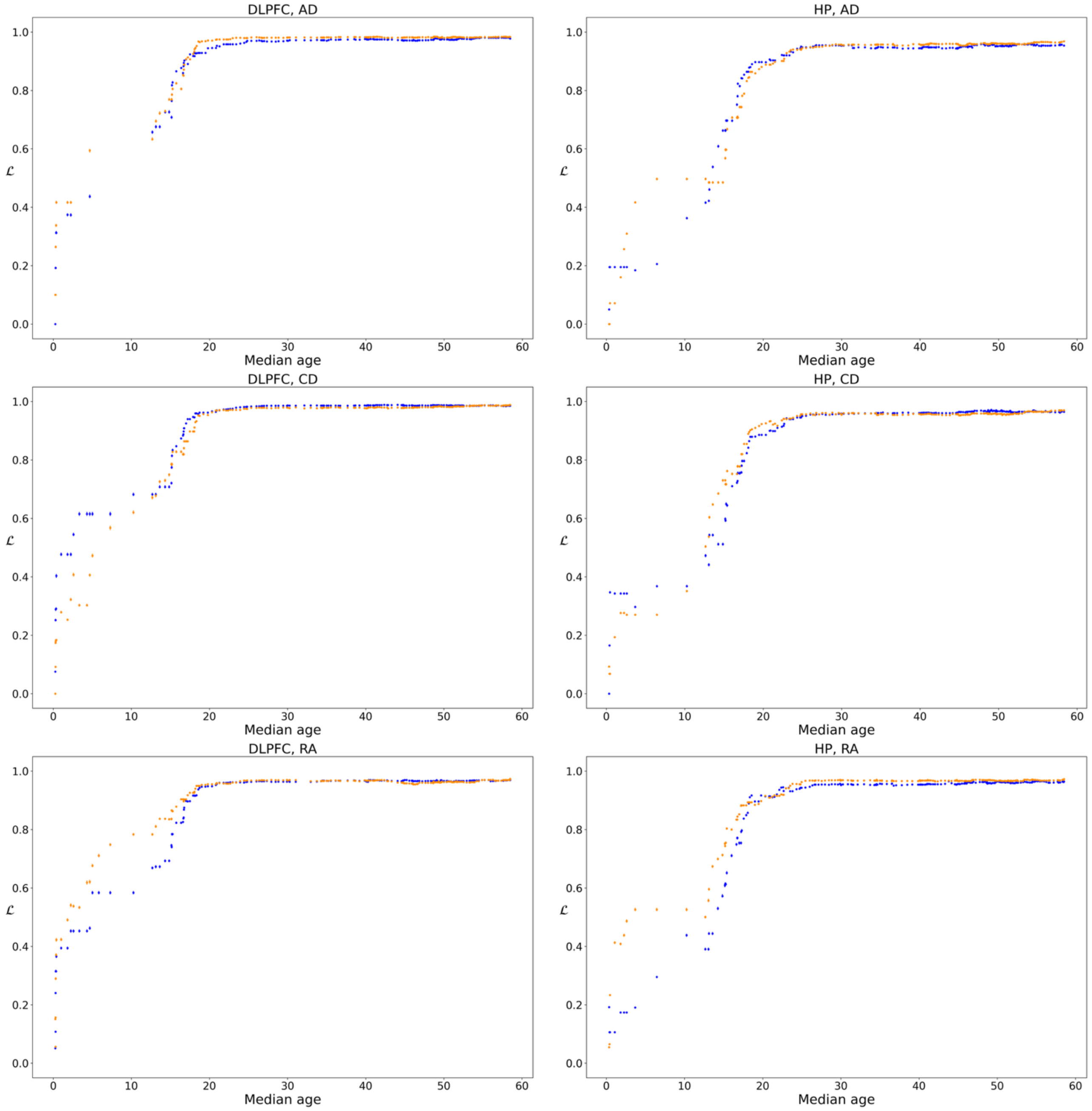


*Figure S1. Liquidity profiles of genes in the DLPFC and HP co-expression networks obtained from high-PRS (orange) and low-PRS (blue) subjects for the considered polygenic disorders of a non-psychiatric nature, AD, CD, and RA, as a function of the median age of the 60-subject sliding windows from which the high-PRS and low-PRS subwindows have been extracted. Dots indicate the median of the liquidity distributions and error bars represent the 95% confidence interval of the median, determined from 10000 bootstrap resamples. DLPFC, dorsolateral prefrontal cortex; HP, hippocampus; AD, Alzheimer's disease; CD, Crohn's disease; RA, rheumatoid arthritis.*

***Comparing PRS-informed liquidity distributions in DLPFC co-expression networks for SCZ in the Poly(A) dataset.*** Figure S2 shows the liquidity profiles referred to the DLPFC co-expression networks obtained from high- and low-PRS subjects for SCZ in the Poly(A) dataset.

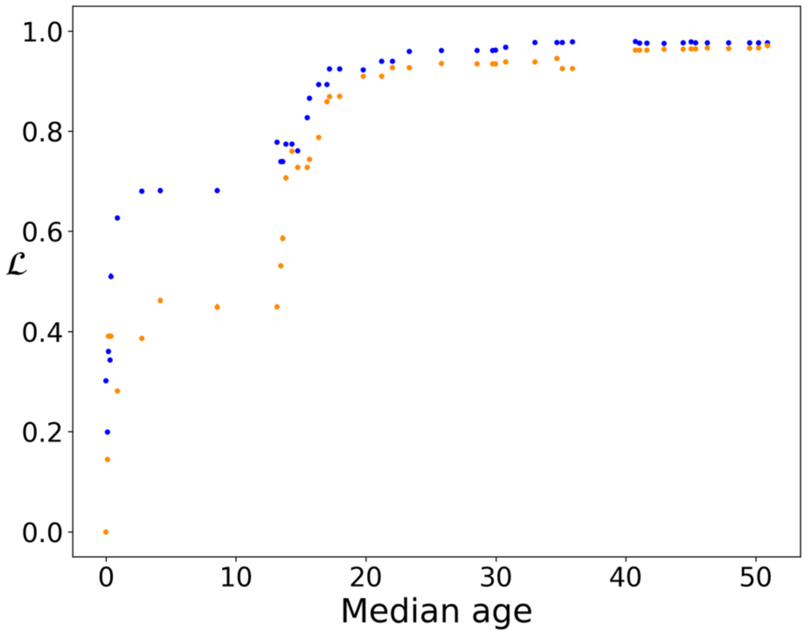


*Figure S2. Liquidity profiles of genes in the DLPFC co-expression networks obtained from high-PRS (orange) and low-PRS (blue) subjects for SCZ in the Poly(A) dataset, as a function of the median age of the 60-subject sliding windows from which the high-PRS and low-PRS subwindows have been extracted. Dots indicate the median of the liquidity distributions and error bars represent the 95% confidence interval of the median, determined from 10000 bootstrap resamples.*

***Comparing high- and low-PRS liquidity distributions of co-expression networks.*** Supplementary Figure S3 shows the comparison between the liquidity distributions of all genes in co-expression networks associated with high- and low-PRS subjects for different combinations of tissue and disorder.

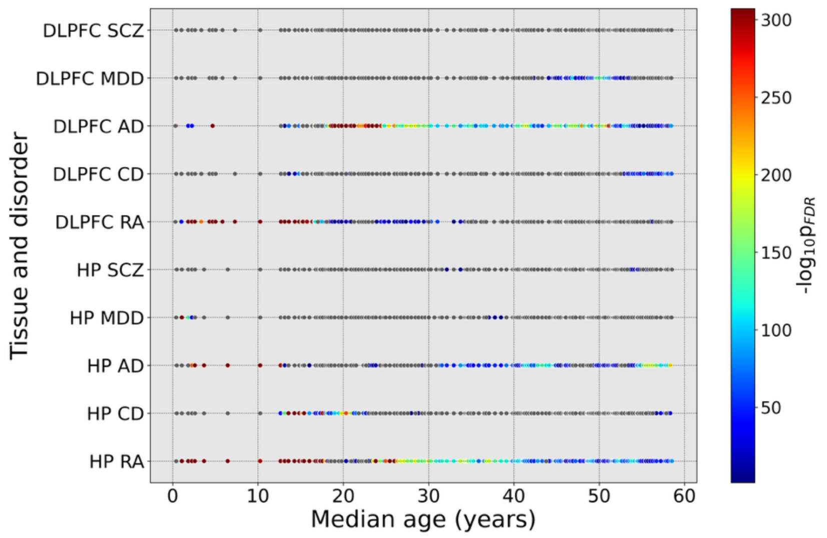


*Figure S3. Comparison between the liquidity distributions of all genes in co-expression networks associated with high- and low-PRS subjects for different combinations of tissue and disorder, in terms of the p-value $p_{FDR}$ (with Benjamini-Hochberg correction for multiple comparisons in all sliding windows) of a Wilcoxon signed-rank test with the alternative hypothesis that the liquidity distribution referred to high-PRS subjects is higher than that of the low-PRS cohort; sliding windows for which the difference between the high- and low-PRS liquidity distributions is not significant are represented as grey dots. Results are expressed as a function of the median age of the 60-subject sliding windows from which the high-PRS and low-PRS cohorts have been extracted. DLPFC, dorsolateral prefrontal cortex; HP, hippocampus; SCZ, schizophrenia; MDD, major depressive disorder; AD, Alzheimer's disease; CD, Crohn's disease; RA, rheumatoid arthritis.*

**Comparing mean expression distributions in the PRS-informed sliding window framework.** Figure S4 shows the time evolution across sliding windows of mean gene expression distributions related to subjects at high- and low-PRS for the considered brain tissues and polygenic disorders of psychiatric nature (SCZ and MDD). Figure S5 shows analogous results for the considered non-psychiatric polygenic disorders (AD, CD, and RA).

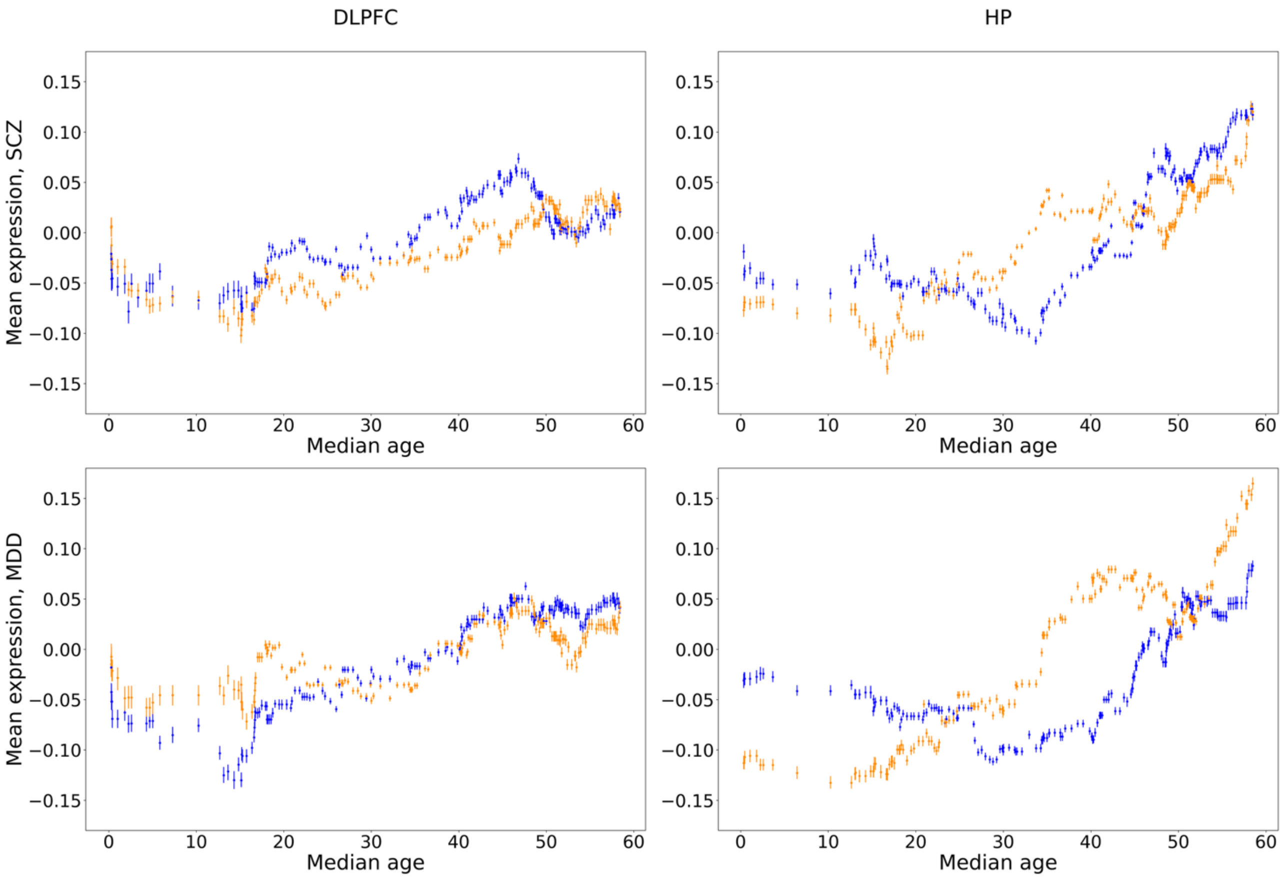


*Figure S4. Mean expression, in terms of the standardized residuals, of genes in the DLPFC (left column) and HP (right column) samples related to the high-PRS (orange) and low-PRS (blue) subjects within each sliding window, for the considered polygenic disorders of psychiatric nature (rows), as a function of the median sliding window age. Dots indicate the median of the distributions and error bars represent the 95% confidence interval of the median, determined from 10000 bootstrap resamples. DLPFC, dorsolateral prefrontal cortex; HP, hippocampus; SCZ, schizophrenia; MDD, major depressive disorder.*

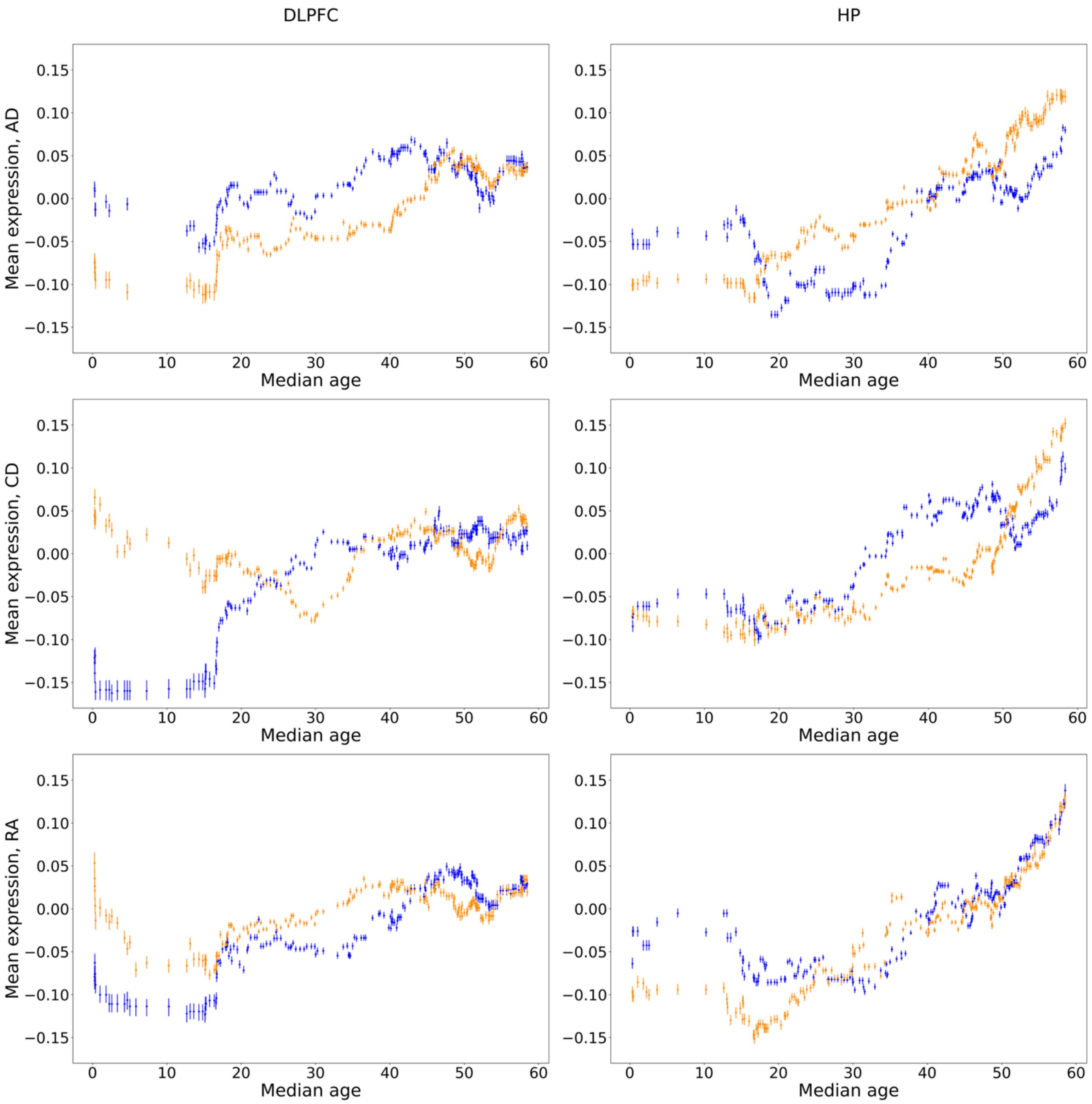


*Figure S5. Mean expression, in terms of the standardized residuals, of genes in the DLPFC (left column) and HP (right column) samples related to the high-PRS (orange) and low-PRS (blue) subjects within each sliding window, for the considered polygenic disorders of a non-psychiatric nature (rows), as a function of the median sliding window age. Dots indicate the median of the distributions and error bars represent the 95% confidence interval of the median, determined from 10000 bootstrap resamples. DLPFC, dorsolateral prefrontal cortex; HP, hippocampus; AD, Alzheimer's disease; CD, Crohn's disease; RA, rheumatoid arthritis.*

***GSEA on the significance ranking of the gene-wise difference between high-PRS and low-PRS liquidity for SCZ in the DLPFC.*** Supplementary File S1 reports the outcomes of the STRING, PANTHER, and gProfiler functional enrichment analyses, from the gene ranking of $-\log_{10} p_{FDR}$ scores, with $p_{FDR}$ the p-value of the Wilcoxon signed-rank test comparing the high- and low-PRS liquidity distributions, after Benjamini-Hochberg correction for multiple comparisons on all genes.

The three tools return a coherent and convergent cluster unequivocally pointing to the GABA neurotransmission pathway and, especially, the GABA-A receptor: (i) *GABA-gated chloride ion channel activity*, *GABA receptor activity*, *GABA-A receptor activity* and *ligand-gated monoatomic anion channel activity*, among molecular function; (ii) *GABA receptor complex* and *GABA-A receptor* *complex*, for cellular component; and (iii) *synaptic transmission, GABAergic* and *regulation of postsynaptic membrane potential*, within biological process.

***GSEA on the significance ranking of the gene-wise difference between high-PRS and low-PRS liquidity for SCZ in DLPFC sliding windows with median age below 16.*** Supplementary File S2 reports the outcomes of the STRING, PANTHER and gProfiler functional enrichment analyses, from the gene ranking of $-log_{10}p_{FDR}$ *scores, with* $p_{FDR}$ the p-value of the Wilcoxon signed rank test comparing the high- and low-PRS liquidity distributions for median age below 16, after Benjamini-Hochberg correction for multiple comparisons on all genes. Supplementary File S3 reports the outcomes of the clusterProfiler statistical overrepresentation test, which compares the list of genes showing significant ($p_{FDR} < 0.05$) differences between high- and low-PRS liquidity for SCZ in DLPFC sliding windows with median age below 16, with the set of all genes.

***Comparing the high- and low-PRS gene expression of GABA-A genes.*** Table S2 reports the results of the Wilcoxon signed-rank test to compare the high and low-PRS mean expressions of the critical genes involved in the gene ontologies *GABA-A receptor activity* (molecular function) and *GABA-A receptor complex* (cellular component) returned by the GSEA on liquidity in the DLPFC SCZ case.

*Table S2. Results of the Wilcoxon signed-rank test to compare the high- and low-PRS gene expression throughout the age range (Expression) or restricting the analysis to sliding windows with median age below 30 (Expression U30), for the GABA-A genes returned by the STRING, PANTHER and gProfiler functional enrichment analyses on liquidity. While STRING and PANTHER tools returned all genes listed in the table, gene GABRQ was missing in the outcome of gProfiler. Each cell reports the alternative hypothesis of the Wilcoxon signed-rank test, with H and L indicating the expression distributions in the high- and low-PRS cohorts, respectively, and the p-value* $p_{FDR}$ *after Benjamini-Hochberg correction for multiple comparisons on all DLPFC genes.*

| Genes | Expression | Expression U30 |
|---|---|---|
| GABRG3 | $H < L, p_{FDR} = 1.2 \cdot 10^{-11}$ | Not significant |
| GABRA2 | $H > L, p_{FDR} = 2.4 \cdot 10^{-2}$ | $H > L, p_{FDR} = 1.3 \cdot 10^{-2}$ |
| GABRG1 | $H > L, p_{FDR} = 4.0 \cdot 10^{-14}$ | $H > L, p_{FDR} = 4.1 \cdot 10^{-9}$ |
| GABRA1 | Not significant | $H > L, p_{FDR} = 1.2 \cdot 10^{-10}$ |
| GABRB3 | $H < L, p_{FDR} = 4.3 \cdot 10^{-8}$ | Not significant |
| GABRA5 | $H < L, p_{FDR} = 1.1 \cdot 10^{-23}$ | $H < L, p_{FDR} = 1.3 \cdot 10^{-3}$ |
| GABRA3 | $H < L, p_{FDR} = 3.4 \cdot 10^{-12}$ | Not significant |
| GABRB2 | $H > L, p_{FDR} = 5.5 \cdot 10^{-9}$ | $H > L, p_{FDR} = 7.2 \cdot 10^{-8}$ |
| GABRD | $H < L, p_{FDR} < 10^{-20}$ | $H < L, p_{FDR} = 1.8 \cdot 10^{-12}$ |
| GABRA4 | Not significant | $H > L, p_{FDR} = 7.2 \cdot 10^{-8}$ |
| GABRG2 | $H > L, p_{FDR} = 1.0 \cdot 10^{-6}$ | $H > L, p_{FDR} = 4.4 \cdot 10^{-9}$ |
| GABRB1 | Not significant | Not significant |
| GABRQ | Not significant | Not significant |

***Focus on glutamate genes returned by GSEA.*** As described in the *Results*, following the GSEA, we focused on GABA-A related genes in terms of both liquidity and gene expression. Within the gene ontologies associated with GABAergic pathways, we explored the regulation of postsynaptic membrane potential (GO:0060078). The identified genes include GRIA1, GRIA2, GRIA3, GRIA4,

GRID1, GRID2, GRIK1, GRIK2, GRIK3, GRIK4, GRIK5, GRIN1, GRIN2A, GRIN2B, GRIN2C, GRIN2D, GRIN3A, GRM1, and GRM5. However, unlike what was observed for GABA, we did not find any gene ontology cluster specifically related with glutamatergic receptor activity (Supplementary File S1).

It is noteworthy that KCC2 is expressed near both inhibitory and excitatory synapses (*1, 2*) and, in addition to its role in promoting the GABA switch, interacts with kainate receptors to maintain excitatory/inhibitory (E/I) balance (*3*). KCC2 also modulates dendritic spines, AMPA receptor diffusion, and glutamatergic transmission (*4-6*).

These findings are consistent with disruptions in excitatory/inhibitory balance and its role in schizophrenia and other neurodevelopmental disorders (*7-10*).

Overall, our data suggest that alterations in gene co-expression stability primarily result from changes in GABA-A regulation, affecting network properties and leading to secondary disruptions in excitatory transmission, given the close interplay between excitatory and inhibitory synapses (*11*).

***GSEA on the significance ranking of the gene-wise difference between high-PRS and low-PRS liquidity for other combinations of tissue and disorder.*** Supplementary Files S4-S12 report the outcomes of the STRING, PANTHER, and gProfiler functional enrichment analyses, from the gene ranking of $-\log_{10} p_{FDR}$ scores, with the p-value of the Wilcoxon signed-rank test comparing the high- and low-PRS liquidity distribution, after Benjamini-Hochberg correction for multiple comparisons on all genes.

As illustrated, the results demonstrate the distinctive liquidity findings observed in the DLPFC for SCZ (sensitivity), as well as the consistent enrichment of terms across other control conditions (specificity). Specifically, in MDD, GSEA results revealed the involvement of the actin cytoskeleton in the hippocampus, which has been shown to play roles in neurogenesis (*12*) and fear memory formation (*13*) —two processes whose disruptions are associated with the pathogenesis of the disease (*14-17*); Additionally, terms related to phospholipid binding appear suggestive of disease involvement, as brain membrane lipids play crucial roles in depression and anxiety (*18*). Regarding AD, our findings are largely associated with the splicing, processing, and metabolism of RNA and mRNA, as well as with lysosomal system pathways, closely linked to the pathogenetic degeneration of the disease (*19-22*). Lastly, CD, which was chosen as a non-brain disorder, exhibited a diverse pattern of enriched terms, which may initially appear unexpected. However, the enrichment in synaptic signaling could be attributed to relative changes in neurotransmitter levels, which mediate symptoms such as mood alterations and abdominal pain throughout the course of the disease (*23, 24*); furthermore, when considering these terms alongside behavior and locomotor activity, their consistency can be attributed to gut-brain signaling, where intestinal inflammation alters mood, circadian rhythm, and appetite behavior (*25*). Additionally, we identified an enrichment cluster associated with mitochondrial activity; while mitochondrial dysfunction in CD has been predominantly described in epithelial intestinal cells (*26*), and was recently proposed as a novel conceptualization of its pathogenesis (*27*), recent studies have further identified this dysfunction within the gut-brain axis (*28-30*). Mitochondrial dysfunction leads to deficits in oxidative phosphorylation, another enriched molecular function we observed. Other enriched terms previously linked to CD include MHC class II protein complex (*31-33*) and carnitine trafficking activity (*34, 35*).

Overall, these results appear biologically significant and provide specificity and sensitivity to our findings regarding schizophrenia. However, the precise implications of liquidity within each pathology require further investigation beyond the scope of this study.

***Comparing the high- and low-PRS NKCC1/KCC2 gene expression at different ages.*** We investigated the impact of PRS on the NKCC1/KCC2 gene expression dynamics across different life stages by grouping subjects by age, using a sliding threshold $Y$ within the juvenile range (5–25 years) identified in Pergola et al. (*36*). Wilcoxon rank-sum tests revealed that, for subjects of age $\leq Y$, where $Y$ falls between 17 and 22 years, the high-PRS group exhibited a significantly higher NKCC1/KCC2 gene expression ratio compared to the low-PRS group. We validated these findings in the replication set of samples sequenced using Poly(A) RNA sequencing. In both the discovery and replication datasets, no significant genetic risk effect on the NKCC1/KCC2 gene expression ratio is observed in older subjects, regardless of the juvenile age threshold $Y$. A comprehensive overview of these statistical comparisons across the analyzed cohorts is reported in Table S3.

*Table S3. Results of the Wilcoxon rank-sum test comparing NKCC1/KCC2 expression in the DLPFC between high- and low-risk subjects for SCZ, stratified by age group (≤ $Y$ and > $Y$ years) in both the original (Ribo-Zero) and replication (Poly(A)) datasets. The alternative hypothesis is that the NKCC1/KCC2 distribution in high-PRS subjects is statistically displaced towards higher values than in low-PRS subjects. Statistically significant differences are highlighted in boldface. The variable $n_{samples}$ denotes the number of subjects within the age-specific cohorts from which the high-risk and low-risk subgroups were selected, following the procedure detailed in the Materials and Methods section.*

| | $(NKCC1/KCC2)_{high\ PRS} > (NKCC1/KCC2)_{low\ PRS}$? | | | | | | | |
|---|---|---|---|---|---|---|---|---|
| | **Original (Ribo-Zero)** | | | | **Replication (Poly(A))** | | | |
| | **Age $\leq Y$** | | **Age $> Y$** | | **Age $\leq Y$** | | **Age $> Y$** | |
| **Age threshold $Y$** | $p$ | $n_{samples}$ | $p$ | $n_{samples}$ | $p$ | $n_{samples}$ | $p$ | $n_{samples}$ |
| **5** | > 0.05 | 44 | > 0.05 | 206 | **0.00396** | 29 | > 0.05 | 62 |
| **6** | > 0.05 | 45 | > 0.05 | 205 | **0.00396** | 29 | > 0.05 | 62 |
| **7** | > 0.05 | 46 | > 0.05 | 204 | **0.00396** | 29 | > 0.05 | 62 |
| **8** | > 0.05 | 46 | > 0.05 | 204 | **0.00396** | 29 | > 0.05 | 62 |
| **9** | > 0.05 | 47 | > 0.05 | 203 | **0.00396** | 29 | > 0.05 | 62 |
| **10** | > 0.05 | 47 | > 0.05 | 203 | **0.00396** | 29 | > 0.05 | 62 |
| **11** | > 0.05 | 47 | > 0.05 | 203 | **0.00396** | 29 | > 0.05 | 62 |
| **12** | > 0.05 | 47 | > 0.05 | 203 | **0.00396** | 29 | > 0.05 | 62 |
| **13** | > 0.05 | 48 | > 0.05 | 202 | **0.00396** | 30 | > 0.05 | 61 |
| **14** | > 0.05 | 50 | > 0.05 | 200 | **0.02682** | 34 | > 0.05 | 57 |
| **15** | > 0.05 | 52 | > 0.05 | 198 | **0.00504** | 36 | > 0.05 | 55 |
| **16** | > 0.05 | 57 | > 0.05 | 193 | **0.01441** | 39 | > 0.05 | 52 |
| **17** | **0.01764** | 64 | > 0.05 | 186 | **0.02975** | 40 | > 0.05 | 51 |
| **18** | **0.01688** | 67 | > 0.05 | 183 | **0.02796** | 42 | > 0.05 | 49 |
| **19** | **0.02941** | 74 | > 0.05 | 176 | **0.02196** | 43 | > 0.05 | 48 |
| **20** | **0.03802** | 77 | > 0.05 | 173 | **0.02196** | 43 | > 0.05 | 48 |
| **21** | **0.03746** | 79 | > 0.05 | 171 | > 0.05 | 44 | > 0.05 | 47 |
| **22** | **0.03494** | 81 | > 0.05 | 169 | **0.03566** | 45 | > 0.05 | 46 |
| **23** | > 0.05 | 85 | > 0.05 | 165 | **0.04996** | 46 | > 0.05 | 45 |
| **24** | > 0.05 | 87 | > 0.05 | 163 | **0.04996** | 46 | > 0.05 | 45 |
| **25** | > 0.05 | 91 | > 0.05 | 159 | > 0.05 | 47 | > 0.05 | 44 |

***Post-correction distributions of confounding factors in high- and low-PRS cohorts.*** This section presents the distributions of confounding factors within the high- and low-PRS subgroups across the analyzed tissues and datasets. Supplementary Figures S6–S17 illustrate the distributions of subject parameters (PC1, PC2, PC3, Age, Sex, RIN, Postmortem interval) and sequencing variables (numMapped, mitoRate, totalAssignedGene, rRNArate, neu) for the subgroups derived from the 60-subject DLPFC sliding windows. Supplementary Figures S18–S29 display the corresponding results for the HP. These figures show the data after correcting for significant differences between the high- and low-PRS cohorts, following the methodology detailed in the ***Construction of high-PRS and low-PRS subwindows*** section of the Materials and Methods and illustrated in Figure 4 of the main text.

Supplementary Figure S30 reports the p-values from statistical tests comparing these distributions across all tissue and disorder combinations after the correction.
Regarding the Poly(A) replication dataset, Supplementary Figure S31 presents the distributions of confounding factors for the 17-subject risk subgroups derived from the 40-subject DLPFC sliding windows for SCZ, adjusted for any significant differences between the low- and high-PRS cohorts extracted from the same window. The summary of the statistical comparisons between these cohorts for all confounding factors is provided in Supplementary Figure S32.
Supplementary Figures S33–S44 characterize the post-correction distributions of all confounding factors for both the Ribo-Zero and Poly(A) datasets when subjects were grouped into "under $Y$" and "over $Y$" cohorts. These analyses utilized varying age thresholds ($Y$) within the 5–25 year juvenile range, following the methodology described in the ***Comparing the high- and low-PRS NKCC1/KCC2 gene expression at different ages*** section.

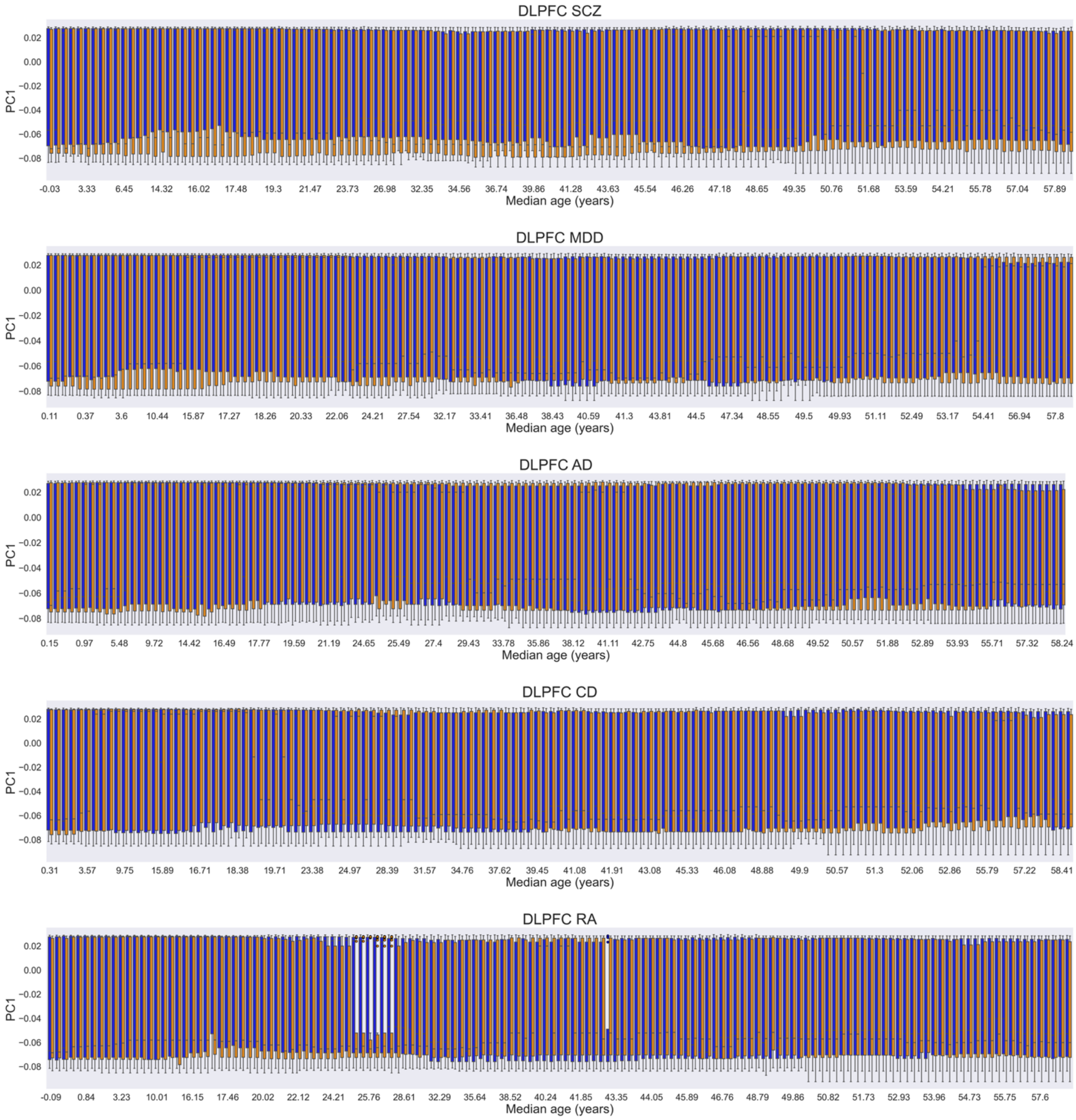


*Figure S6. Distribution of the first genomic eigenvariate (PC1) within subgroups of 25 low-PRS (blue) and 25 high-PRS (orange) subjects for the considered disorders (rows), derived from 60-subject DLPFC sliding windows. Adjacent boxplot pairs represent low- and high-PRS cohorts originating from the same window; distributions are shown after correcting for significant differences in confounding factors. Results are plotted as a function of the median age of the sliding windows from which the high- and low-PRS subgroups were extracted. DLPFC, dorsolateral prefrontal cortex; SCZ, schizophrenia; MDD, major depressive disorder; AD, Alzheimer's disease; CD, Crohn's disease; RA, rheumatoid arthritis.*

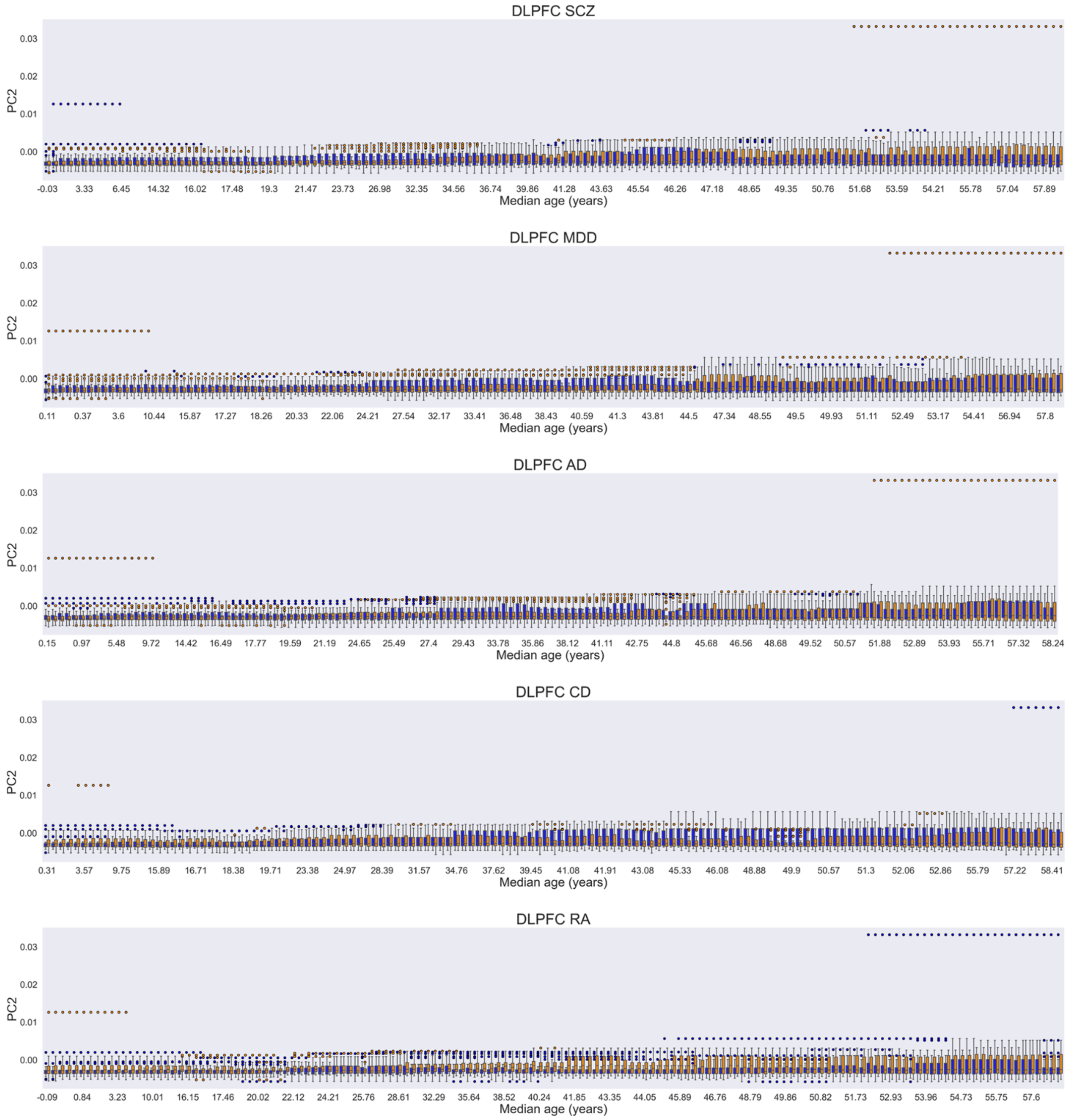


*Figure S7. Distribution of the second genomic eigenvariate (PC2) within subgroups of 25 low-PRS (blue) and 25 high-PRS (orange) subjects for the considered disorders (rows), derived from 60-subject DLPFC sliding windows. Adjacent boxplot pairs represent low- and high-PRS cohorts originating from the same window; distributions are shown after correcting for significant differences in confounding factors. Results are plotted as a function of the median age of the sliding windows from which the high- and low-PRS subgroups were extracted. DLPFC, dorsolateral prefrontal cortex; SCZ, schizophrenia; MDD, major depressive disorder; AD, Alzheimer's disease; CD, Crohn's disease; RA, rheumatoid arthritis.*

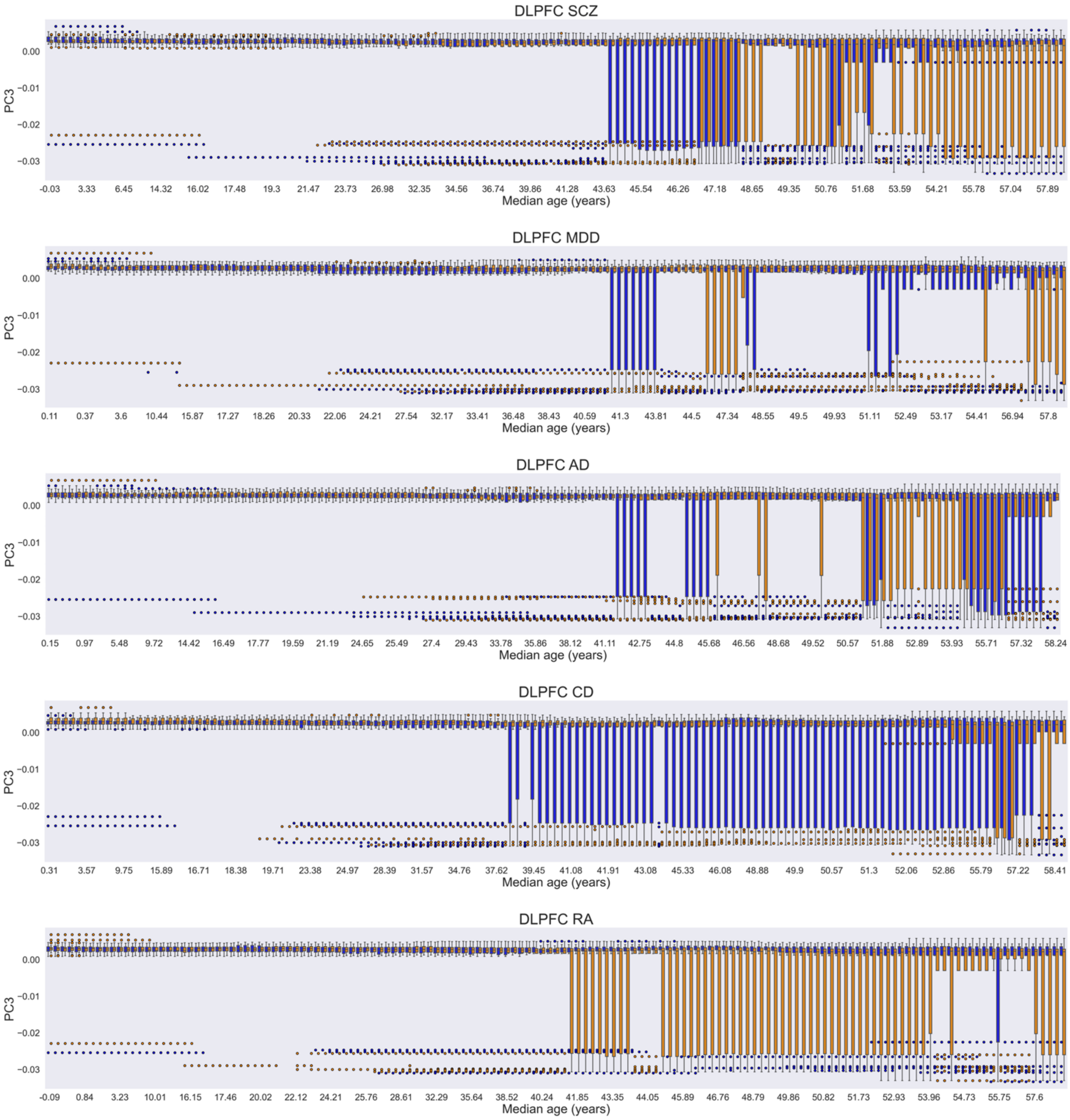


*Figure S8. Distribution of the third genomic eigenvariate (PC3) within subgroups of 25 low-PRS (blue) and 25 high-PRS (orange) subjects for the considered disorders (rows), derived from 60-subject DLPFC sliding windows. Adjacent boxplot pairs represent low- and high-PRS cohorts originating from the same window; distributions are shown after correcting for significant differences in confounding factors. Results are plotted as a function of the median age of the sliding windows from which the high- and low-PRS subgroups were extracted. DLPFC, dorsolateral prefrontal cortex; SCZ, schizophrenia; MDD, major depressive disorder; AD, Alzheimer's disease; CD, Crohn's disease; RA, rheumatoid arthritis.*

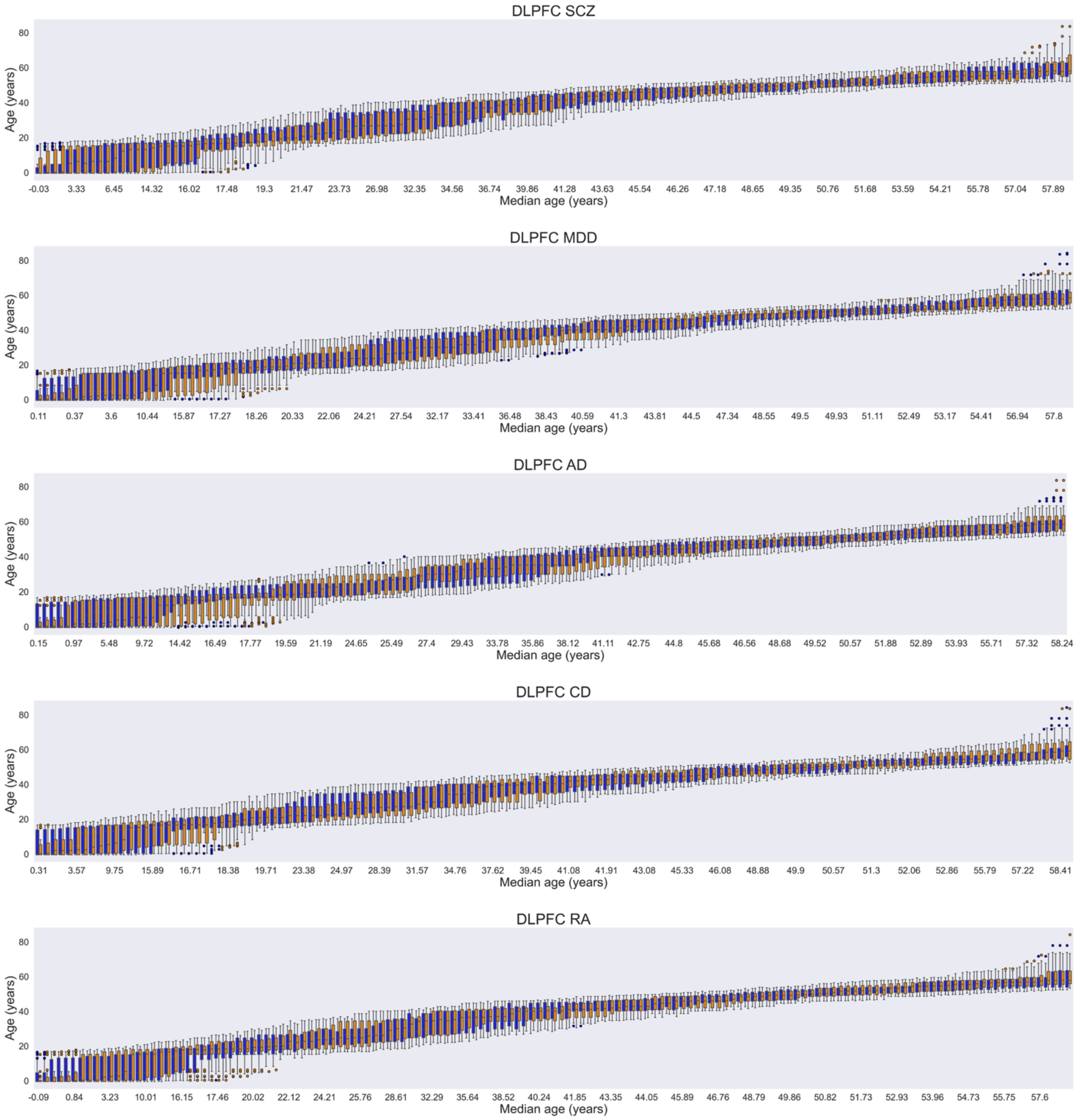


*Figure S9. Age distribution within subgroups of 25 low-PRS (blue) and 25 high-PRS (orange) subjects for the considered disorders (rows), derived from 60-subject DLPFC sliding windows. Adjacent boxplot pairs represent low- and high-PRS cohorts originating from the same window; distributions are shown after correcting for significant differences in confounding factors. Results are plotted as a function of the median age of the sliding windows from which the high- and low-PRS subgroups were extracted. DLPFC, dorsolateral prefrontal cortex; SCZ, schizophrenia; MDD, major depressive disorder; AD, Alzheimer's disease; CD, Crohn's disease; RA, rheumatoid arthritis.*

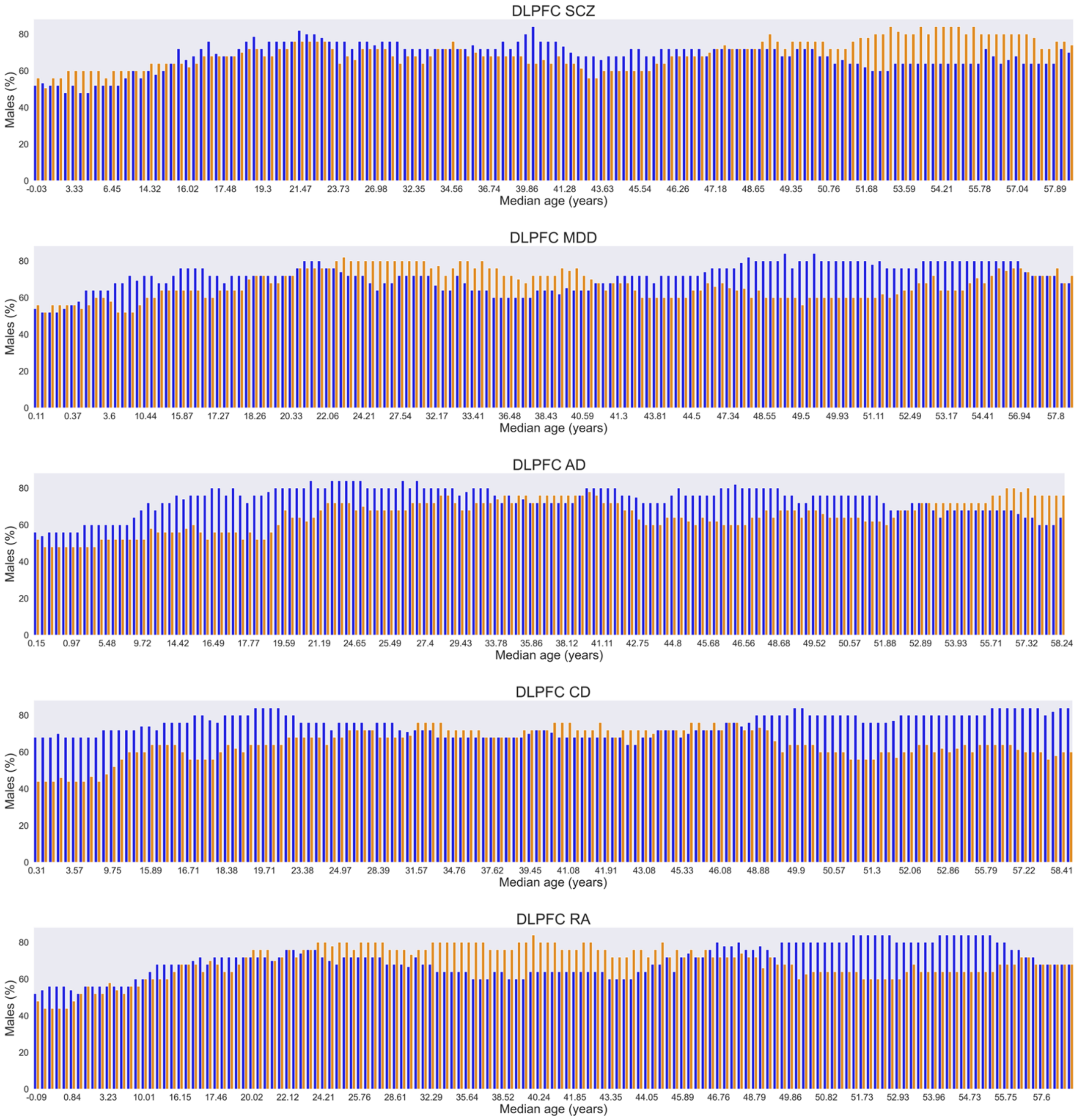


*Figure S10. Percentage of males within subgroups of 25 low-PRS (blue) and 25 high-PRS (orange) subjects for the considered disorders (rows), derived from 60-subject DLPFC sliding windows. Adjacent bar pairs represent low- and high-PRS cohorts originating from the same window; distributions are shown after correcting for significant differences in confounding factors. Results are plotted as a function of the median age of the sliding windows from which the high- and low-PRS subgroups were extracted. DLPFC, dorsolateral prefrontal cortex; SCZ, schizophrenia; MDD, major depressive disorder; AD, Alzheimer's disease; CD, Crohn's disease; RA, rheumatoid arthritis.*

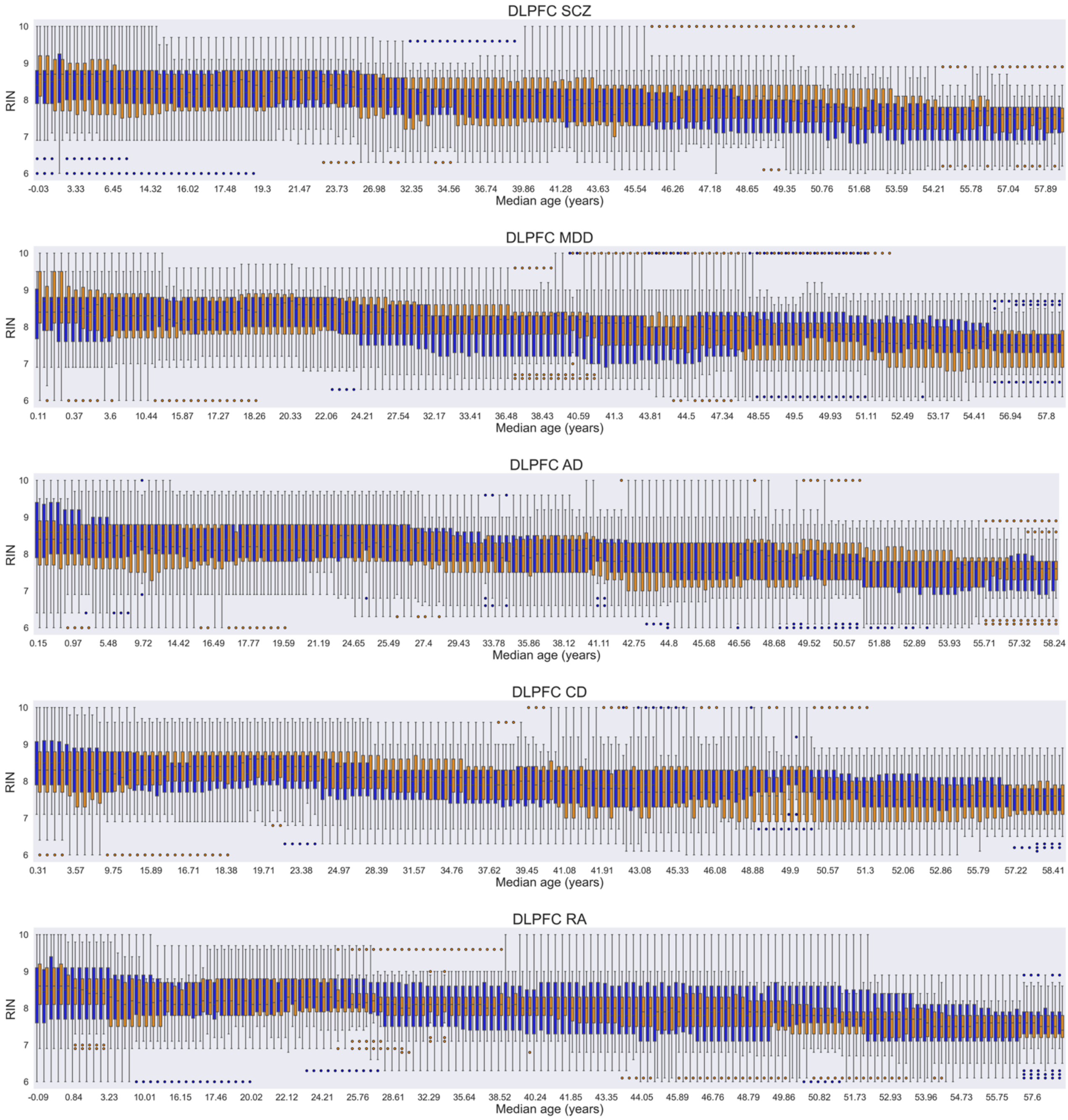


*Figure 11. RNA integrity number (RIN) distribution within subgroups of 25 low-PRS (blue) and 25 high-PRS (orange) subjects for the considered disorders (rows), derived from 60-subject DLPFC sliding windows. Adjacent boxplot pairs represent low- and high-PRS cohorts originating from the same window; distributions are shown after correcting for significant differences in confounding factors. Results are plotted as a function of the median age of the sliding windows from which the high- and low-PRS subgroups were extracted. DLPFC, dorsolateral prefrontal cortex; SCZ, schizophrenia; MDD, major depressive disorder; AD, Alzheimer's disease; CD, Crohn's disease; RA, rheumatoid arthritis.*

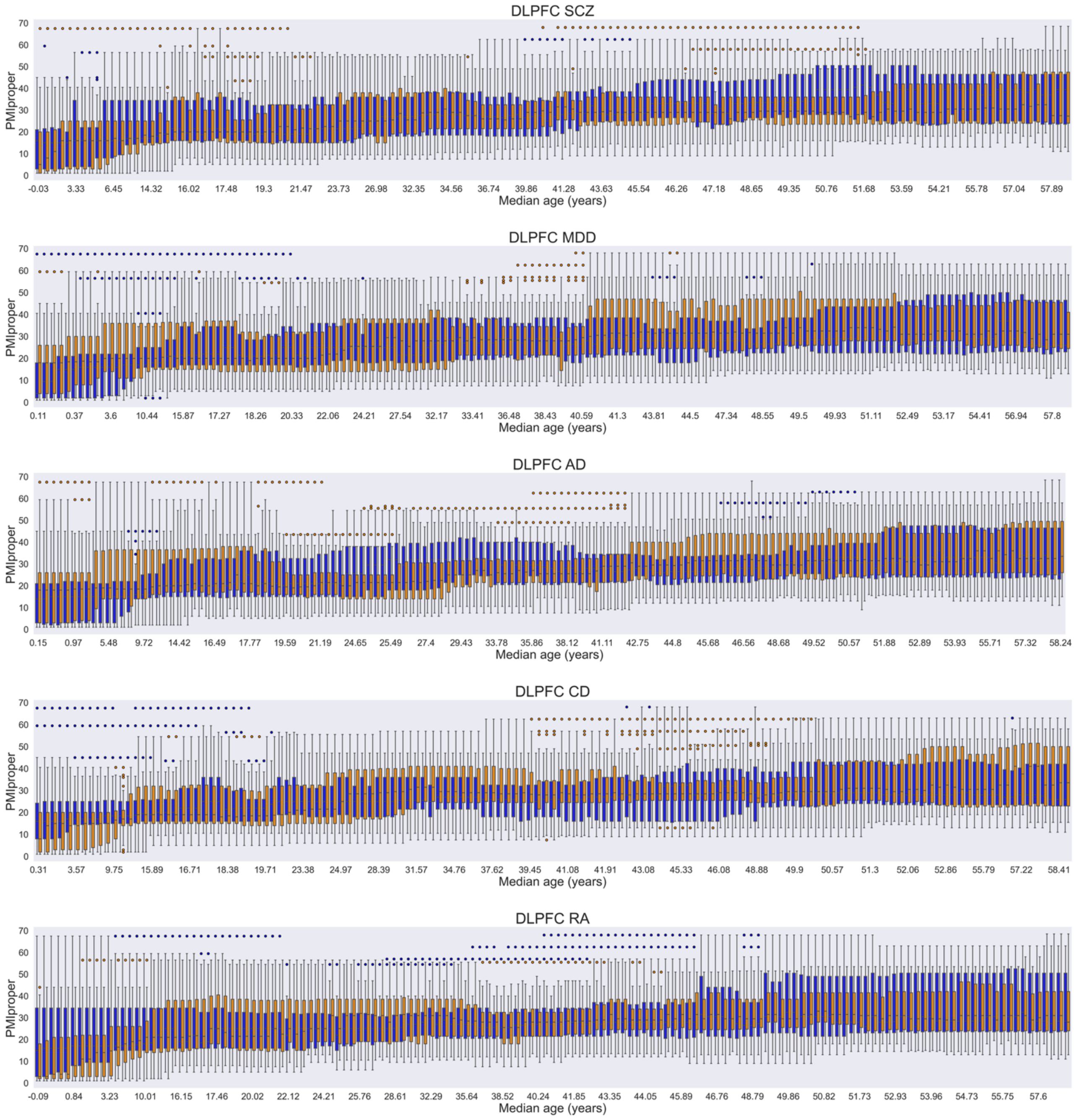


*Figure 12. Distribution of the postmortem interval variable (PMIproper) within subgroups of 25 low-PRS (blue) and 25 high-PRS (orange) subjects for the considered disorders (rows), derived from 60-subject DLPFC sliding windows. Adjacent boxplot pairs represent low- and high-PRS cohorts originating from the same window; distributions are shown after correcting for significant differences in confounding factors. Results are plotted as a function of the median age of the sliding windows from which the high- and low-PRS subgroups were extracted. DLPFC, dorsolateral prefrontal cortex; SCZ, schizophrenia; MDD, major depressive disorder; AD, Alzheimer's disease; CD, Crohn's disease; RA, rheumatoid arthritis.*

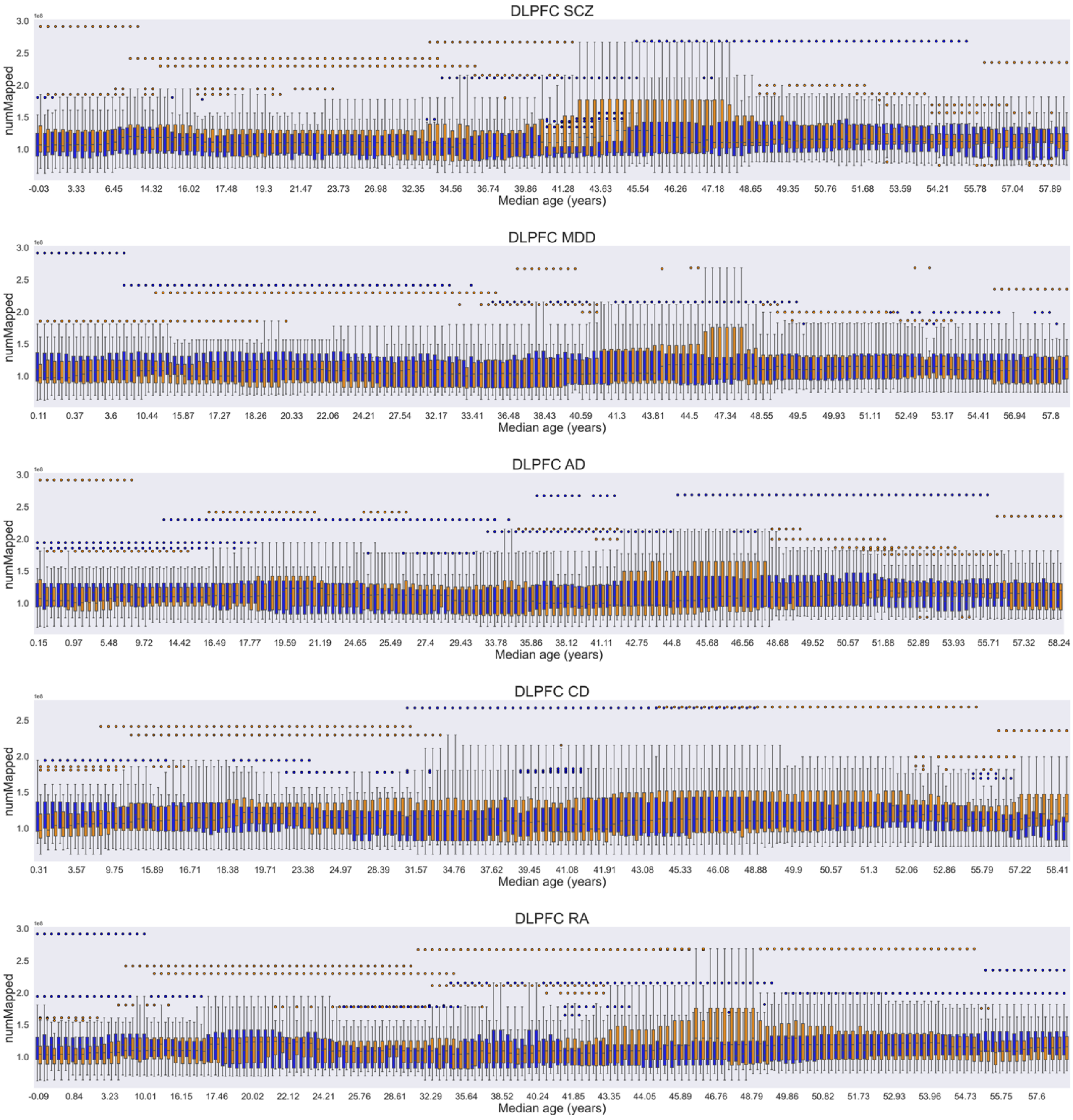


*Figure S13. Distribution of the number of reads which successfully mapped to the reference genome during alignment (numMapped) within subgroups of 25 low-PRS (blue) and 25 high-PRS (orange) subjects for the considered disorders (rows), derived from 60-subject DLPFC sliding windows. Adjacent boxplot pairs represent low- and high-PRS cohorts originating from the same window; distributions are shown after correcting for significant differences in confounding factors. Results are plotted as a function of the median age of the sliding windows from which the high- and low-PRS subgroups were extracted. DLPFC, dorsolateral prefrontal cortex; SCZ, schizophrenia; MDD, major depressive disorder; AD, Alzheimer's disease; CD, Crohn's disease; RA, rheumatoid arthritis.*

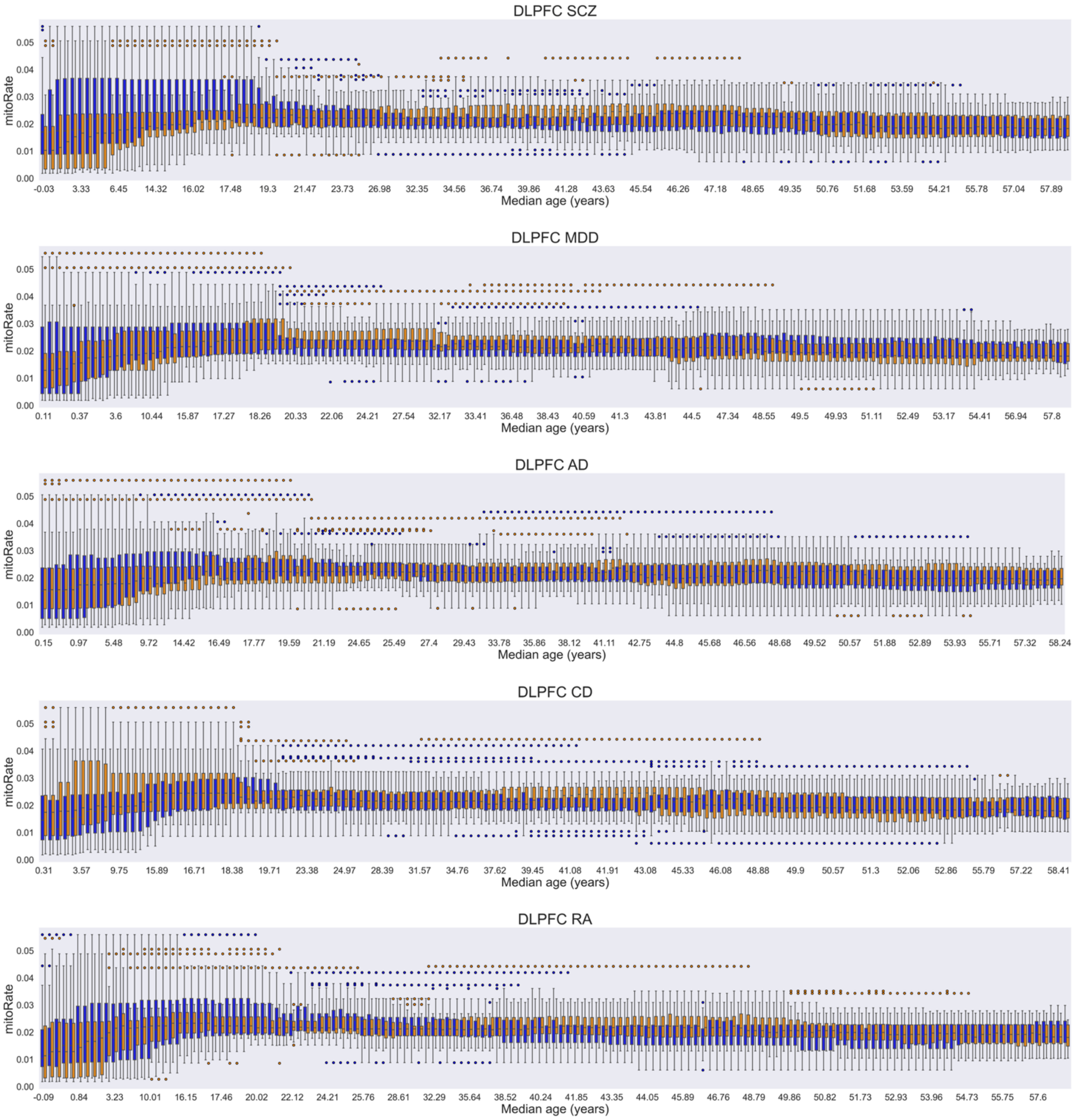


*Figure S14. Distribution of the decimal fraction of reads which mapped to the mitochondrial chromosome, of those which map at all (mitoRate), within subgroups of 25 low-PRS (blue) and 25 high-PRS (orange) subjects for the considered disorders (rows), derived from 60-subject DLPFC sliding windows. Adjacent boxplot pairs represent low- and high-PRS cohorts originating from the same window; distributions are shown after correcting for significant differences in confounding factors. Results are plotted as a function of the median age of the sliding windows from which the high- and low-PRS subgroups were extracted. DLPFC, dorsolateral prefrontal cortex; SCZ, schizophrenia; MDD, major depressive disorder; AD, Alzheimer's disease; CD, Crohn's disease; RA, rheumatoid arthritis.*

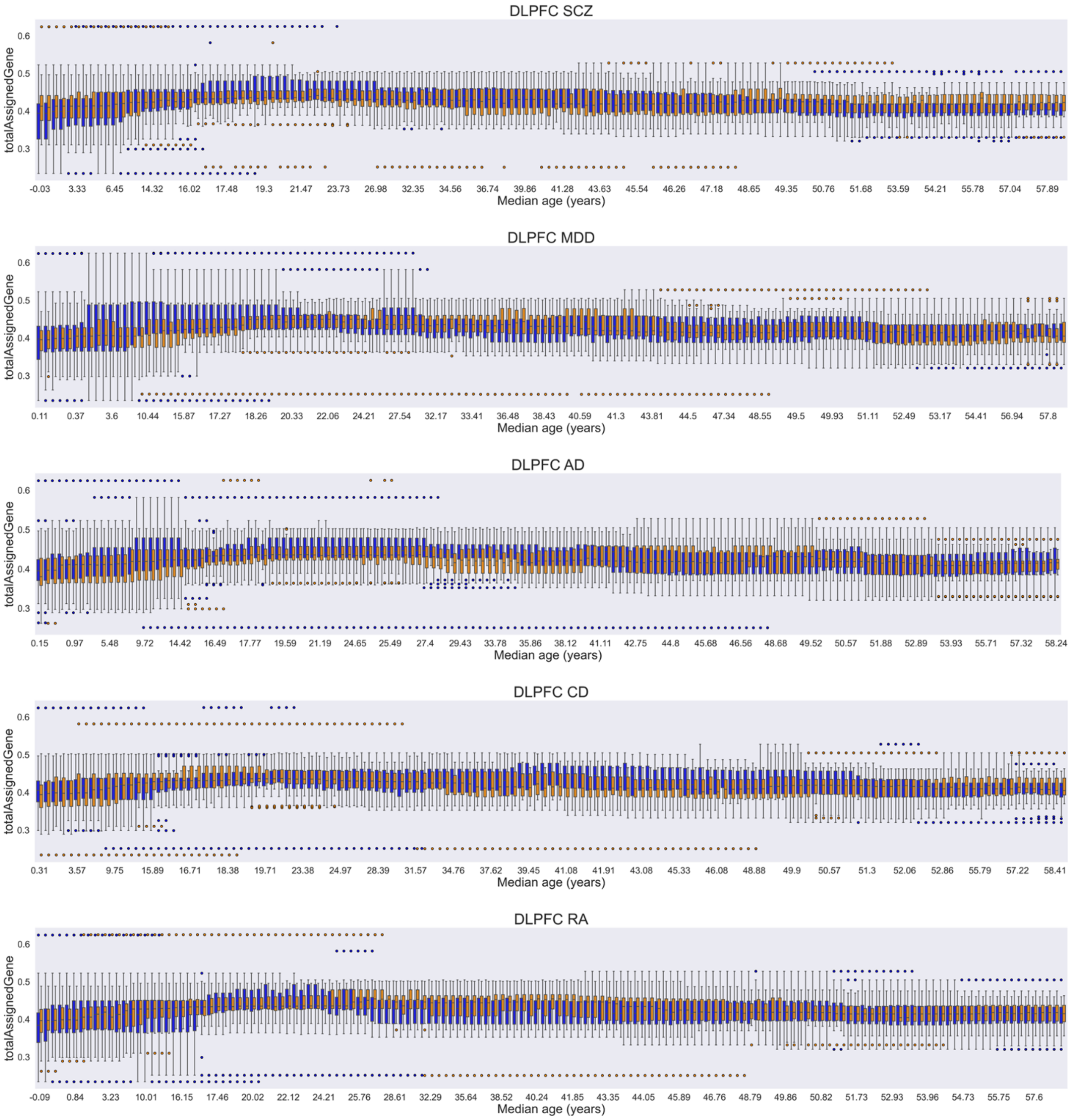


*Figure S15. Distribution of the decimal fraction of reads assigned unambiguously to a gene, with featureCounts of those in total (totalAssignedGene), within subgroups of 25 low-PRS (blue) and 25 high-PRS (orange) subjects for the considered disorders (rows), derived from 60-subject DLPFC sliding windows. Adjacent boxplot pairs represent low- and high-PRS cohorts originating from the same window; distributions are shown after correcting for significant differences in confounding factors. Results are plotted as a function of the median age of the sliding windows from which the high- and low-PRS subgroups were extracted. DLPFC, dorsolateral prefrontal cortex; SCZ, schizophrenia; MDD, major depressive disorder; AD, Alzheimer's disease; CD, Crohn's disease; RA, rheumatoid arthritis.*

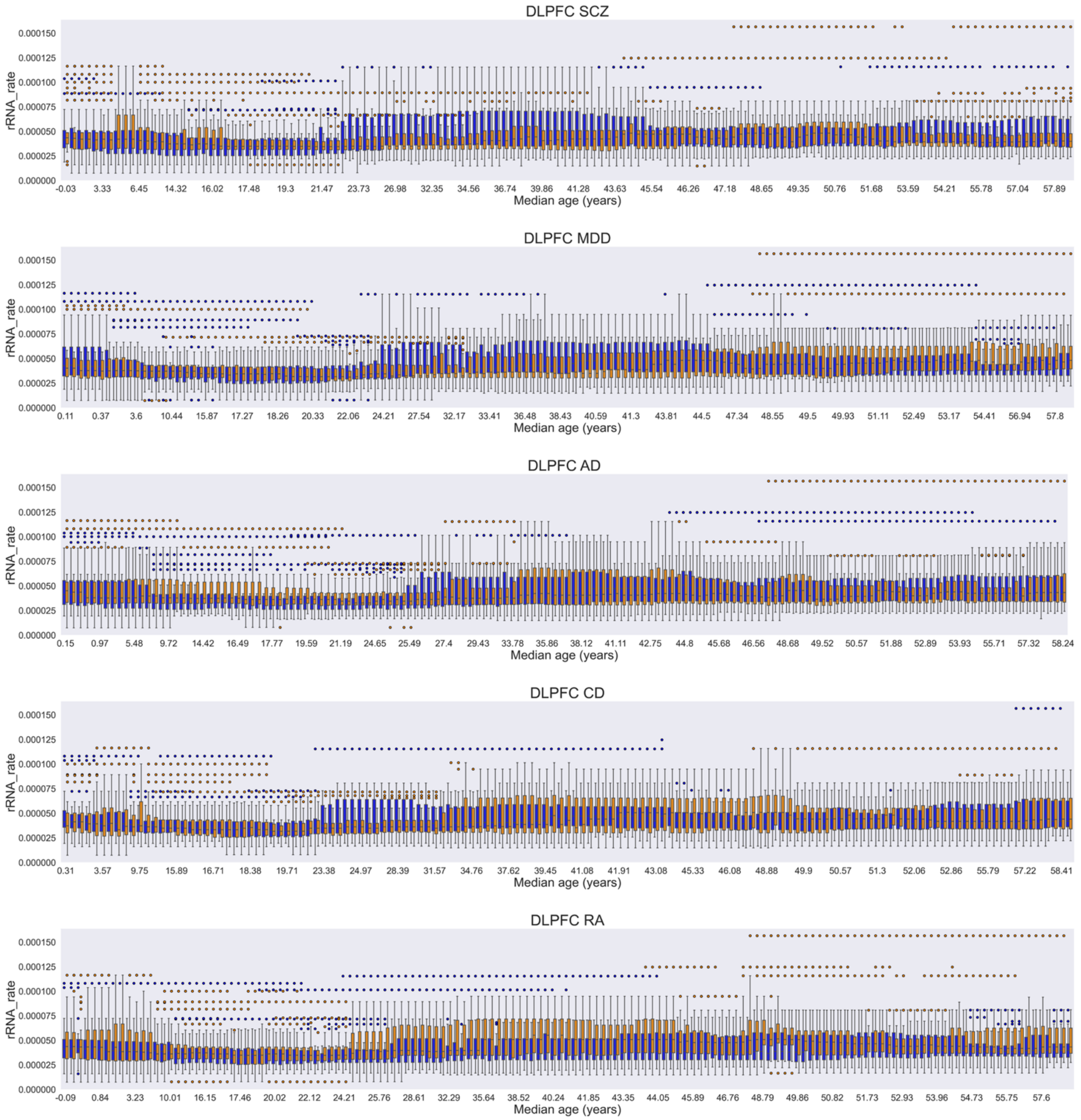


*Figure S16. Distribution of the decimal fraction of reads assigned to a gene whose type is 'rRNA', of those assigned to any gene (rRNA_rate), within subgroups of 25 low-PRS (blue) and 25 high-PRS (orange) subjects for the considered disorders (rows), derived from 60-subject DLPFC sliding windows. Adjacent boxplot pairs represent low- and high-PRS cohorts originating from the same window; distributions are shown after correcting for significant differences in confounding factors. Results are plotted as a function of the median age of the sliding windows from which the high- and low-PRS subgroups were extracted. DLPFC, dorsolateral prefrontal cortex; SCZ, schizophrenia; MDD, major depressive disorder; AD, Alzheimer's disease; CD, Crohn's disease; RA, rheumatoid arthritis.*

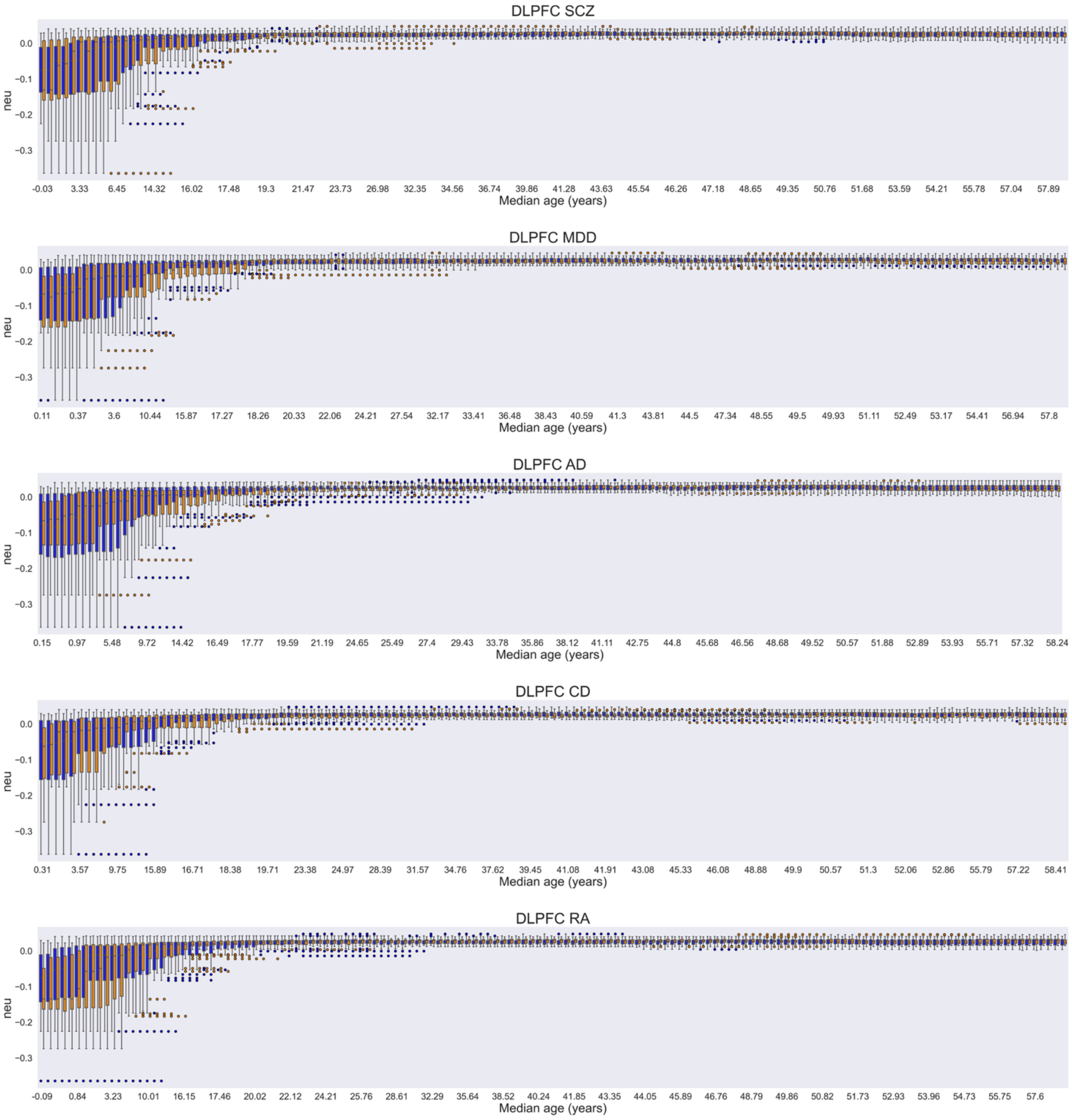


*Figure S17. Distribution of the estimated individual neuronal proportion (neu) within subgroups of 25 low-PRS (blue) and 25 high-PRS (orange) subjects for the considered disorders (rows), derived from 60-subject DLPFC sliding windows. Adjacent boxplot pairs represent low- and high-PRS cohorts originating from the same window; distributions are shown after correcting for significant differences in confounding factors. Results are plotted as a function of the median age of the sliding windows from which the high- and low-PRS subgroups were extracted. DLPFC, dorsolateral prefrontal cortex; SCZ, schizophrenia; MDD, major depressive disorder; AD, Alzheimer's disease; CD, Crohn's disease; RA, rheumatoid arthritis.*

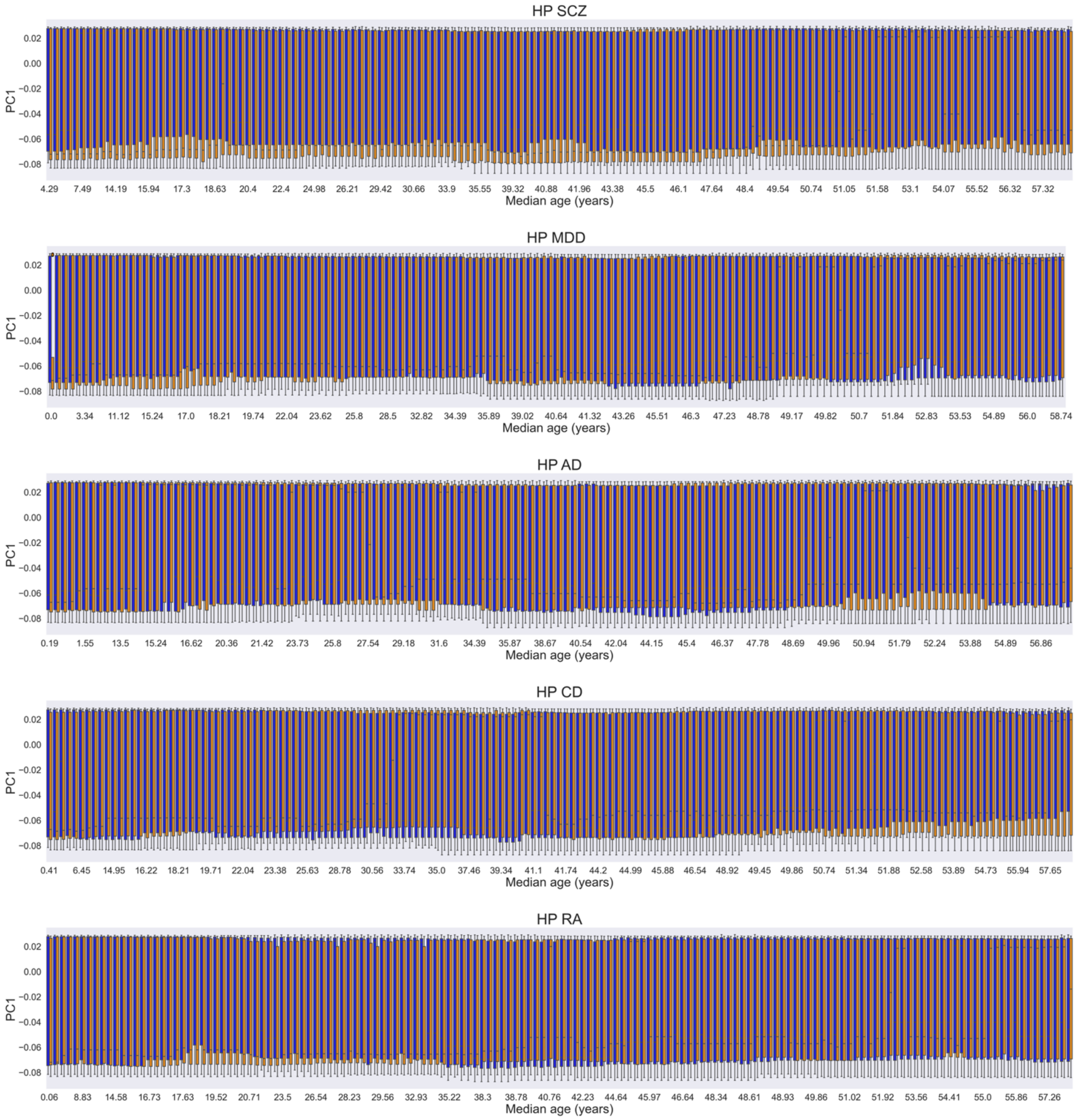


*Figure S18. Distribution of the first genomic eigenvariate (PC1) within subgroups of 25 low-PRS (blue) and 25 high-PRS (orange) subjects for the considered disorders (rows), derived from 60-subject HP sliding windows. Adjacent boxplot pairs represent low- and high-PRS cohorts originating from the same window; distributions are shown after correcting for significant differences in confounding factors. Results are plotted as a function of the median age of the sliding windows from which the high- and low-PRS subgroups were extracted. HP, hippocampus; SCZ, schizophrenia; MDD, major depressive disorder; AD, Alzheimer's disease; CD, Crohn's disease; RA, rheumatoid arthritis.*

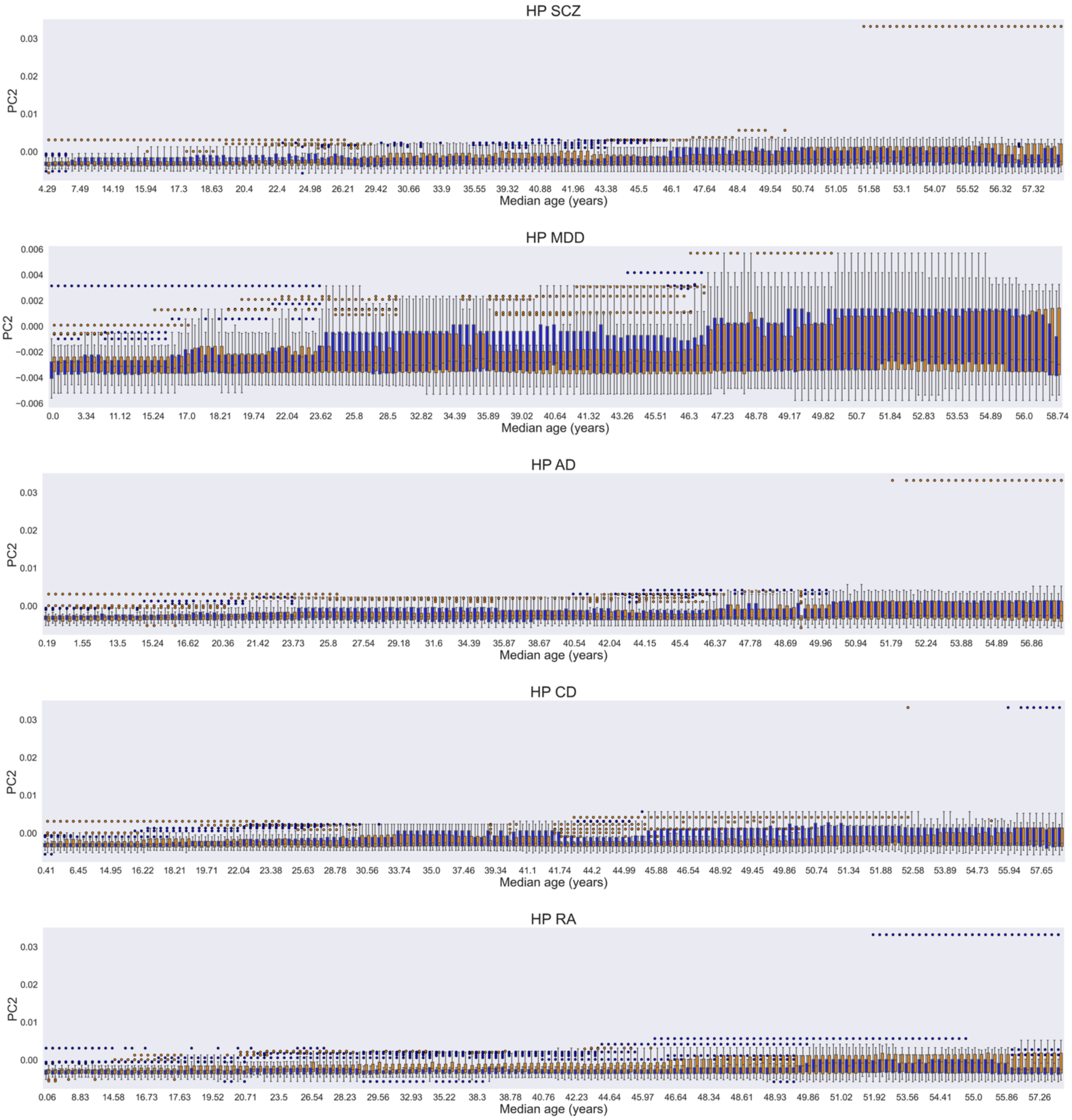


*Figure S19. Distribution of the second genomic eigenvariate (PC2) within subgroups of 25 low-PRS (blue) and 25 high-PRS (orange) subjects for the considered disorders (rows), derived from 60-subject HP sliding windows. Adjacent boxplot pairs represent low- and high-PRS cohorts originating from the same window; distributions are shown after correcting for significant differences in confounding factors. Results are plotted as a function of the median age of the sliding windows from which the high- and low-PRS subgroups were extracted. HP, hippocampus; SCZ, schizophrenia; MDD, major depressive disorder; AD, Alzheimer's disease; CD, Crohn's disease; RA, rheumatoid arthritis.*

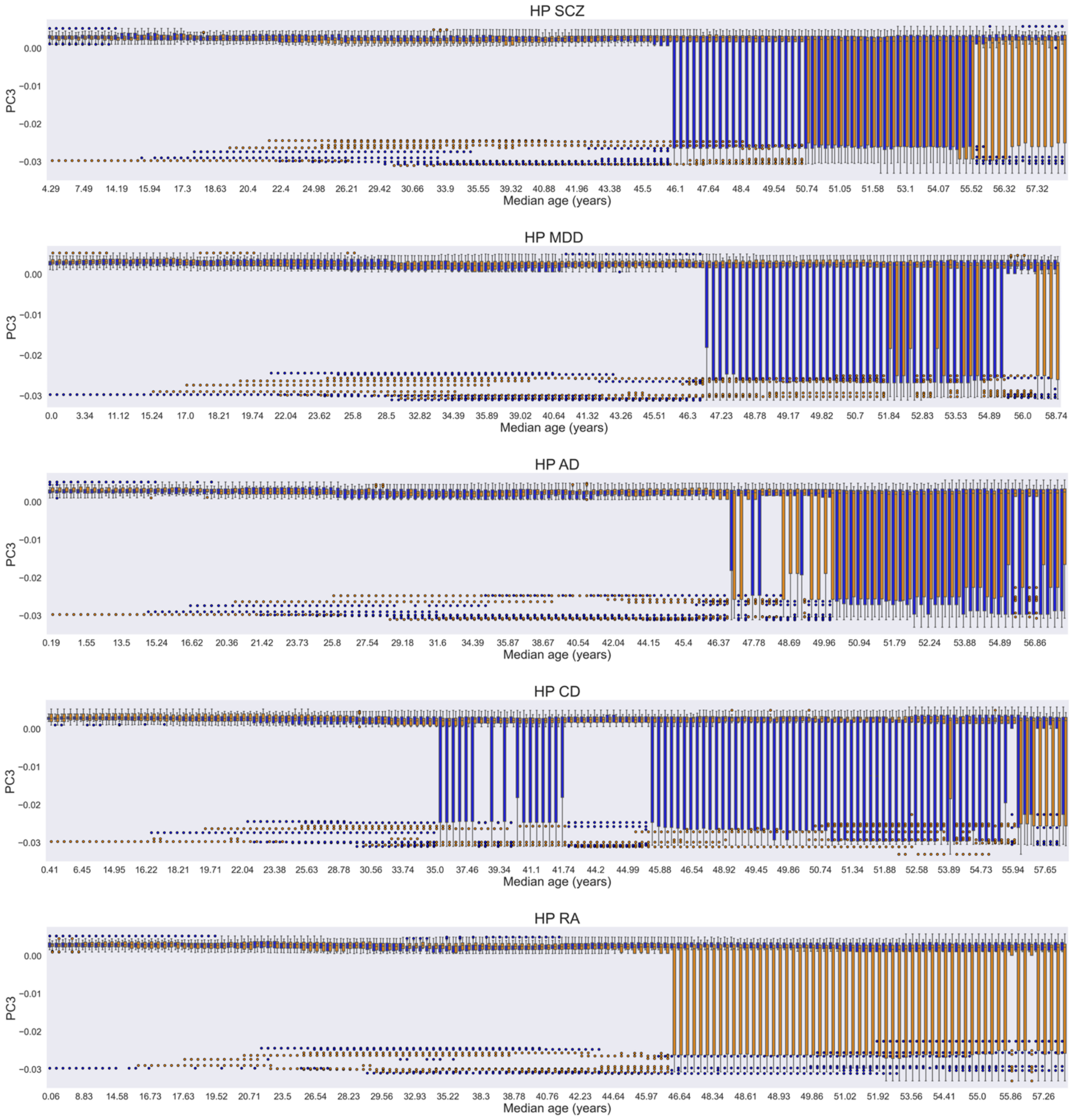


*Figure S20. Distribution of the third genomic eigenvariate (PC3) within subgroups of 25 low-PRS (blue) and 25 high-PRS (orange) subjects for the considered disorders (rows), derived from 60-subject HP sliding windows. Adjacent boxplot pairs represent low- and high-PRS cohorts originating from the same window; distributions are shown after correcting for significant differences in confounding factors. Results are plotted as a function of the median age of the sliding windows from which the high- and low-PRS subgroups were extracted. HP, hippocampus; SCZ, schizophrenia; MDD, major depressive disorder; AD, Alzheimer's disease; CD, Crohn's disease; RA, rheumatoid arthritis.*

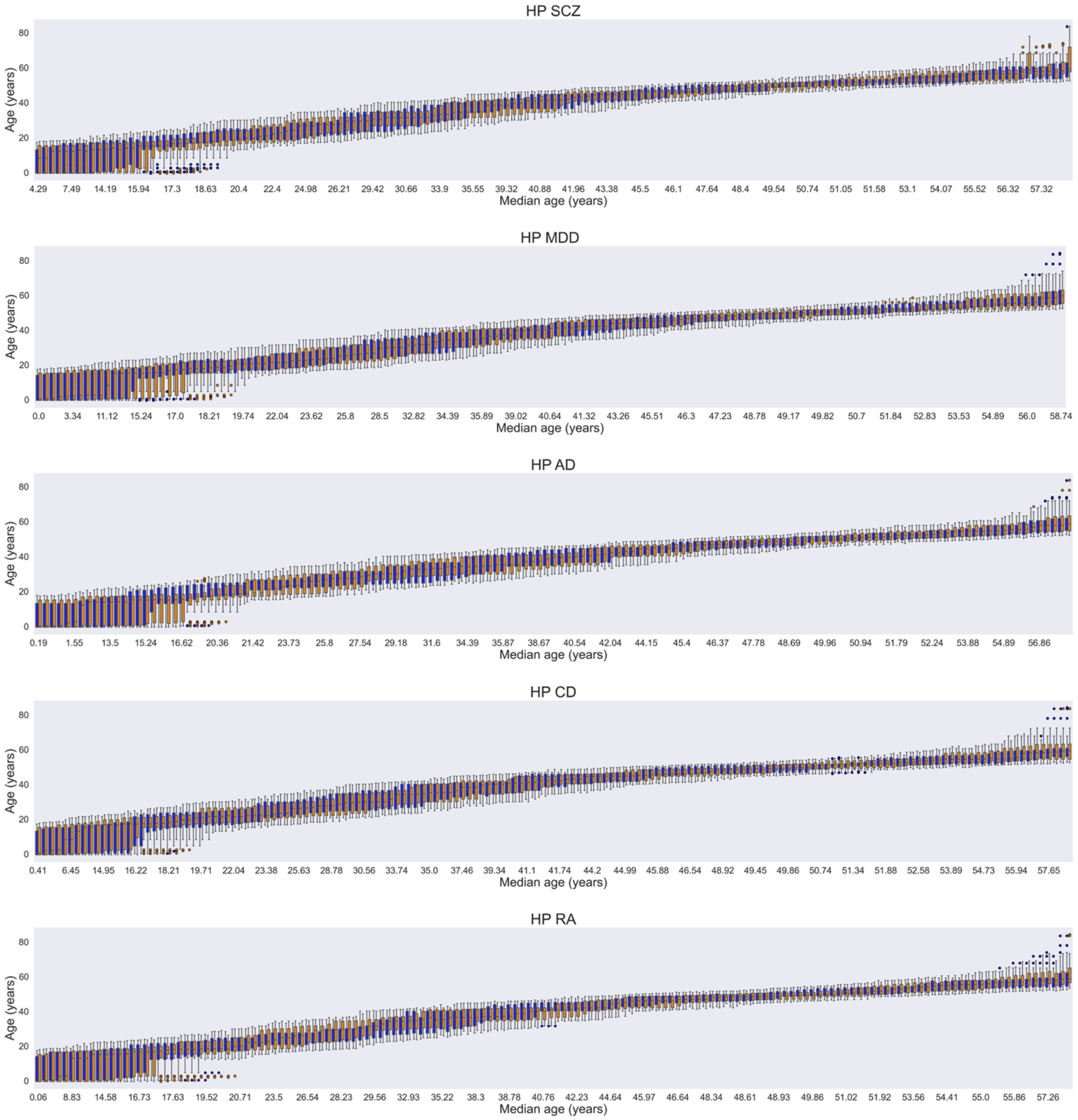


*Figure S21. Age distribution within subgroups of 25 low-PRS (blue) and 25 high-PRS (orange) subjects for the considered disorders (rows), derived from 60-subject HP sliding windows. Adjacent boxplot pairs represent low- and high-PRS cohorts originating from the same window; distributions are shown after correcting for significant differences in confounding factors. Results are plotted as a function of the median age of the sliding windows from which the high- and low-PRS subgroups were extracted. HP, hippocampus; SCZ, schizophrenia; MDD, major depressive disorder; AD, Alzheimer's disease; CD, Crohn's disease; RA, rheumatoid arthritis.*

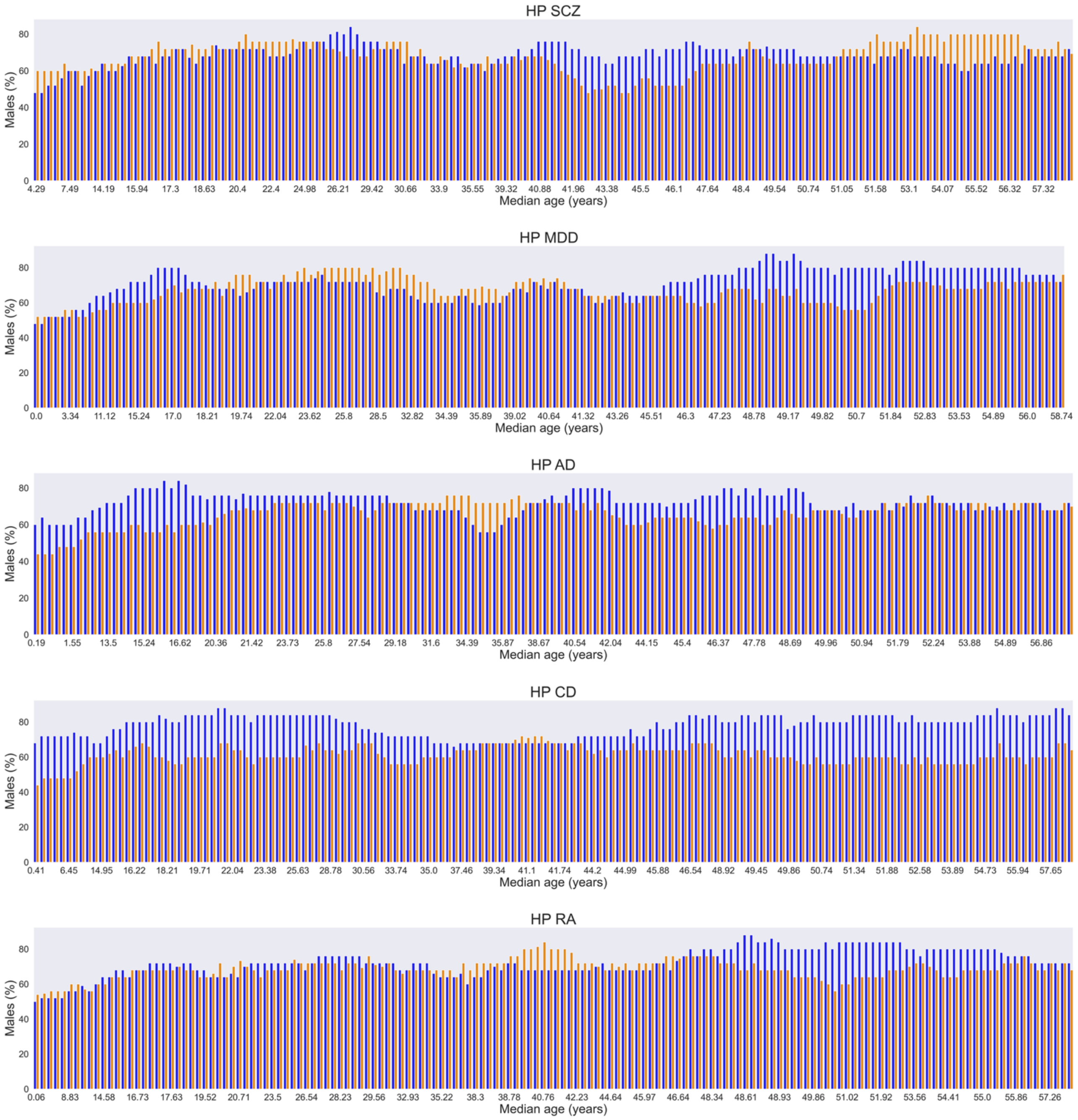


*Figure S22. Percentage of males within subgroups of 25 low-PRS (blue) and 25 high-PRS (orange) subjects for the considered disorders (rows), derived from 60-subject HP sliding windows. Adjacent bar pairs represent low- and high-PRS cohorts originating from the same window; distributions are shown after correcting for significant differences in confounding factors. Results are plotted as a function of the median age of the sliding windows from which the high- and low-PRS subgroups were extracted. HP, hippocampus; SCZ, schizophrenia; MDD, major depressive disorder; AD, Alzheimer's disease; CD, Crohn's disease; RA, rheumatoid arthritis.*

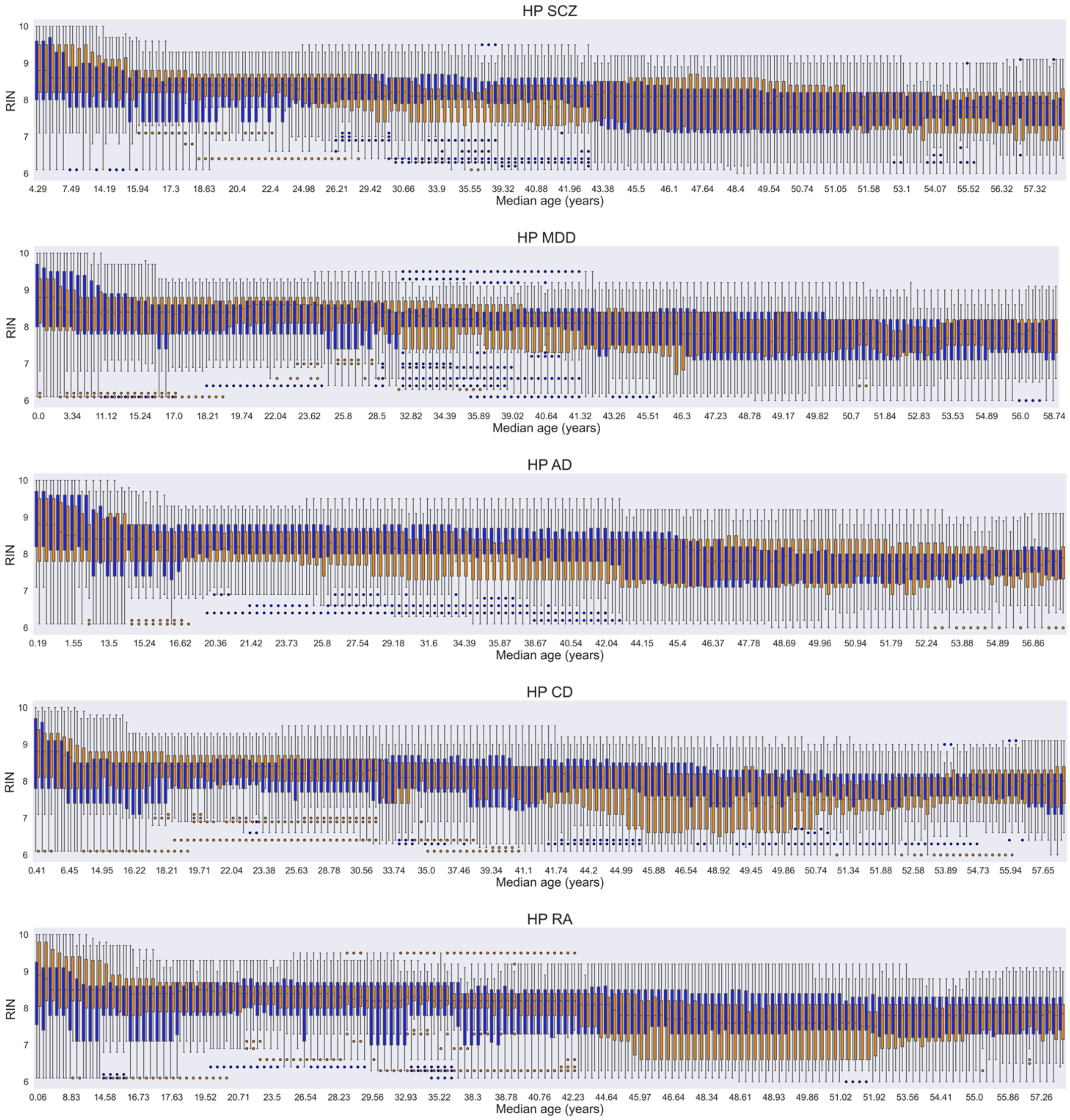


*Figure 23. RNA integrity number (RIN) distribution within subgroups of 25 low-PRS (blue) and 25 high-PRS (orange) subjects for the considered disorders (rows), derived from 60-subject HP sliding windows. Adjacent boxplot pairs represent low- and high-PRS cohorts originating from the same window; distributions are shown after correcting for significant differences in confounding factors. Results are plotted as a function of the median age of the sliding windows from which the high- and low-PRS subgroups were extracted. HP, hippocampus; SCZ, schizophrenia; MDD, major depressive disorder; AD, Alzheimer's disease; CD, Crohn's disease; RA, rheumatoid arthritis.*

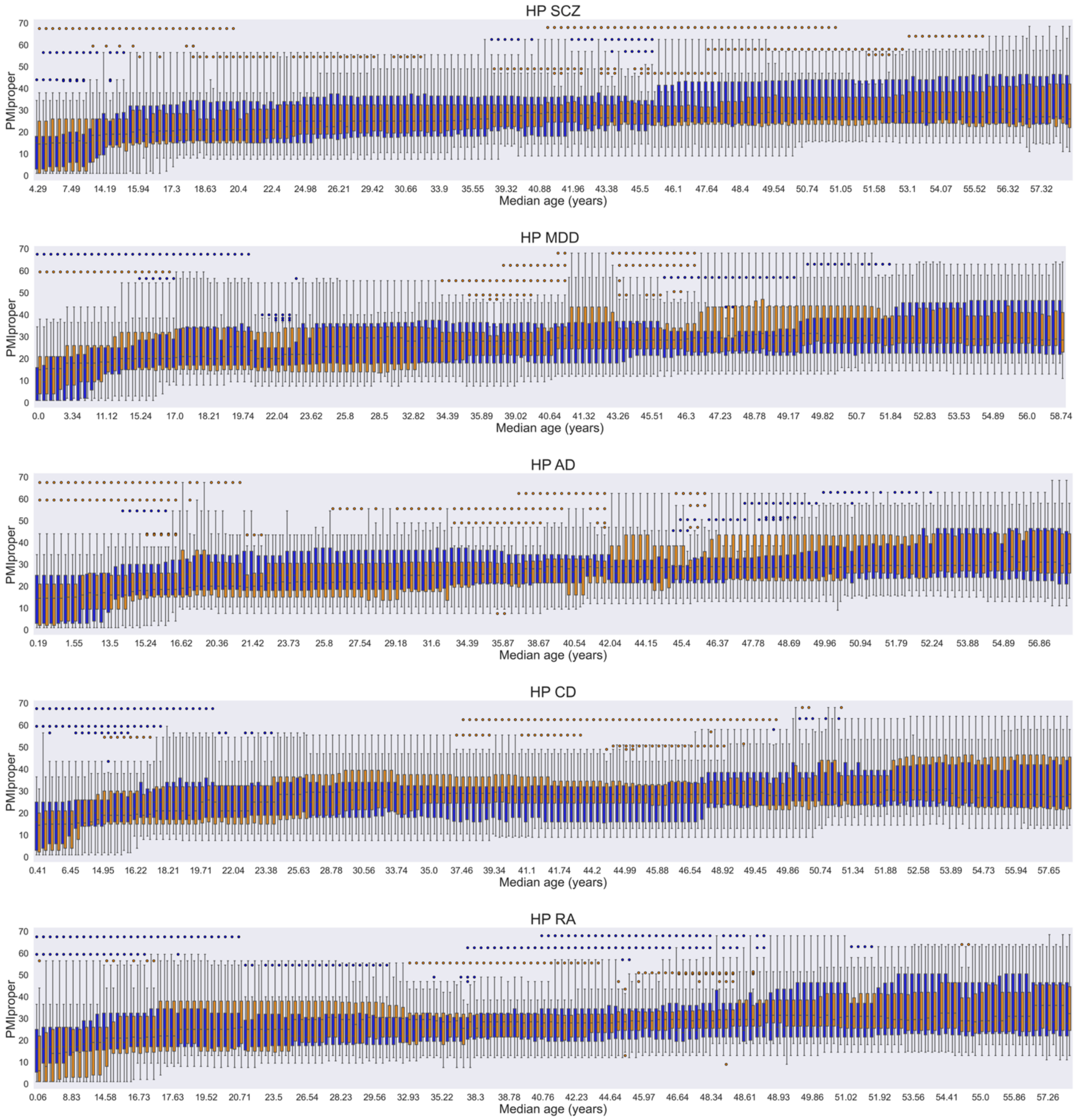


*Figure 24. Distribution of the postmortem interval variable (PMIproper) within subgroups of 25 low-PRS (blue) and 25 high-PRS (orange) subjects for the considered disorders (rows), derived from 60-subject HP sliding windows. Adjacent boxplot pairs represent low- and high-PRS cohorts originating from the same window; distributions are shown after correcting for significant differences in confounding factors. Results are plotted as a function of the median age of the sliding windows from which the high- and low-PRS subgroups were extracted. HP, hippocampus; SCZ, schizophrenia; MDD, major depressive disorder; AD, Alzheimer's disease; CD, Crohn's disease; RA, rheumatoid arthritis.*

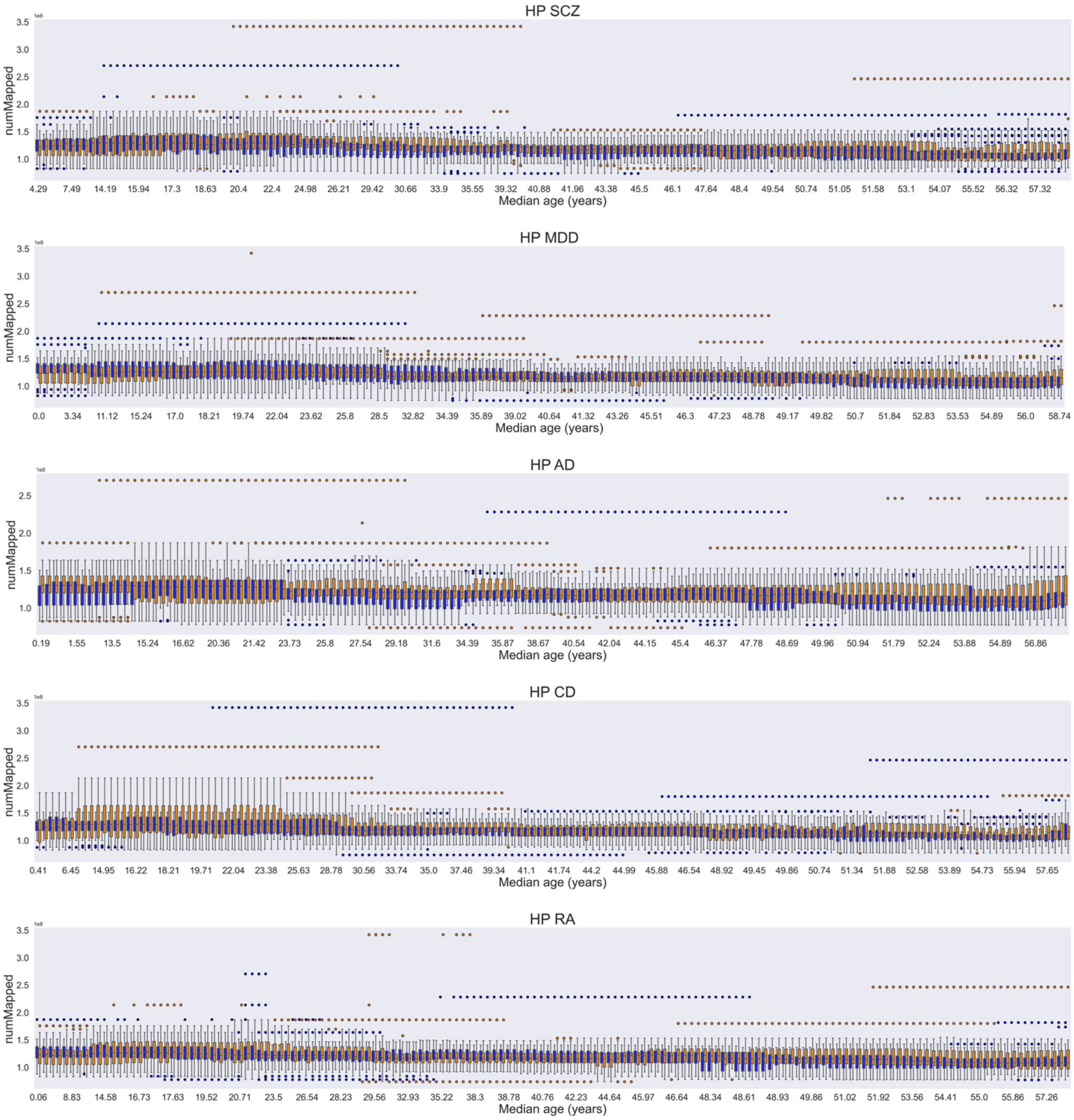


*Figure S25. Distribution of the number of reads which successfully mapped to the reference genome during alignment (numMapped) within subgroups of 25 low-PRS (blue) and 25 high-PRS (orange) subjects for the considered disorders (rows), derived from 60-subject HP sliding windows. Adjacent boxplot pairs represent low- and high-PRS cohorts originating from the same window; distributions are shown after correcting for significant differences in confounding factors. Results are plotted as a function of the median age of the sliding windows from which the high- and low-PRS subgroups were extracted. HP, hippocampus; SCZ, schizophrenia; MDD, major depressive disorder; AD, Alzheimer's disease; CD, Crohn's disease; RA, rheumatoid arthritis.*

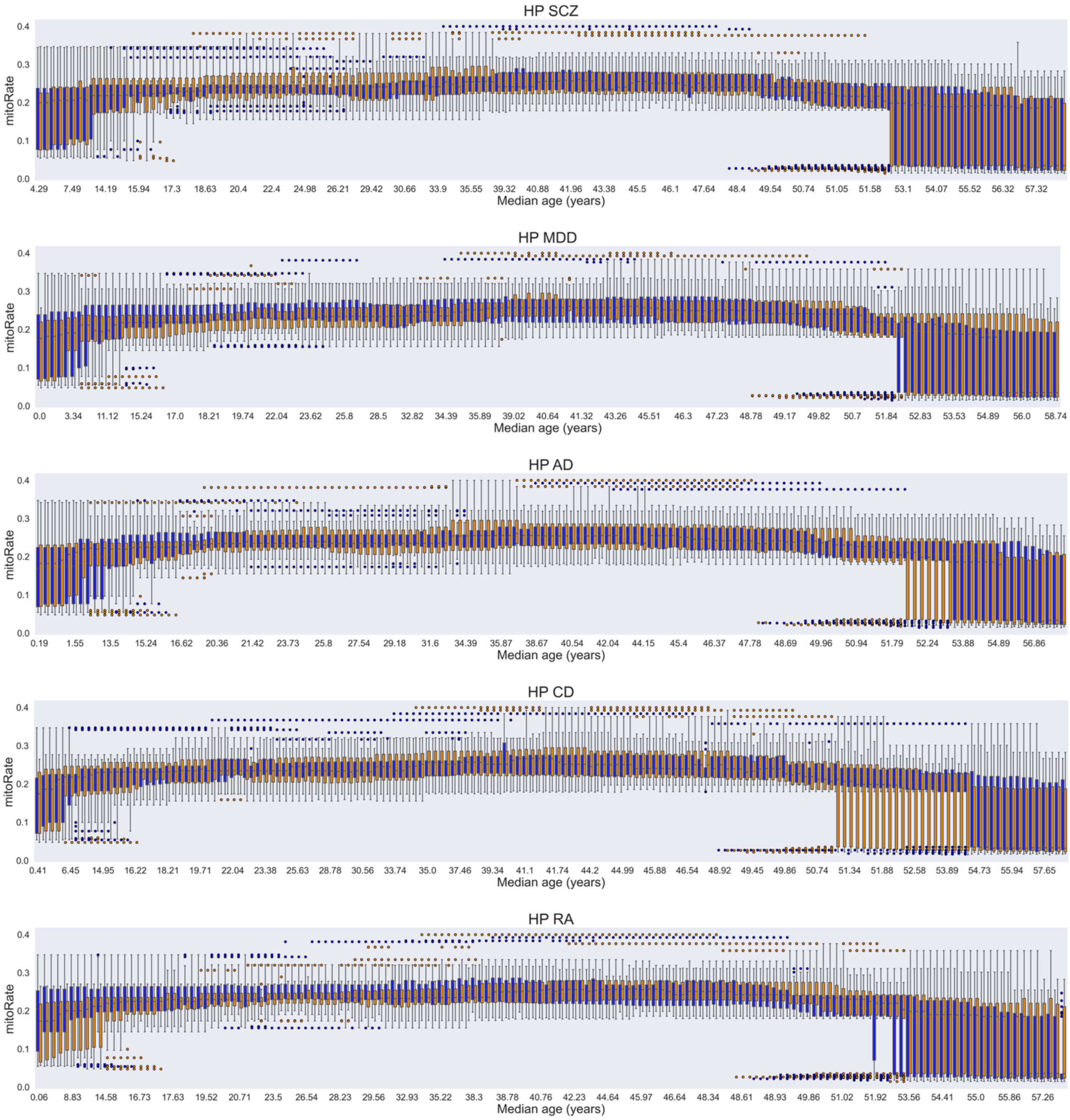


*Figure S26. Distribution of the decimal fraction of reads which mapped to the mitochondrial chromosome, of those which map at all (mitoRate), within subgroups of 25 low-PRS (blue) and 25 high-PRS (orange) subjects for the considered disorders (rows), derived from 60-subject HP sliding windows. Adjacent boxplot pairs represent low- and high-PRS cohorts originating from the same window; distributions are shown after correcting for significant differences in confounding factors. Results are plotted as a function of the median age of the sliding windows from which the high- and low-PRS subgroups were extracted. HP, hippocampus; SCZ, schizophrenia; MDD, major depressive disorder; AD, Alzheimer's disease; CD, Crohn's disease; RA, rheumatoid arthritis.*

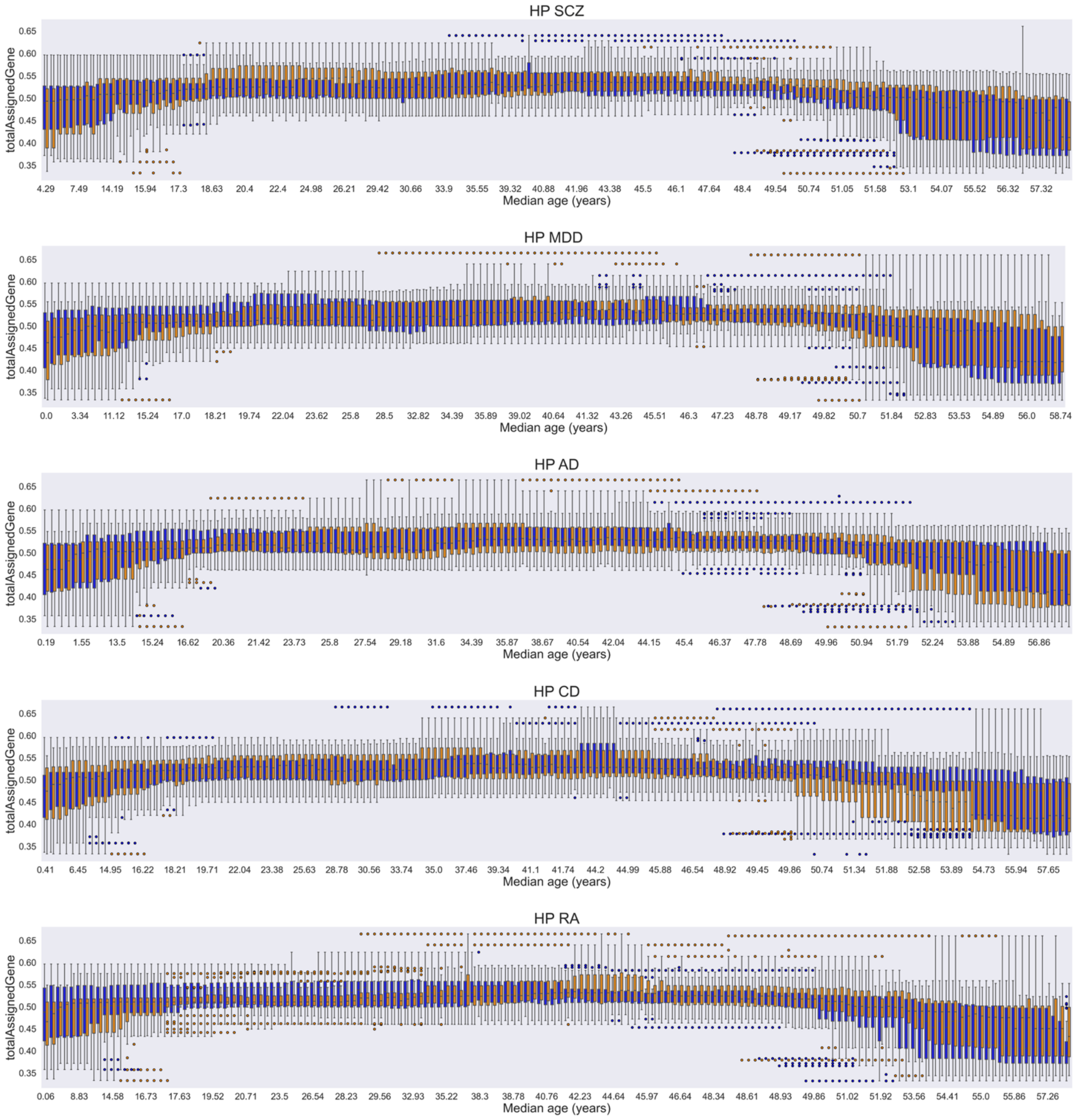


*Figure S27. Distribution of the decimal fraction of reads assigned unambiguously to a gene, with featureCounts of those in total (totalAssignedGene), within subgroups of 25 low-PRS (blue) and 25 high-PRS (orange) subjects for the considered disorders (rows), derived from 60-subject HP sliding windows. Adjacent boxplot pairs represent low- and high-PRS cohorts originating from the same window; distributions are shown after correcting for significant differences in confounding factors. Results are plotted as a function of the median age of the sliding windows from which the high- and low-PRS subgroups were extracted. HP, hippocampus; SCZ, schizophrenia; MDD, major depressive disorder; AD, Alzheimer's disease; CD, Crohn's disease; RA, rheumatoid arthritis.*

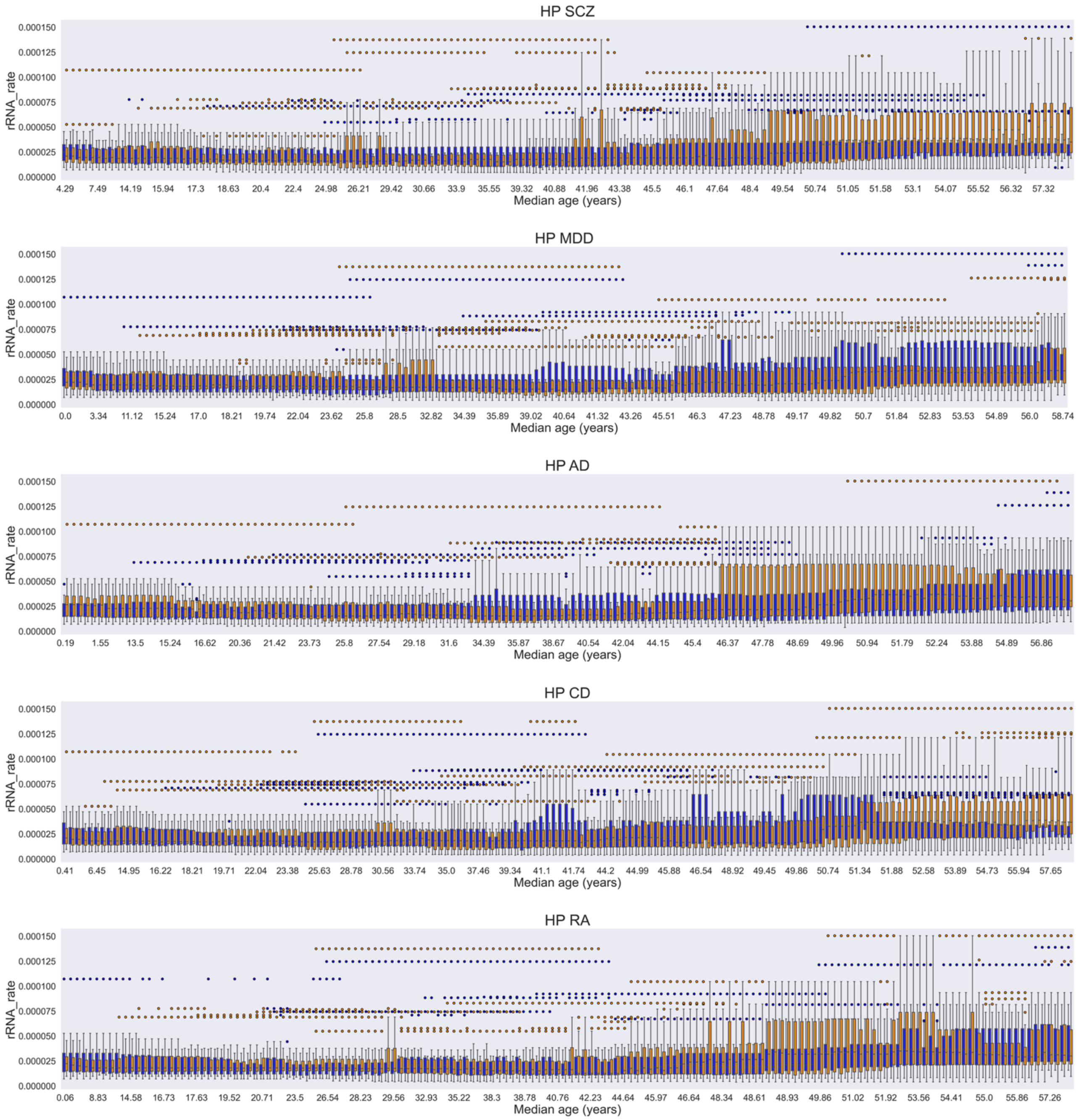


*Figure S28. Distribution of the decimal fraction of reads assigned to a gene whose type is 'rRNA', of those assigned to any gene (rRNA_rate), within subgroups of 25 low-PRS (blue) and 25 high-PRS (orange) subjects for the considered disorders (rows), derived from 60-subject HP sliding windows. Adjacent boxplot pairs represent low- and high-PRS cohorts originating from the same window; distributions are shown after correcting for significant differences in confounding factors. Results are plotted as a function of the median age of the sliding windows from which the high- and low-PRS subgroups were extracted. HP, hippocampus; SCZ, schizophrenia; MDD, major depressive disorder; AD, Alzheimer's disease; CD, Crohn's disease; RA, rheumatoid arthritis.*

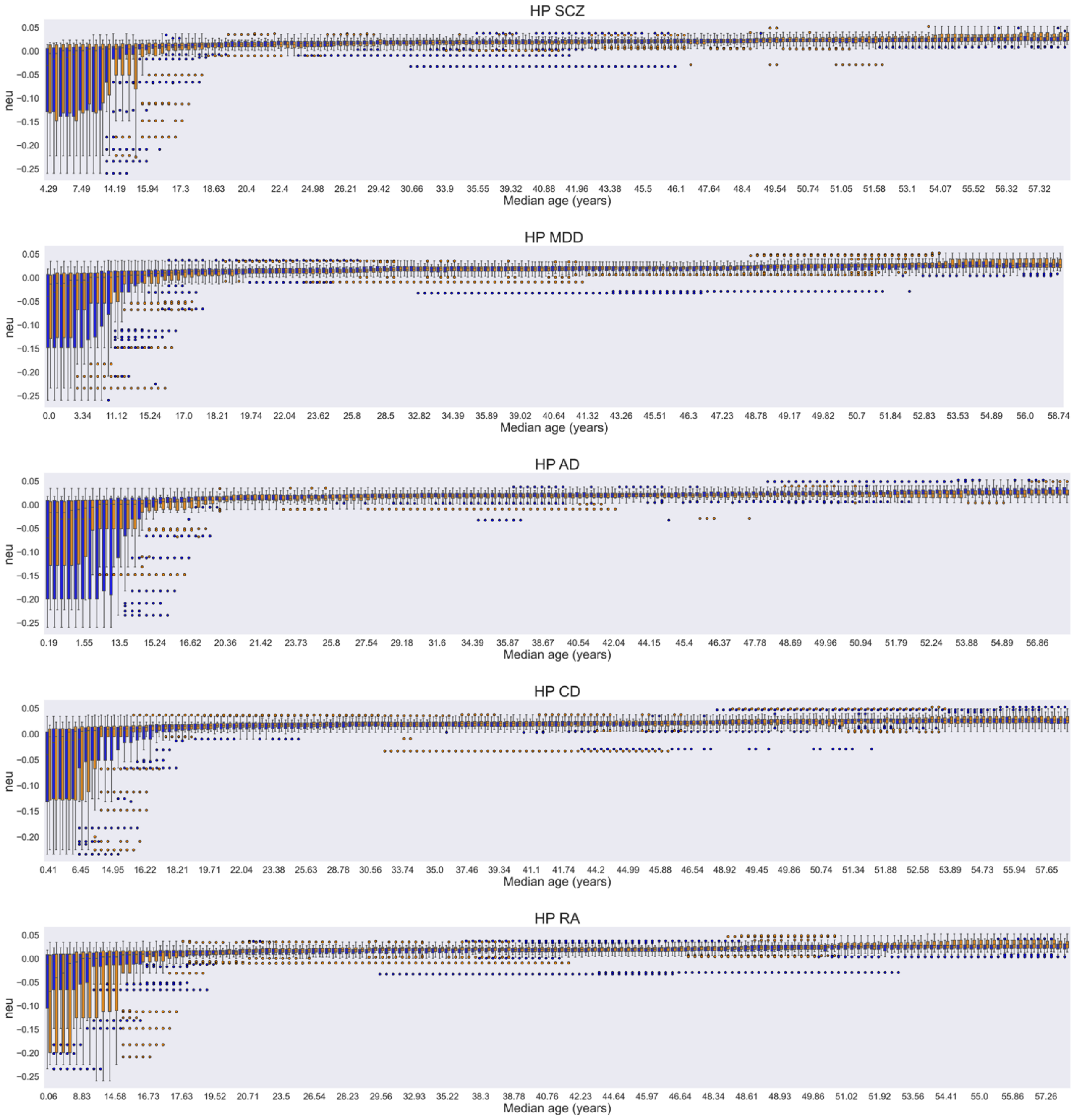


*Figure S29. Distribution of the estimated individual neuronal proportion (neu) within subgroups of 25 low-PRS (blue) and 25 high-PRS (orange) subjects for the considered disorders (rows), derived from 60-subject HP sliding windows. Adjacent boxplot pairs represent low- and high-PRS cohorts originating from the same window; distributions are shown after correcting for significant differences in confounding factors. Results are plotted as a function of the median age of the sliding windows from which the high- and low-PRS subgroups were extracted. HP, hippocampus; SCZ, schizophrenia; MDD, major depressive disorder; AD, Alzheimer's disease; CD, Crohn's disease; RA, rheumatoid arthritis.*

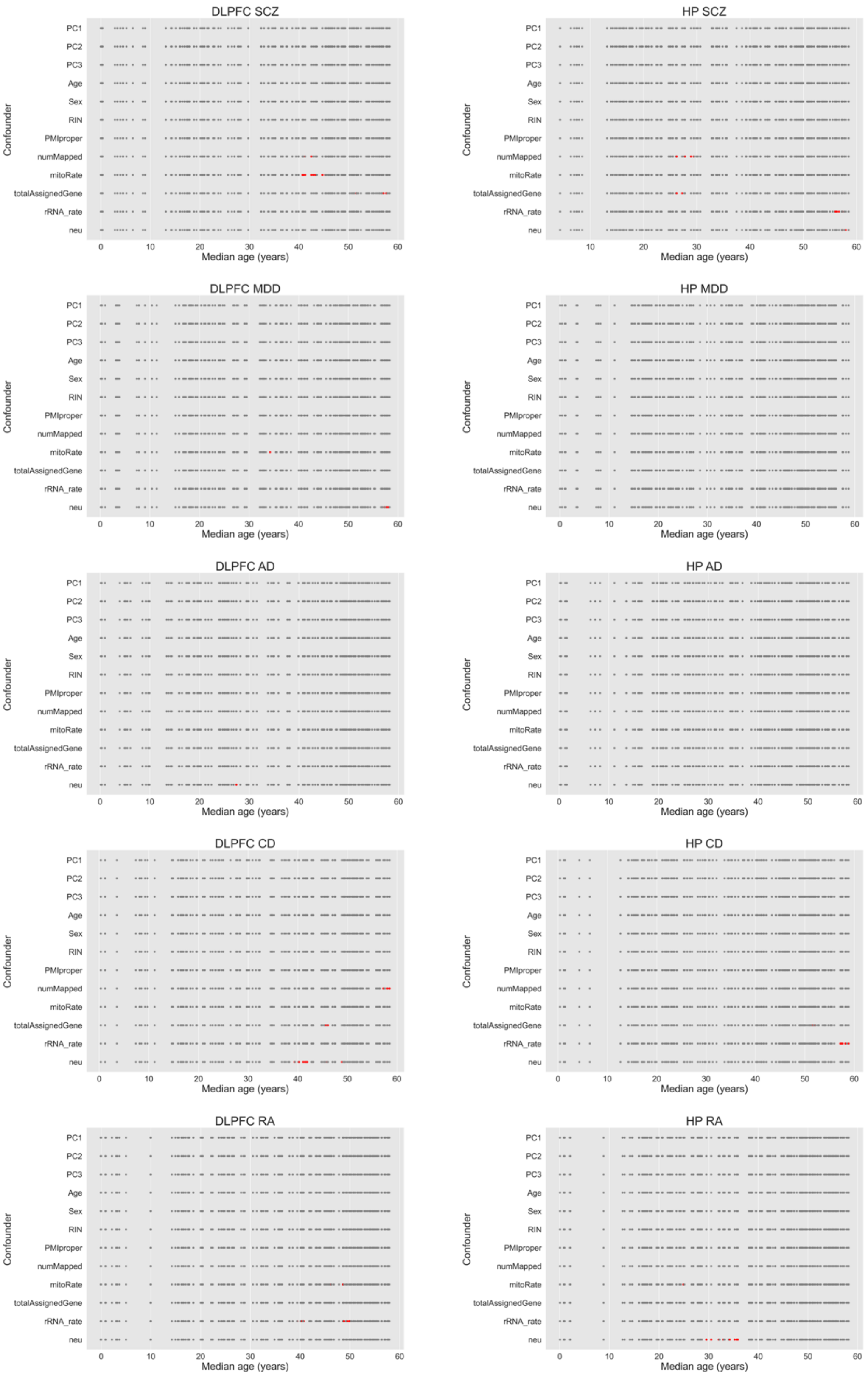


*Figure S30. Summary of residual significant differences in confounding factor distributions between high- and low-PRS subgroups across various tissue-disorder combinations. Each data point represents a sliding window corrected for significant differences between risk groups. Significance is expressed via p-values obtained from Wilcoxon rank-sum tests for continuous confounders and z-score tests for the discrete variable Sex. Gray dots indicate windows where the difference between the high- and low-PRS distributions of a given confounding factor is not significant ($p \geq 0.05$), while windows where residual significant differences ($p < 0.05$) persist after correction are highlighted in red. Results are plotted as a function of the median age of the sliding windows from which the high- and low-PRS subgroups were extracted. DLPFC, dorsolateral prefrontal cortex; HP, hippocampus; SCZ, schizophrenia; MDD, major depressive disorder; AD, Alzheimer's disease; CD, Crohn's disease; RA, rheumatoid arthritis.*

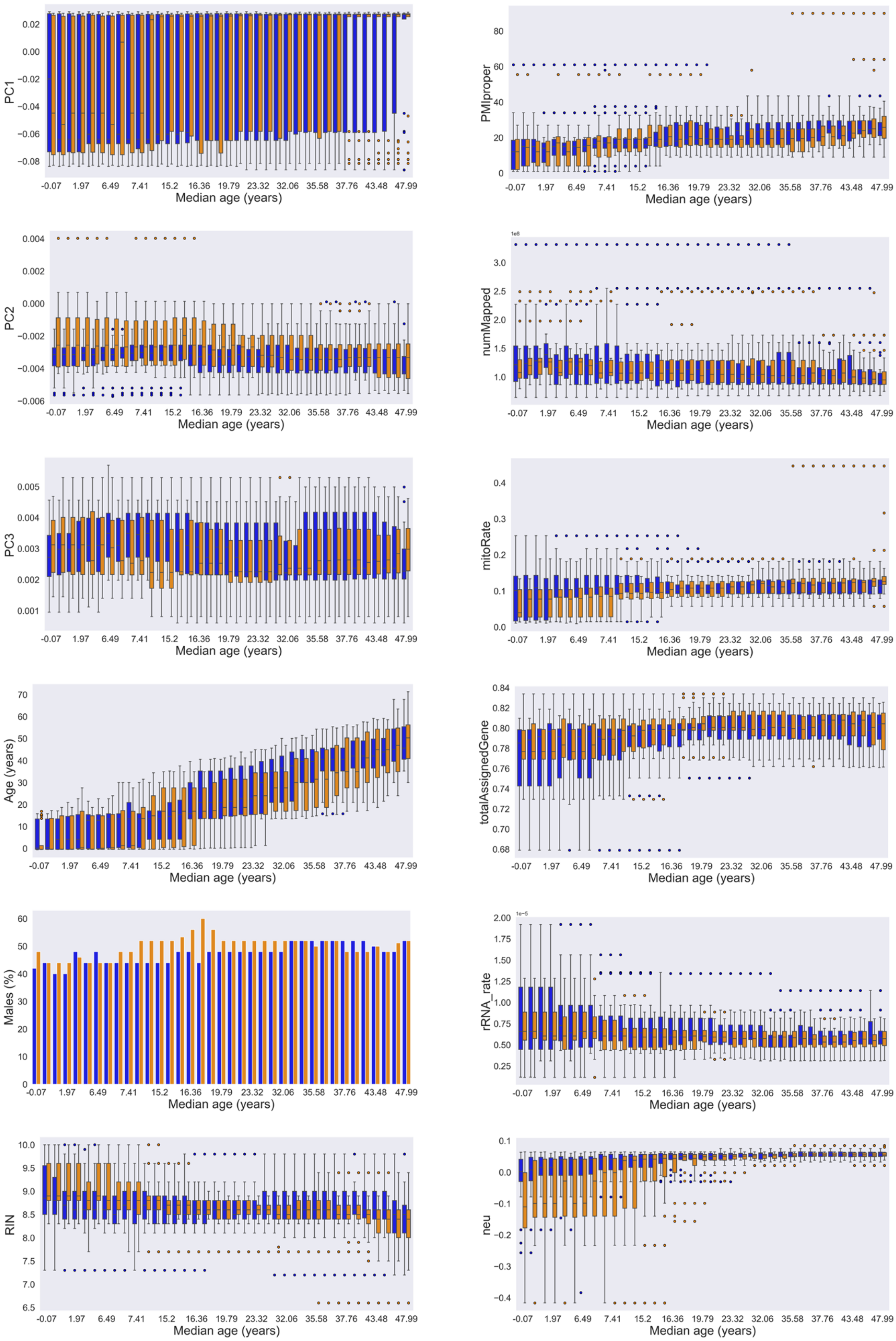


*Figure S31. Distributions of confounding factors within subgroups of 17 low-PRS (blue) and 17 high-PRS (orange) subjects for SCZ, derived from 40-subject DLPFC sliding windows in the Poly(A) replication dataset. In each panel, representing a specific confounding factor, adjacent pairs of distributions indicate low- and high-PRS cohorts originating from the same window; results are shown after correcting for significant differences in confounding factors. Distributions are plotted as a function of the median age of the sliding windows from which the high- and low-PRS subgroups were extracted.*

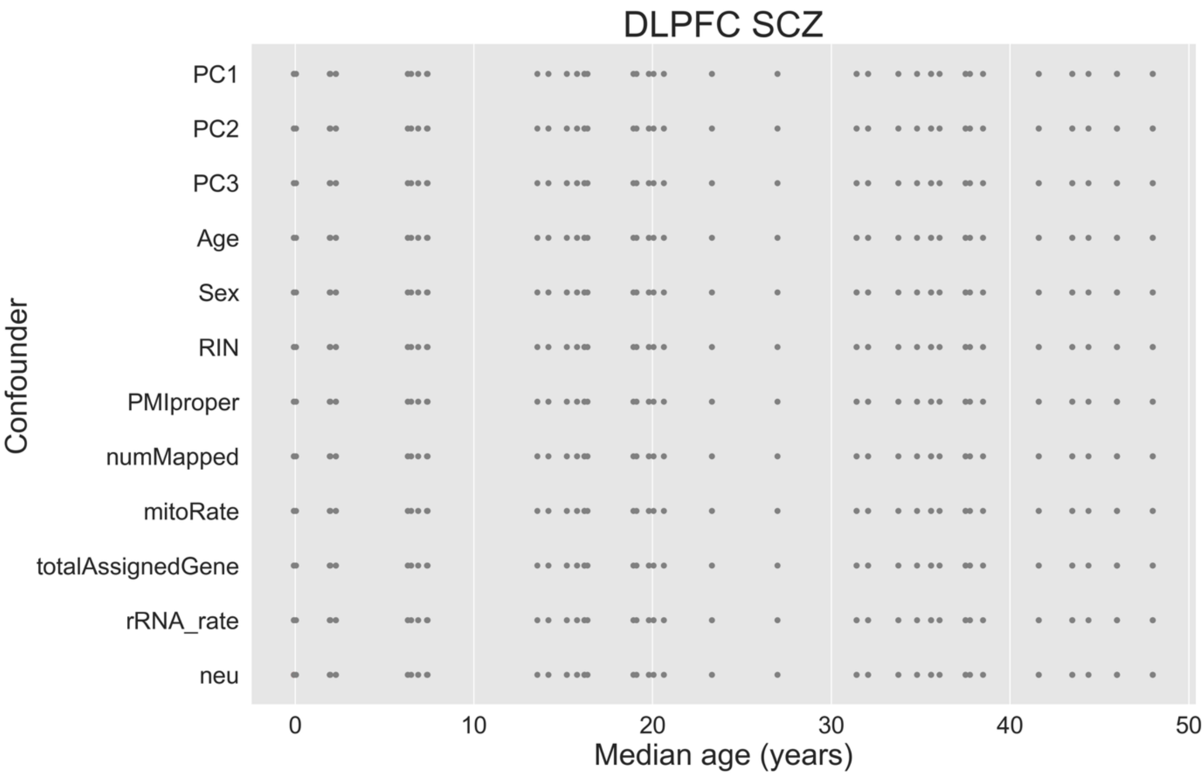


*Figure S32. Statistical comparison of confounding factor distributions between high- and low-PRS subgroups in the Poly(A) replication dataset for the DLPFC-SCZ case. Each data point represents a sliding window corrected for significant differences between risk groups. Significance is expressed via p-values obtained from Wilcoxon rank-sum tests for continuous confounders and z-score tests for the discrete variable Sex. Following the correction procedure, no residual significant differences ($p < 0.05$) persist across any of the windows; consequently, all data points are represented as gray dots, indicating non-significant differences ($p \geq 0.05$) between the high- and low-PRS distributions. Results are plotted as a function of the median age of the sliding windows from which the high- and low-PRS subgroups were extracted. DLPFC, dorsolateral prefrontal cortex; SCZ, schizophrenia.*

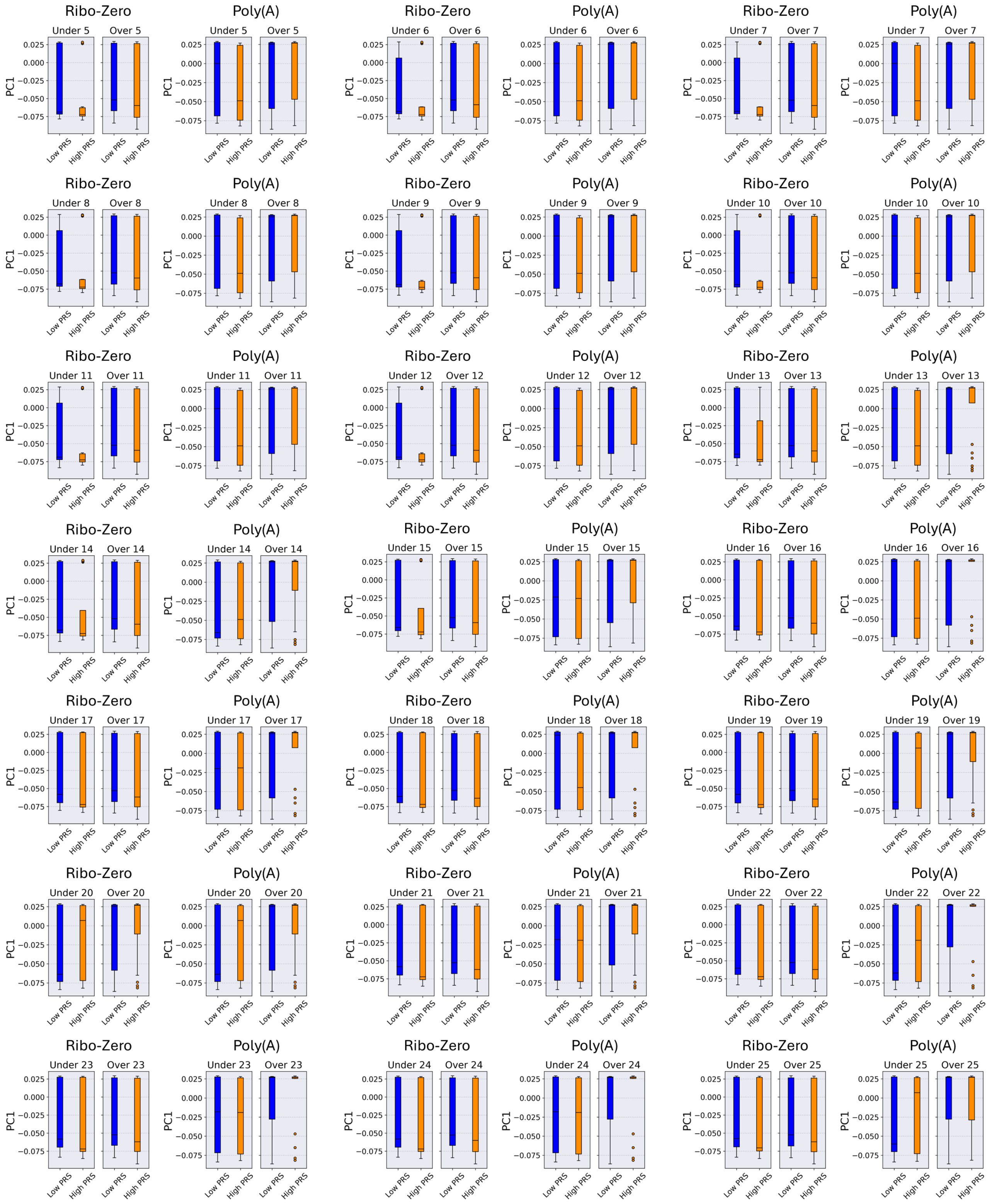


*Figure S33. Distributions of the first genomic eigenvariate (PC1) for low-PRS (blue) and high-PRS (orange) cohorts, grouped into the "Under" and "Over" age ranges for both Ribo-Zero and Poly(A) datasets. Results are displayed in separate panels for each age threshold Y in the 5—25 juvenile range; within each panel, data are organized by dataset and age group, and adjacent pairs of boxplots represent the corresponding low-PRS and high-PRS cohorts. Distributions are shown after correcting for significant differences in confounding factors between pairs of high-PRS and low-PRS cohorts originating from the same dataset and age group.*

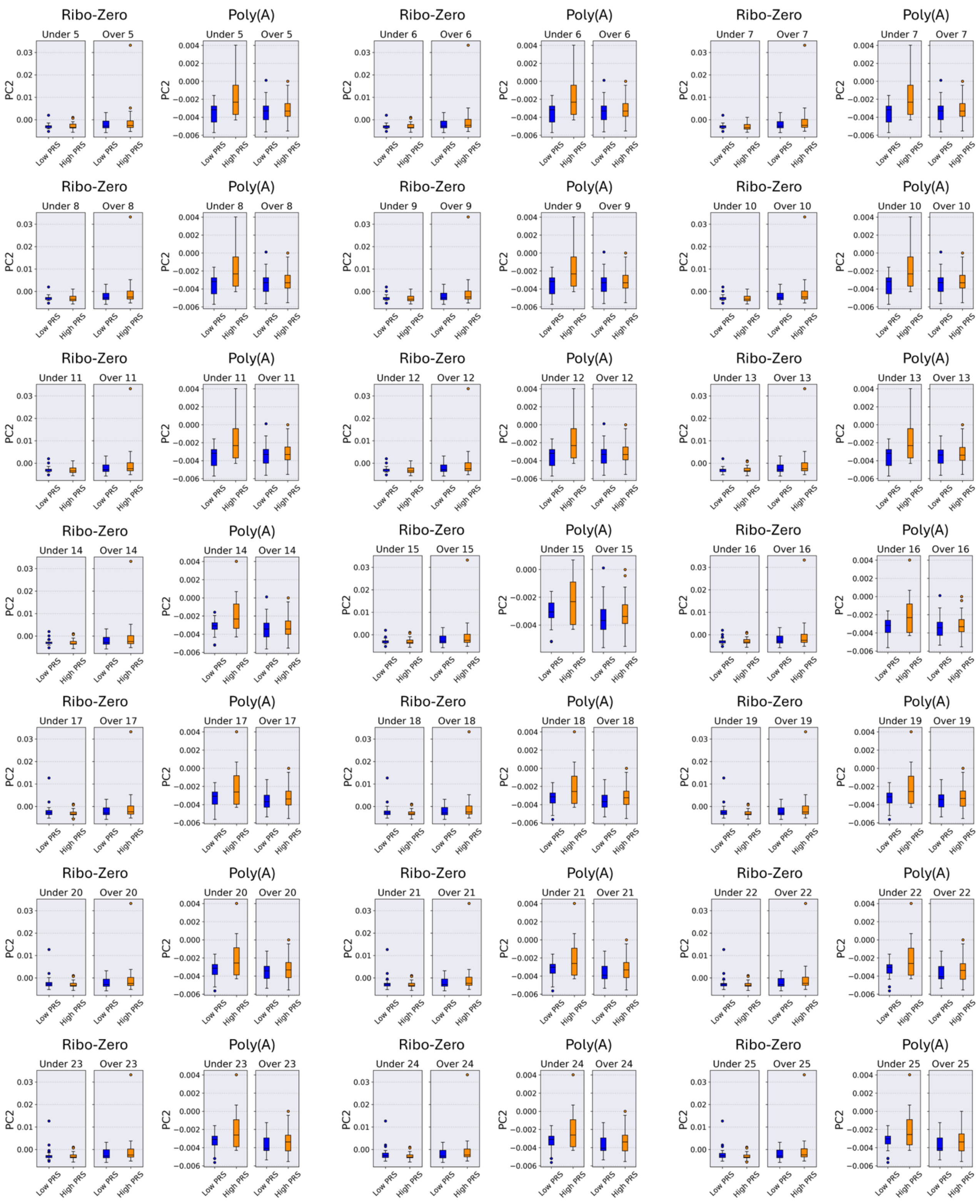

*Figure S34. Distributions of the second genomic eigenvariate (PC2) for low-PRS (blue) and high-PRS (orange) cohorts, grouped into the "Under" and "Over" age ranges for both Ribo-Zero and Poly(A) datasets. Results are displayed in separate panels for each age threshold Y in the 5—25 juvenile range; within each panel, data are organized by dataset and age group, and adjacent pairs of boxplots represent the corresponding low-PRS and high-PRS cohorts. Distributions are shown after correcting for significant differences in confounding factors between pairs of high-PRS and low-PRS cohorts originating from the same dataset and age group.*

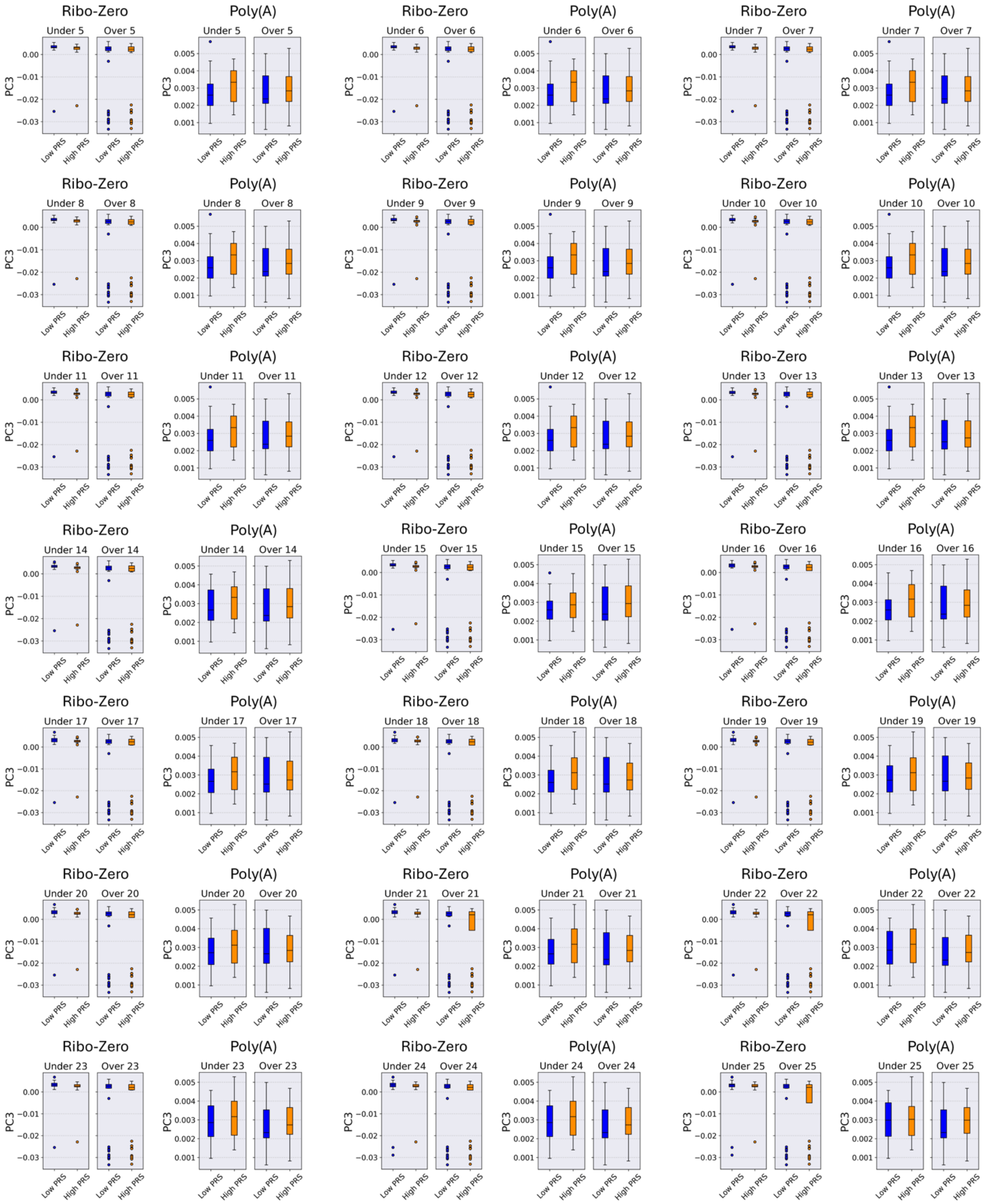


*Figure S35. Distributions of the third genomic eigenvariate (PC3) for low-PRS (blue) and high-PRS (orange) cohorts, grouped into the "Under" and "Over" age ranges for both Ribo-Zero and Poly(A) datasets. Results are displayed in separate panels for each age threshold $Y$ in the 5—25 juvenile range; within each panel, data are organized by dataset and age group, and adjacent pairs of boxplots represent the corresponding low-PRS and high-PRS cohorts. Distributions are shown after correcting for significant differences in confounding factors between pairs of high-PRS and low-PRS cohorts originating from the same dataset and age group.*

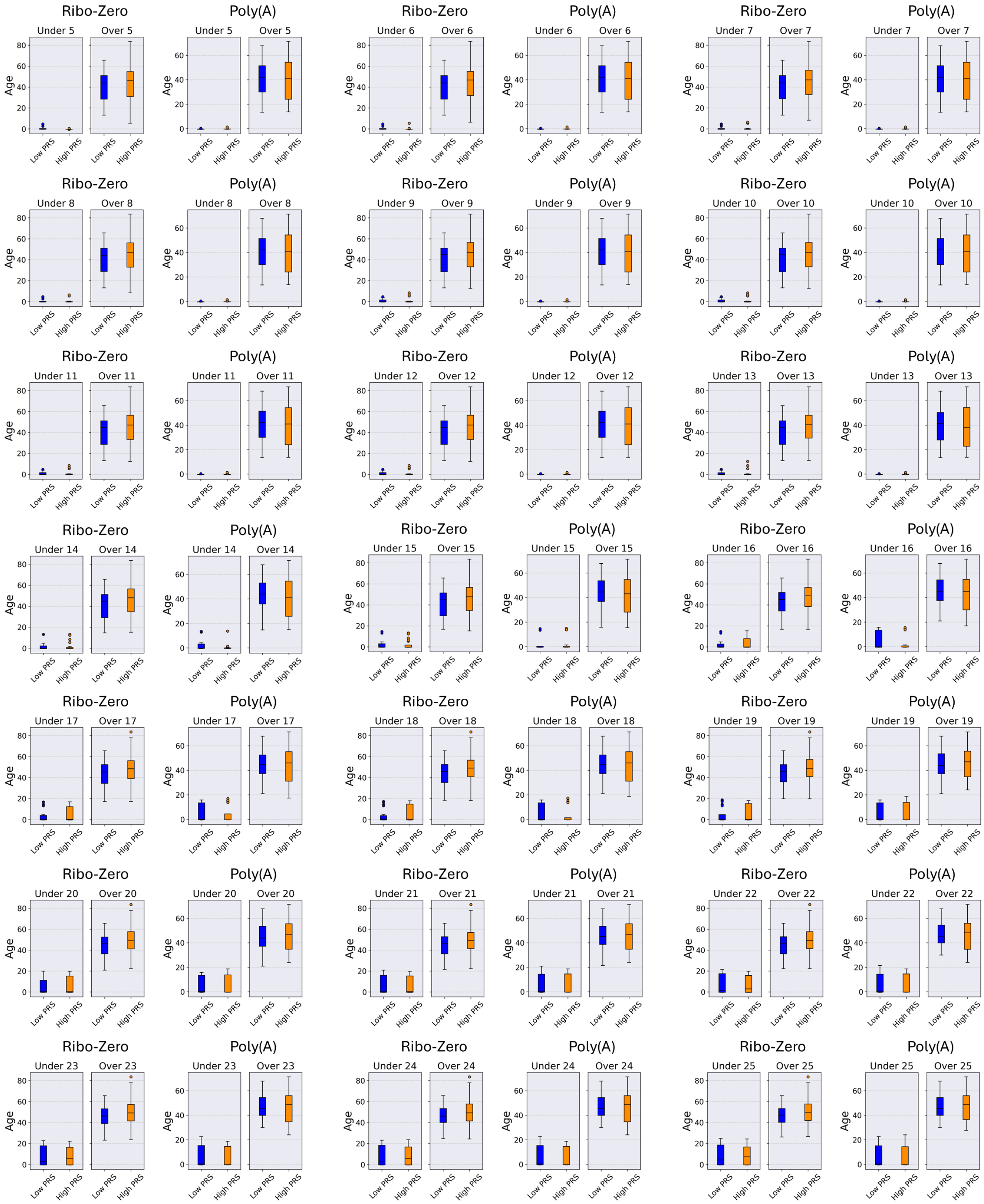

*Figure S36. Age distributions for low-PRS (blue) and high-PRS (orange) cohorts, grouped into the "Under" and "Over" age ranges for both Ribo-Zero and Poly(A) datasets. Results are displayed in separate panels for each age threshold Y in the 5—25 juvenile range; within each panel, data are organized by dataset and age group, and adjacent pairs of boxplots represent the corresponding low-PRS and high-PRS cohorts. Distributions are shown after correcting for significant differences in confounding factors between pairs of high-PRS and low-PRS cohorts originating from the same dataset and age group.*

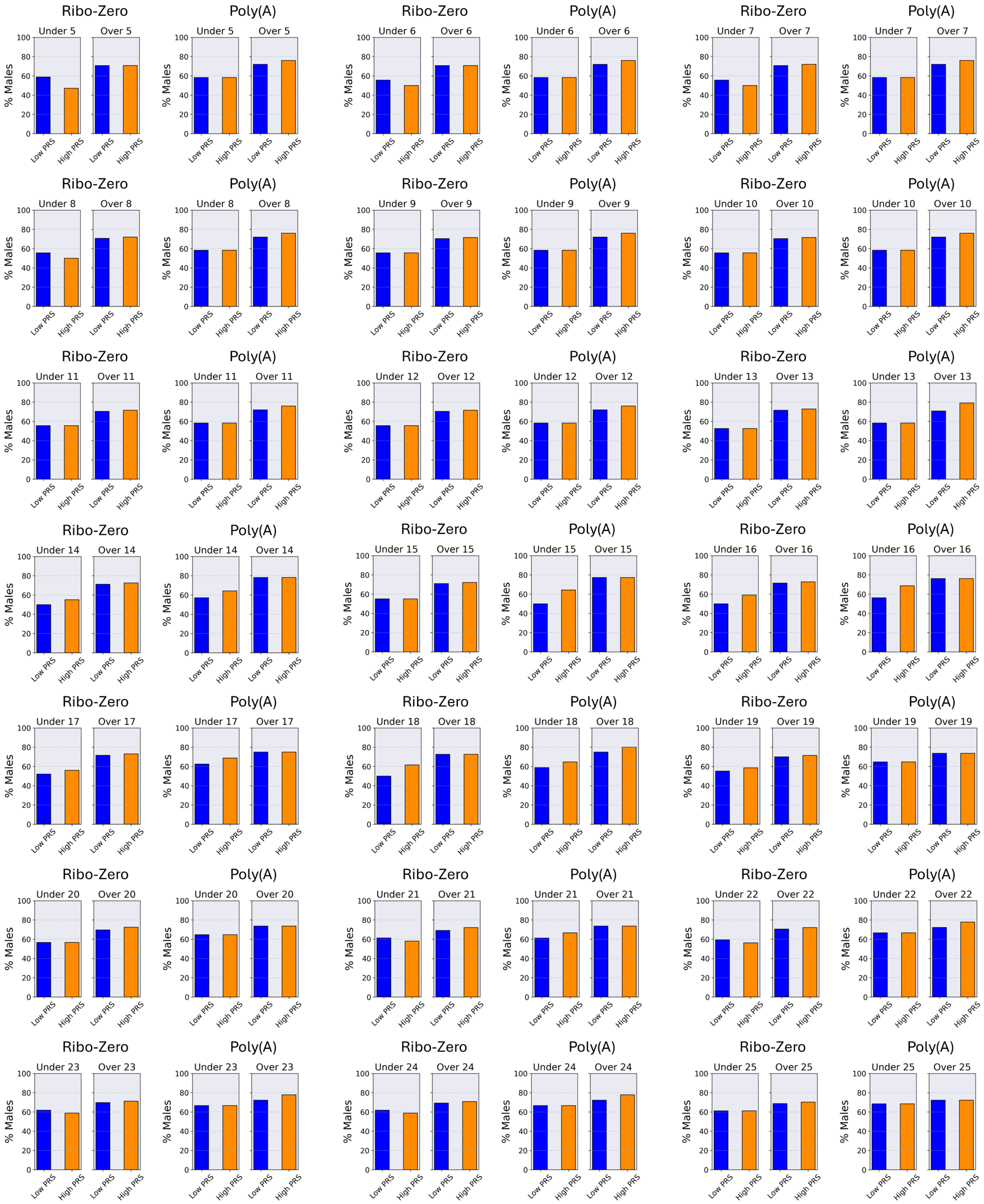

*Figure S37. Percentage of males in low-PRS (blue) and high-PRS (orange) cohorts, grouped into the "Under" and "Over" age ranges for both Ribo-Zero and Poly(A) datasets. Results are displayed in separate panels for each age threshold Y in the 5—25 juvenile range; within each panel, data are organized by dataset and age group, and adjacent bar pairs represent the corresponding low-PRS and high-PRS cohorts. Distributions are shown after correcting for significant differences in confounding factors between pairs of high-PRS and low-PRS cohorts originating from the same dataset and age group.*

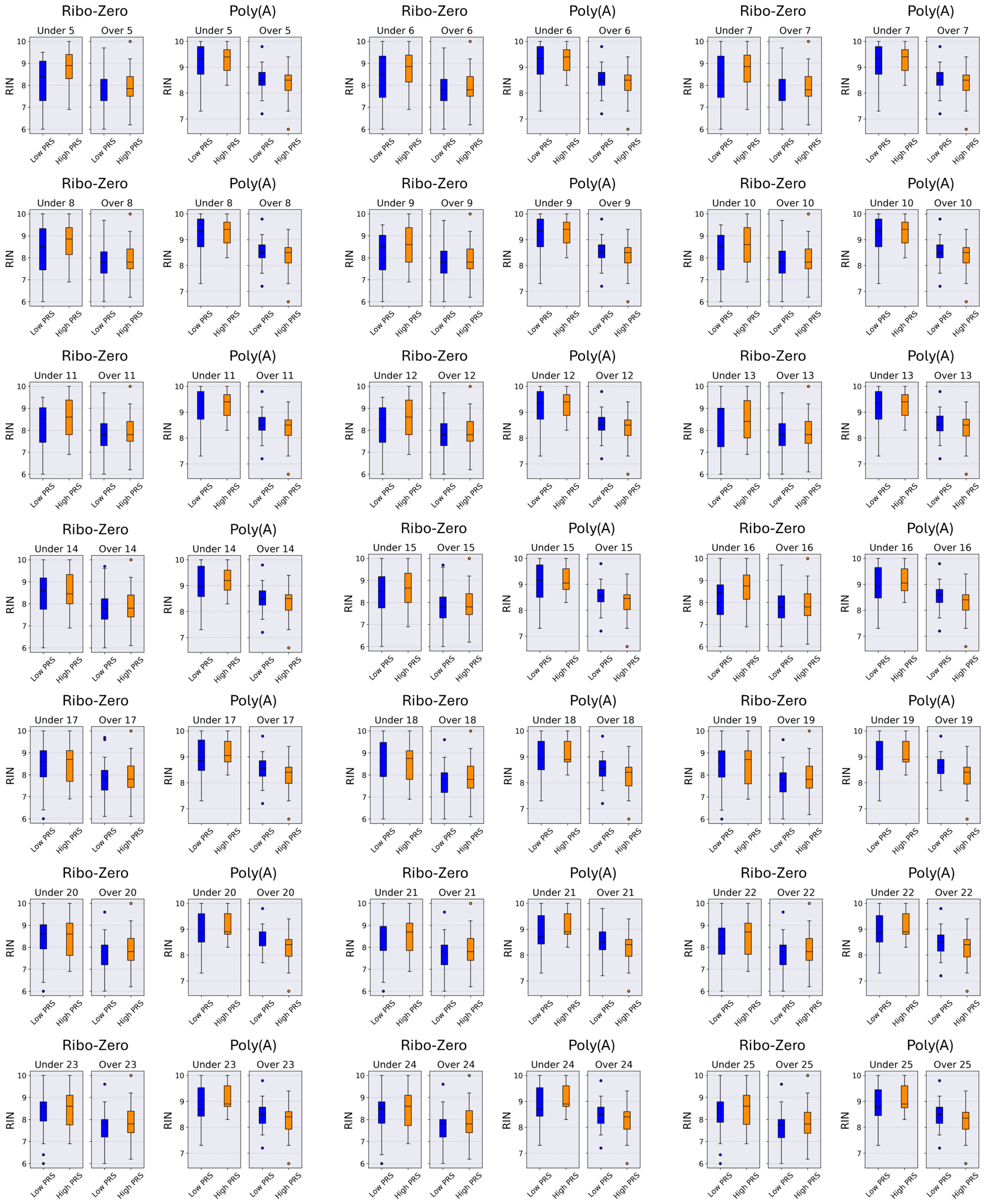


*Figure S38. RNA integrity number (RIN) distributions for low-PRS (blue) and high-PRS (orange) cohorts, grouped into the "Under" and "Over" age ranges for both Ribo-Zero and Poly(A) datasets. Results are displayed in separate panels for each age threshold Y in the 5—25 juvenile range; within each panel, data are organized by dataset and age group, and adjacent pairs of boxplots represent the corresponding low-PRS and high-PRS cohorts. Distributions are shown after correcting for significant differences in confounding factors between pairs of high-PRS and low-PRS cohorts originating from the same dataset and age group.*

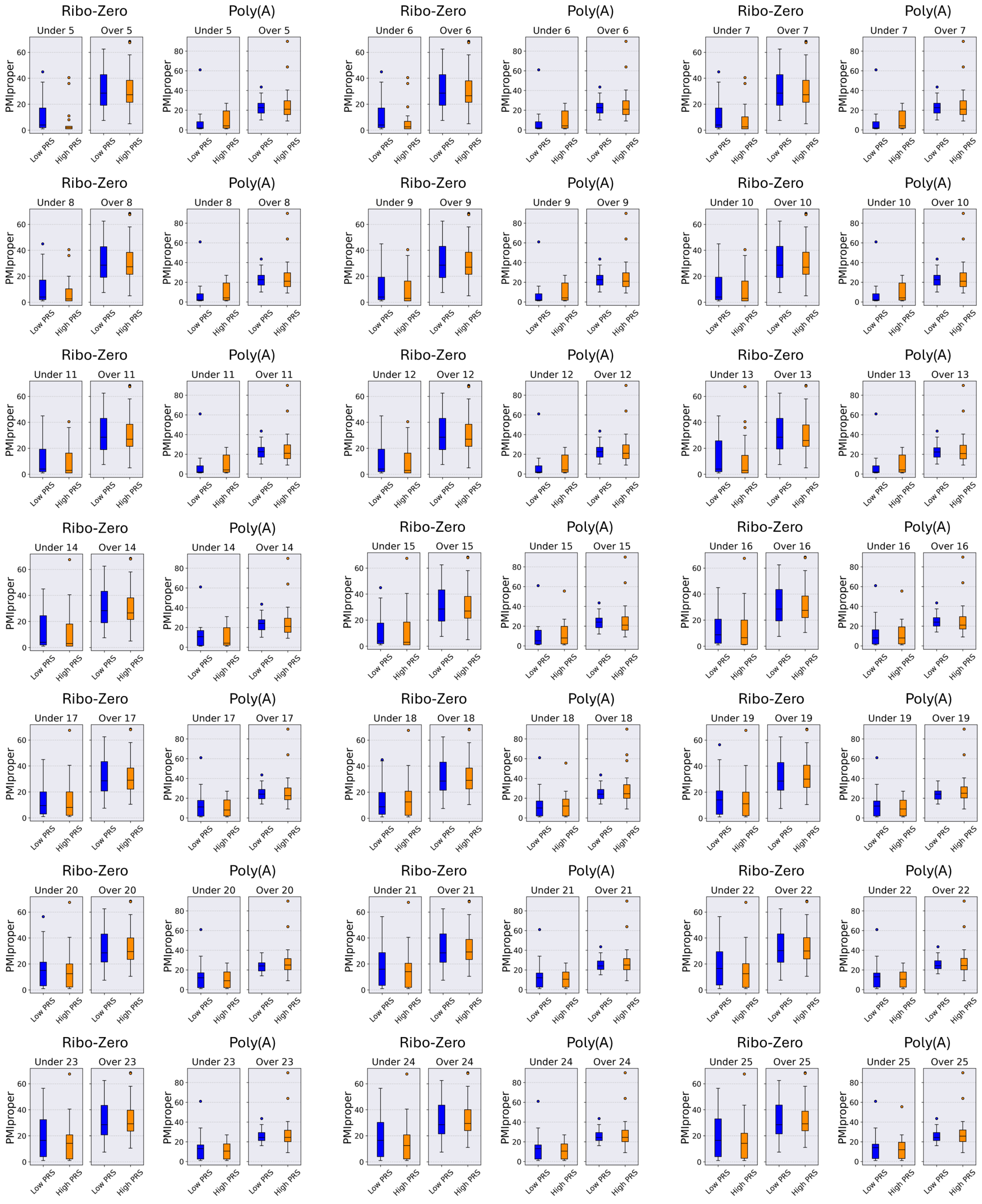


*Figure S39. Distribution of the postmortem interval variable (PMIproper) for low-PRS (blue) and high-PRS (orange) cohorts, grouped into the "Under" and "Over" age ranges for both Ribo-Zero and Poly(A) datasets. Results are displayed in separate panels for each age threshold Y in the 5—25 juvenile range; within each panel, data are organized by dataset and age group, and adjacent pairs of boxplots represent the corresponding low-PRS and high-PRS cohorts. Distributions are shown after correcting for significant differences in confounding factors between pairs of high-PRS and low-PRS cohorts originating from the same dataset and age group.*

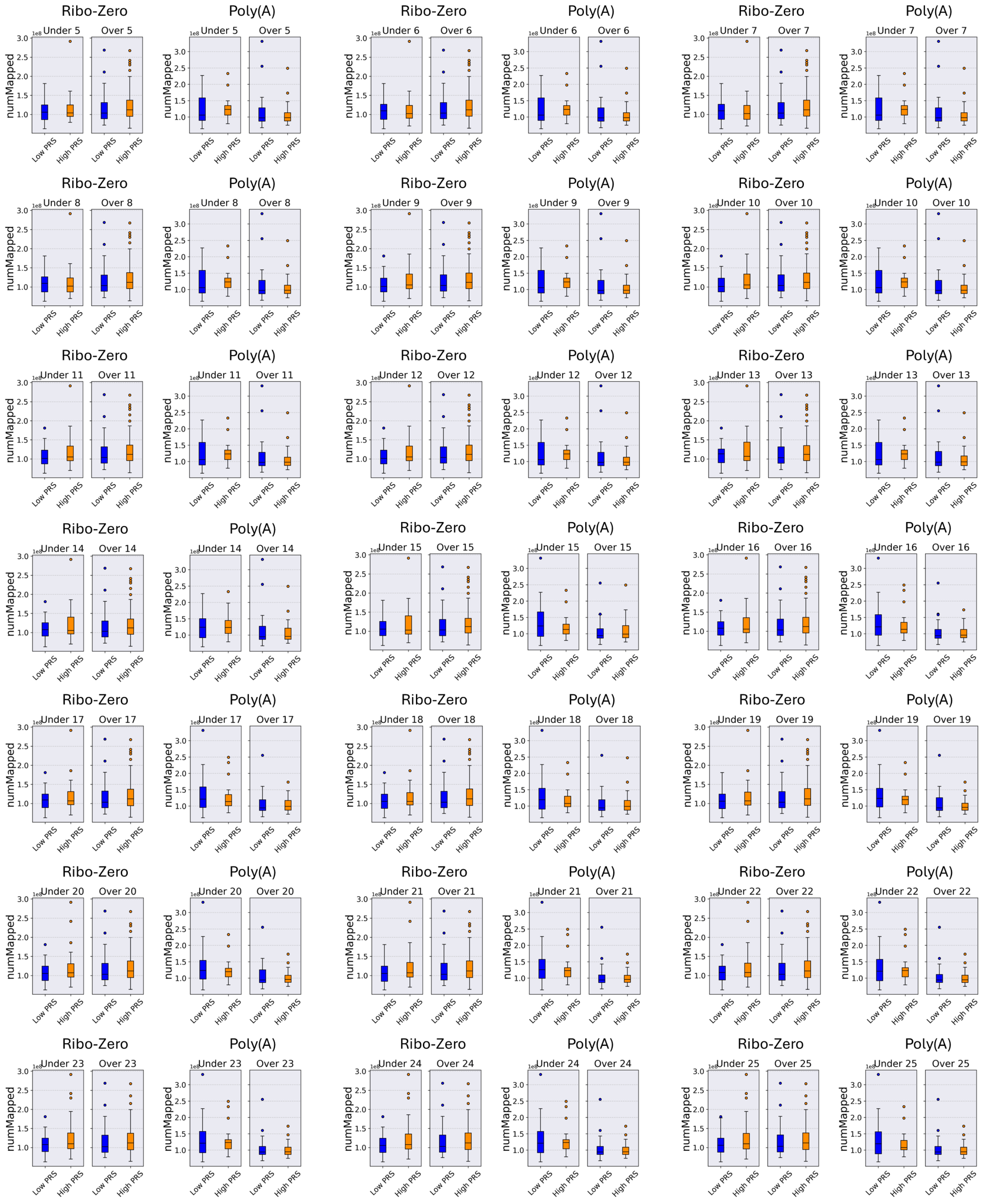


*Figure S40. Distribution of the number of reads which successfully mapped to the reference genome during alignment (numMapped) for low-PRS (blue) and high-PRS (orange) cohorts, grouped into the "Under" and "Over" age ranges for both Ribo-Zero and Poly(A) datasets. Results are displayed in separate panels for each age threshold $Y$ in the 5—25 juvenile range; within each panel, data are organized by dataset and age group, and adjacent pairs of boxplots represent the corresponding low-PRS and high-PRS cohorts. Distributions are shown after correcting for significant differences in confounding factors between pairs of high-PRS and low-PRS cohorts originating from the same dataset and age group.*

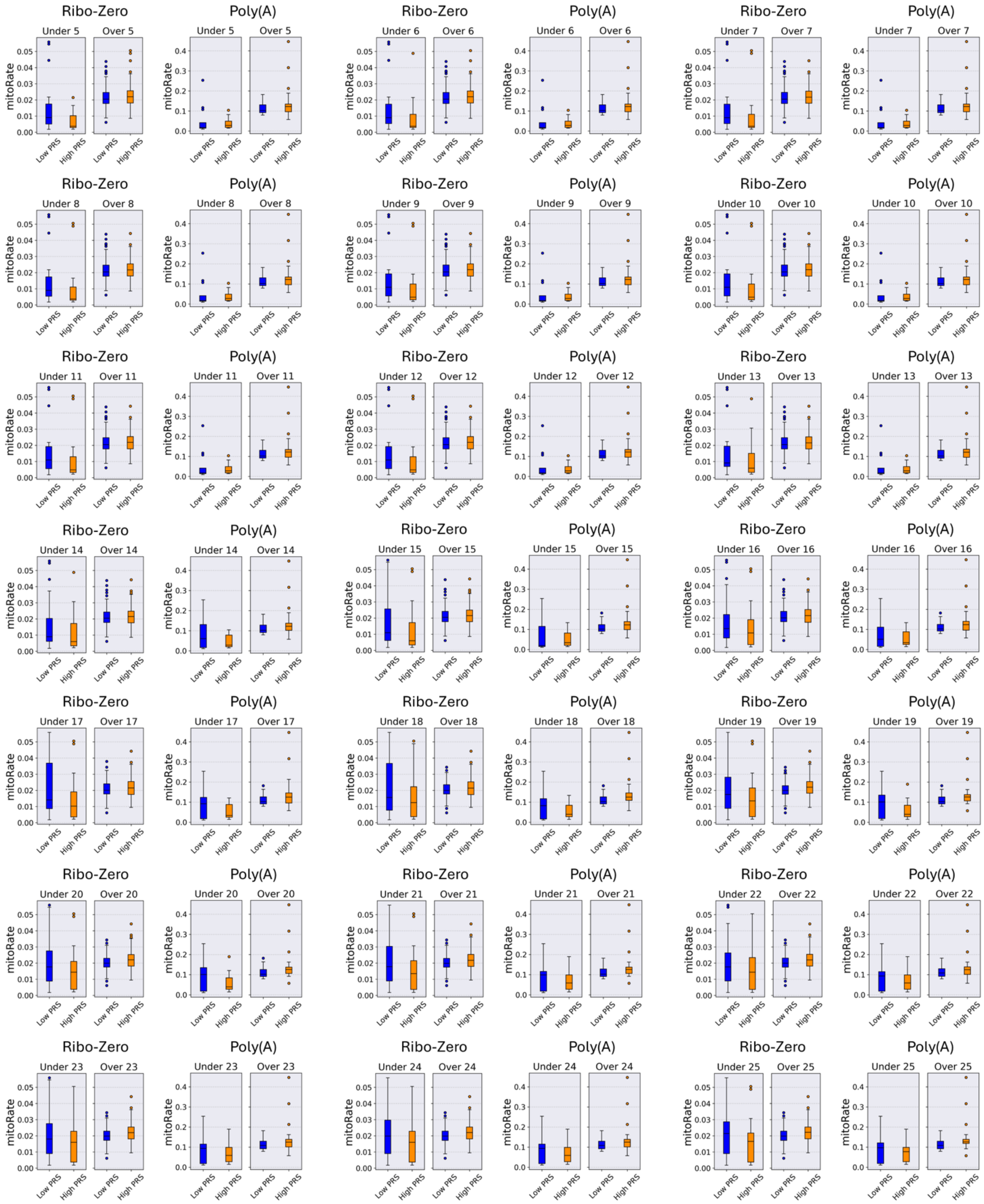


*Figure S41. Distribution of the decimal fraction of reads which mapped to the mitochondrial chromosome, of those which map at all (mitoRate), for low-PRS (blue) and high-PRS (orange) cohorts, grouped into the "Under" and "Over" age ranges for both Ribo-Zero and Poly(A) datasets. Results are displayed in separate panels for each age threshold Y in the 5—25 juvenile range; within each panel, data are organized by dataset and age group, and adjacent pairs of boxplots represent the corresponding low-PRS and high-PRS cohorts. Distributions are shown after correcting for significant differences in confounding factors between pairs of high-PRS and low-PRS cohorts originating from the same dataset and age group.*

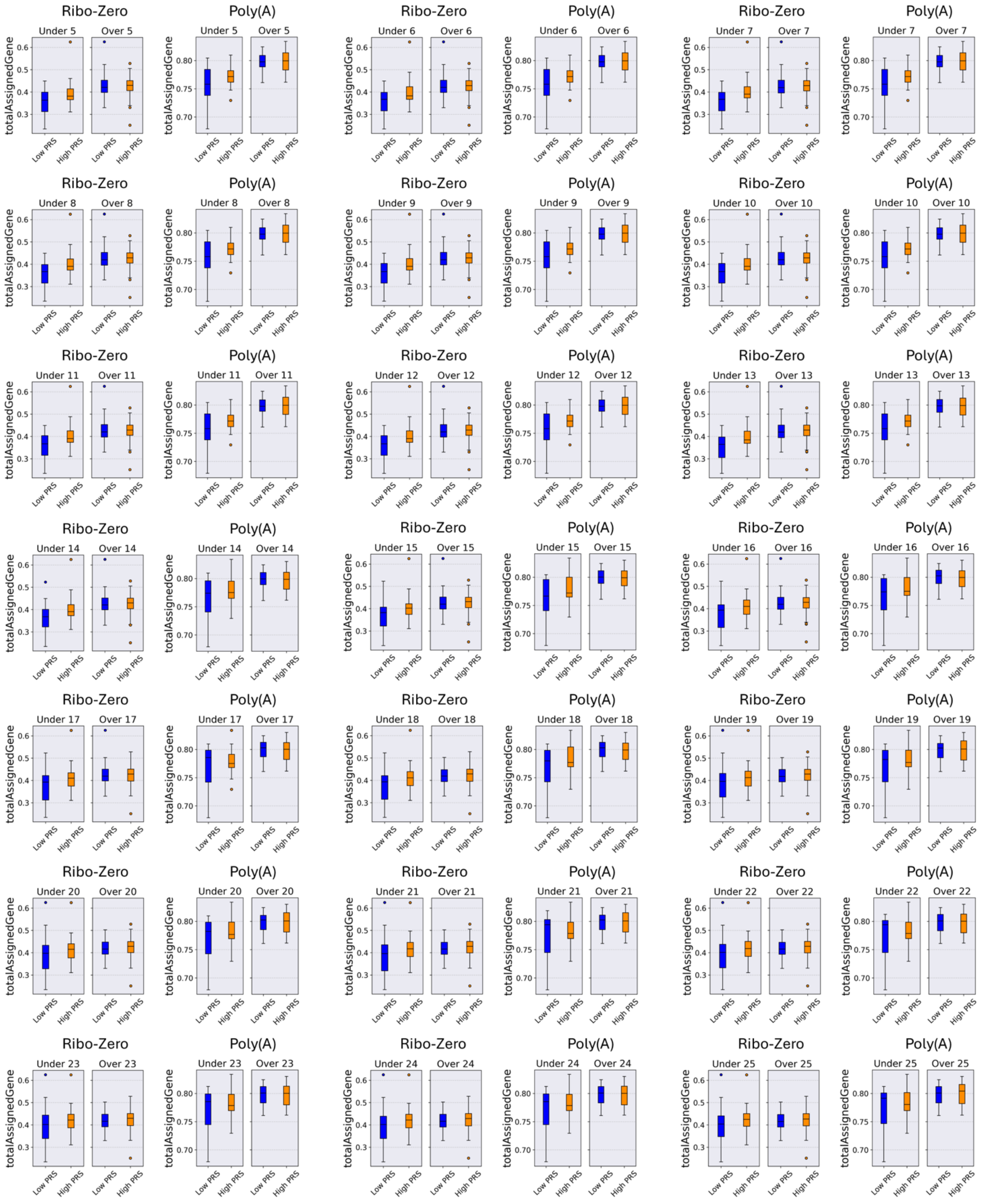


*Figure S42. Distribution of the decimal fraction of reads assigned unambiguously to a gene, with featureCounts of those in total (totalAssignedGene), for low-PRS (blue) and high-PRS (orange) cohorts, grouped into the "Under" and "Over" age ranges for both Ribo-Zero and Poly(A) datasets. Results are displayed in separate panels for each age threshold Y in the 5—25 juvenile range; within each panel, data are organized by dataset and age group, and adjacent pairs of boxplots represent the corresponding low-PRS and high-PRS cohorts. Distributions are shown after correcting for significant differences in confounding factors between pairs of high-PRS and low-PRS cohorts originating from the same dataset and age group.*

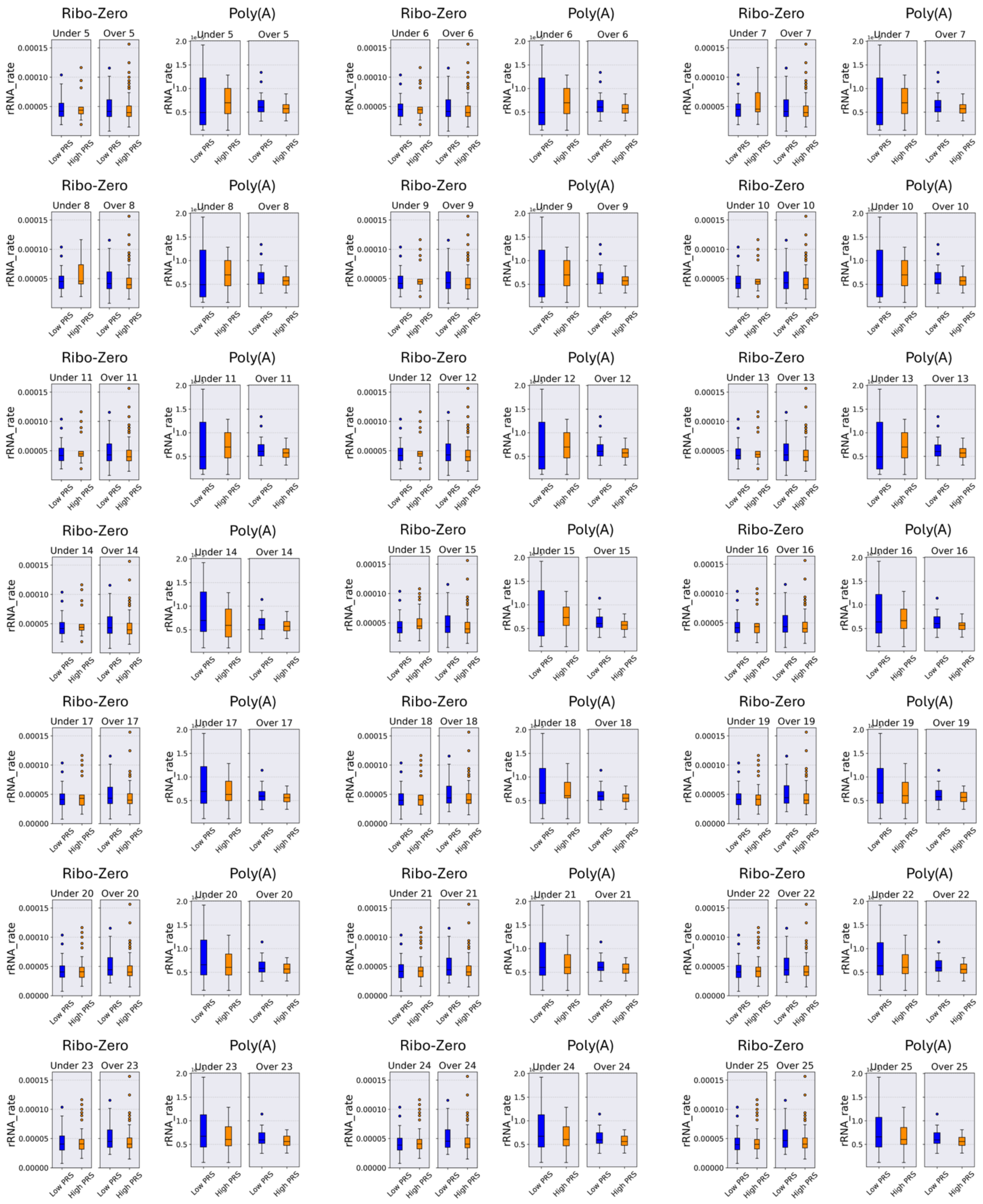


*Figure S43. Distribution of the decimal fraction of reads assigned to a gene whose type is 'rRNA', of those assigned to any gene (rRNA_rate), for low-PRS (blue) and high-PRS (orange) cohorts, grouped into the "Under" and "Over" age ranges for both Ribo-Zero and Poly(A) datasets. Results are displayed in separate panels for each age threshold Y in the 5—25 juvenile range; within each panel, data are organized by dataset and age group, and adjacent pairs of boxplots represent the corresponding low-PRS and high-PRS cohorts. Distributions are shown after correcting for significant differences in confounding factors between pairs of high-PRS and low-PRS cohorts originating from the same dataset and age group.*

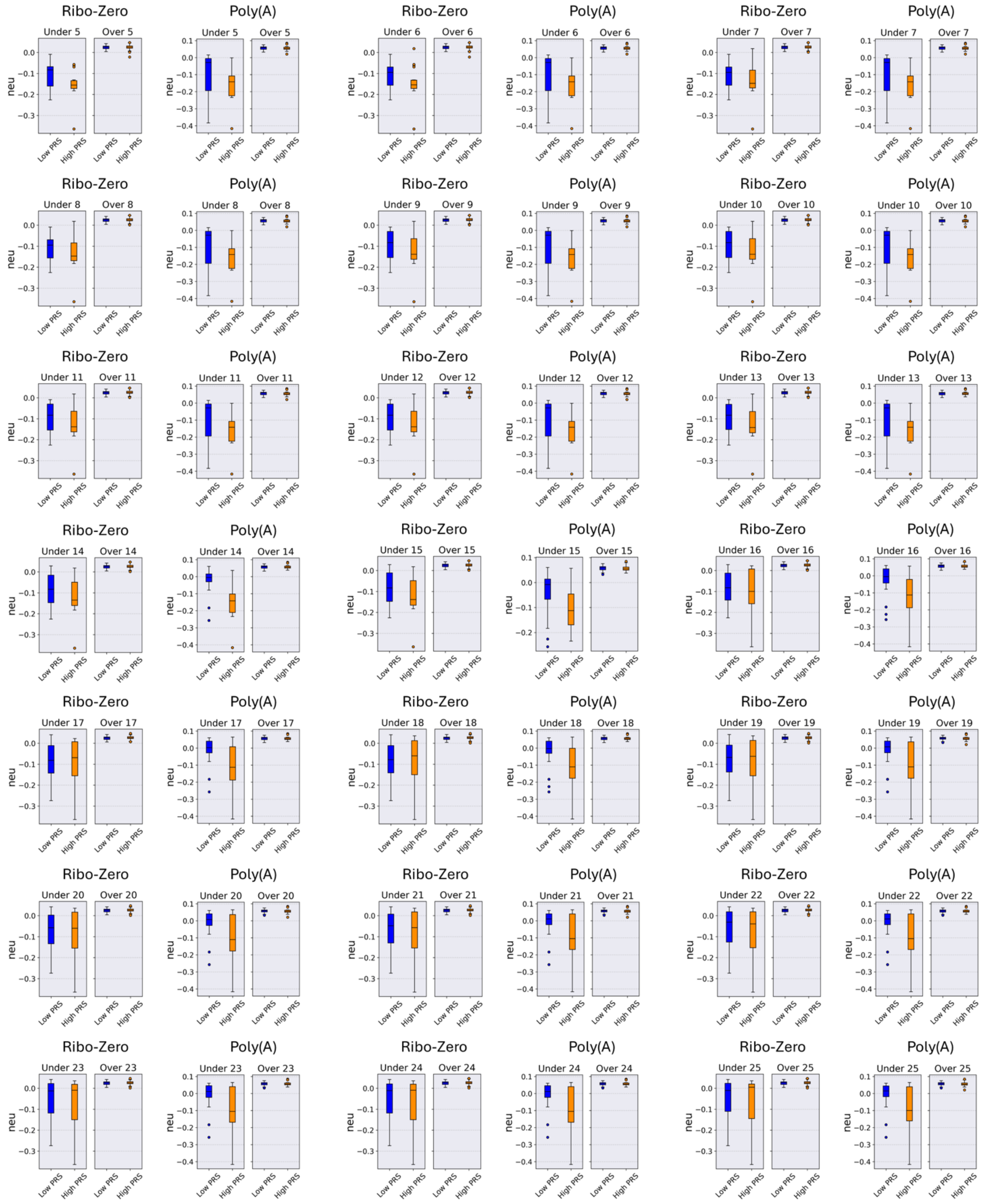


*Figure S44. Distribution of the estimated individual neuronal proportion (neu) for low-PRS (blue) and high-PRS (orange) cohorts, grouped into the "Under" and "Over" age ranges for both Ribo-Zero and Poly(A) datasets. Results are displayed in separate panels for each age threshold Y in the 5—25 juvenile range; within each panel, data are organized by dataset and age group, and adjacent pairs of boxplots represent the corresponding low-PRS and high-PRS cohorts. Distributions are shown after correcting for significant differences in confounding factors between pairs of high-PRS and low-PRS cohorts originating from the same dataset and age group.*
